# Information contracts determine what inverse perturbation benchmarks measure

**DOI:** 10.64898/2026.10.19.760271

**Authors:** Jianfu Li, L. Frank Huang, Wenyu Song, Ying Li, James W. Jakub, Hong Qin, Yan Luo, Cui Tao

## Abstract

Inverse perturbation benchmarks rank interventions toward a desired cellular state, but high accuracy can answer a different question from biological prioritization. We present InverseBench, which makes candidate information and endpoint construction explicit, and FSTN, a linear pipeline evaluated under it. With identical inputs, candidates and validation, nested ridge reaches Spearman 0.959 for a transcriptional principal-component proxy but 0.445 for independently measured K562 growth fitness across 7,354 genes. Controlled response-entry availability across seven tasks gives heterogeneous gains, 0.442 for CD4 activation and 0.116 for external fitness. Yet predefined shortlist comparisons show no primary-cutoff penalty in K562 or a melanoma co-culture screen, and secondary cutoffs reverse the K562 ordering. FSTN’s nonnegative set selector recovers 0.840 of measured two-gene constituents in three picks, against 0.771 for an unconstrained greedy. The framework separates proxy reconstruction, response-informed prioritization and unseen-intervention ranking: an endpoint-dependent accuracy gap does not by itself mean worse biological decisions.

---

Machine learning increasingly predicts cellular responses to perturbations [1,2,3,4,5,6]. Yet predicting a response and ranking interventions toward a desired state are different tasks. A ranking can exploit a measured response for every candidate, or must infer responses for interventions never performed in the source context. It can target a transcriptional summary constructed from the same assay, a predefined marker program, a surface protein or independently measured fitness. These choices determine what success means before model capacity enters the comparison.

Recent studies show that simple references can rival complex perturbation models [7,8,9,10]. An explanation based only on model classes remains incomplete when candidate coverage, information access and target construction differ. Regulatory and fate models [11,12,13,14,15], prescriptive approaches [16,17,18,19,20] and transport methods [21,22,23] operate under different inputs and objectives. Their comparison requires a task contract specifying available measurements, withheld interventions, endpoint provenance and training-only selection.

Those benchmarks largely score how well a model predicts a perturbation’s response. Ranking asks something narrower and more consequential: given a desired destination, which intervention should be tried first. The distinction is not academic. Adoptive tumour-infiltrating lymphocyte therapy improves progression-free survival over ipilimumab in advanced melanoma [24], yet a cell’s transcriptional phenotype alone does not establish that it is tumour-reactive [25], so a ranking scored against a transcriptional summary can succeed on its own terms while selecting the wrong interventions. Two things then decide the answer before any model is chosen. The first is what the ranker is allowed to know, which ranges from a measured response for every candidate to no candidate-specific measurement at all. The second is where the destination comes from, since a target distilled from the same assay that supplies the features is not the same object as an independently measured phenotype. Neither is a nuisance parameter to be averaged over. Both change what a high score means, so unless each is stated a score cannot be placed against another obtained under different access or a different destination.

InverseBench is the benchmark we propose for this task, and FSTN is the reference pipeline we propose to solve it: a shared state embedding, a response model whose continuous-target ridge supplies the ranking score, and a nonnegative set selector for combinations. Throughout the comparisons we name estimators by what they do rather than by the pipeline they belong to, because the contrasts are between objectives and information budgets, not brands. InverseBench makes that contract experimentally testable. We vary candidate-response entries while holding candidates, outcomes, learners and validation fixed. We compare a transcriptional proxy with an independent functional endpoint on exactly the same interventions and source inputs. Finally, we distinguish sensitivity to the endpoint or training objective from numerical variation across computational settings. The contribution is an evaluation framework with a consequential diagnostic result: near-perfect reconstruction of a dominant transcriptional axis can coexist with substantially lower ranking of an independently measured biological outcome. Here we show that this gap survives removing model selection, that the value of measured candidate responses is large on some tasks and indistinguishable from zero on others, and that a large difference in endpoint predictability need not translate into a worse intervention shortlist. We also report where the framework’s own supporting theory fails, since a diagnostic that cannot predict which real task will reward learning should be presented as a negative result rather than a tool.

## Results

### An information contract separates three ranking problems

Track A ranks held-out candidates supplied with their measured source-context response. Track B ranks candidates without their own measured response, using side information such as a gene’s expression across separate anchor interventions. Both differ from inferring a successful cellular conversion or a minimal intervention combination. Holding out candidate outcomes does not by itself make a task an unseen-intervention prediction problem (Fig. 1a).

**Figure 1.**
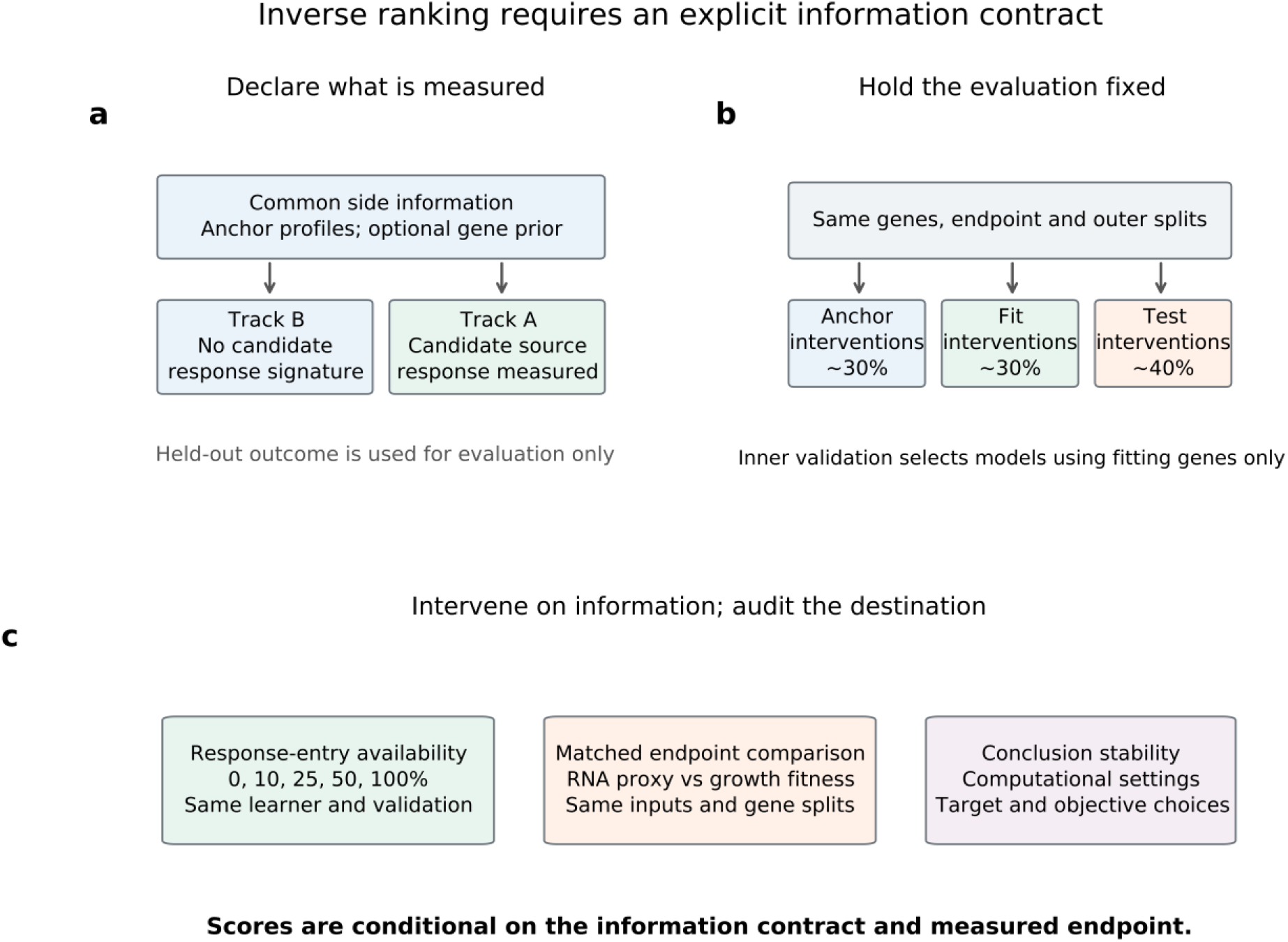
Information contracts for inverse intervention ranking. a, Track B uses common anchor-response profiles and optional gene priors; Track A additionally receives a measured candidate-source response. b, Identical outer partitions separate anchor, fitting and test genes. Inner validation uses fitting genes only. c, Controlled response-entry availability, a matched endpoint comparison and conclusion-stability checks test distinct sources of apparent success. These tasks do not measure experimental cell conversion or minimum intervention sets.

We constructed seven tasks spanning six study groups (Table 1). The two Frangieh conditions share a study and are not biological replications. Four tasks retain established cross-condition axes; three add fixed biological endpoints: activated CD69/IL2RA expression, surface PD-L1 protein and external K562 growth fitness [26,27,28]. These are review-motivated exploratory extensions. Endpoints and eligibility were fixed before their respective model fits; the full study was not prospectively preregistered. Song data had previously been examined with an unsupervised endpoint. Neither multiple splits nor additional endpoints create independent biological cohorts.

**Table 1.**
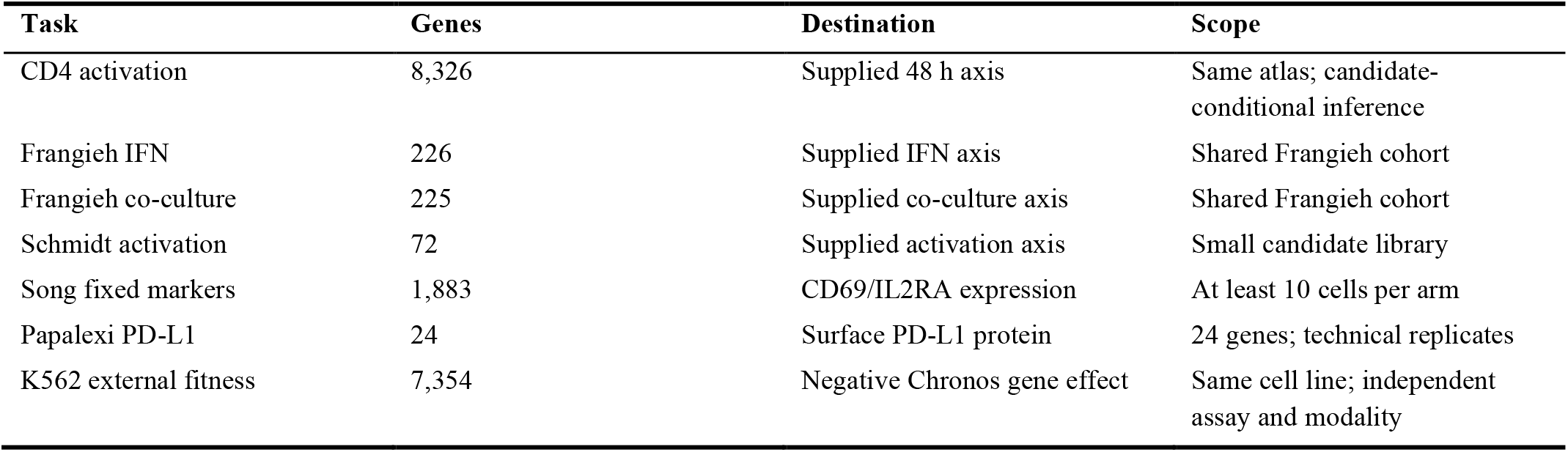
Tasks, eligible gene interventions and endpoint provenance. Seven tasks represent six study groups. Counts are after shared candidate eligibility, not independent biological sample counts.

Each outer split reserves approximately 30% of eligible genes as anchors, 30% for fitting and 40% for testing. Track B profiles contain expression of a candidate gene across anchor interventions only. Fitting and test candidates therefore have the same information structure; neither contributes its own intervention row to that profile. All models share eligible candidates, outcomes and outer partitions. Inner validation uses fitting genes only (Fig. 1b).

### Controlled availability reveals a heterogeneous value of source responses

For a controlled comparison, we add outcome-independently masked entries from each candidate’s source response to its common anchor profile. Nested masks reveal 0, 10, 25, 50 or 100% of selected response entries; their indicators disclose which entries are missing. Candidate populations, outcomes, ridge selection rules and outer splits remain fixed. This intervention measures the value of response entries under a specified pipeline, rather than the number or cost of experiments (Fig. 2).

**Figure 2.**
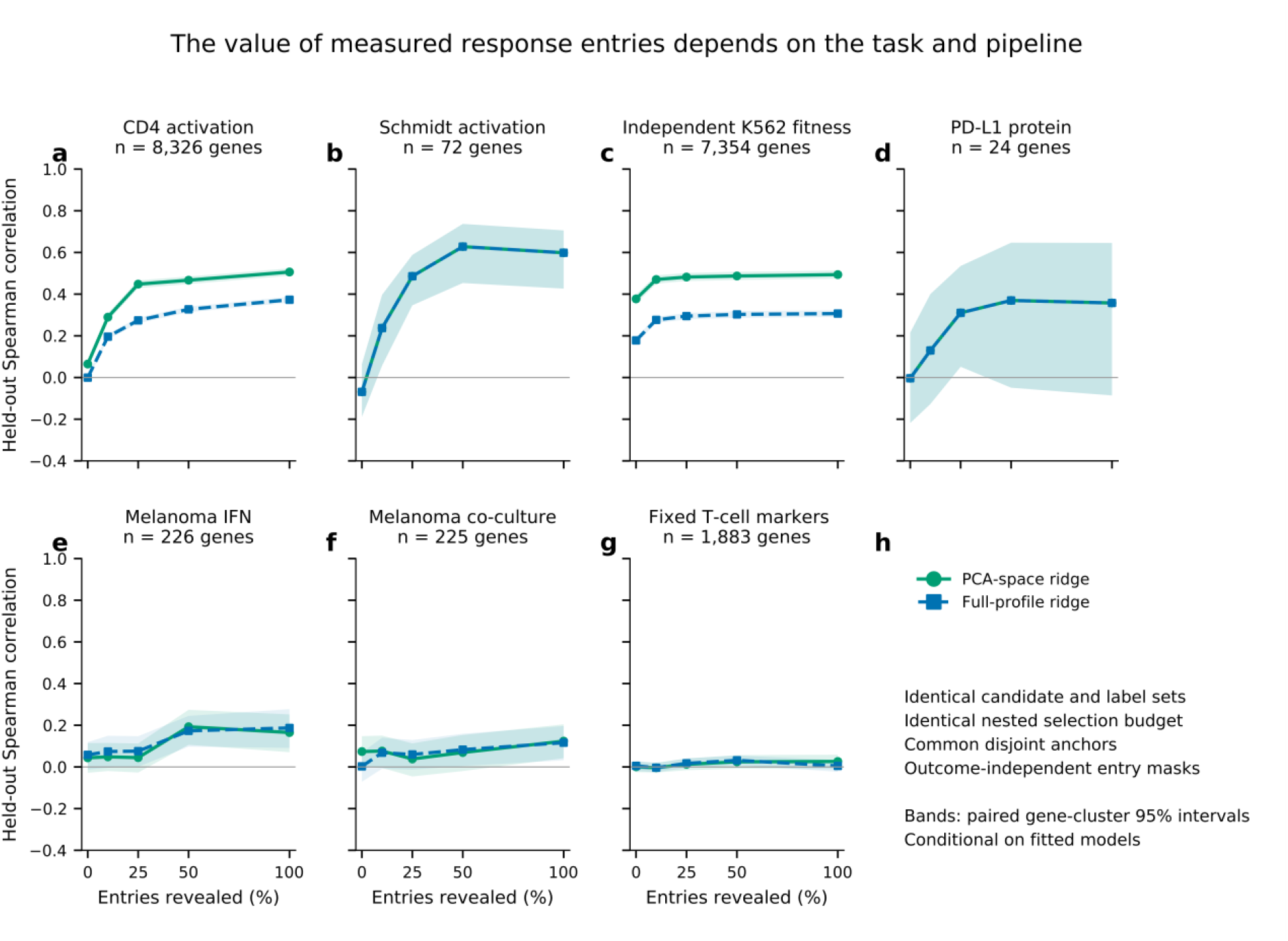
Controlled response-entry availability across seven tasks. a-g, Mean held-out Spearman correlation for CD4 activation, Schmidt activation, external K562 fitness, surface PD-L1, Frangieh IFN, Frangieh co-culture and fixed Song T-cell markers. Each fraction reveals a random nested subset of source-response entries for every candidate, with common anchor profiles retained. Solid curves use inner-fold PCA; dashed curves retain full profiles. h, Common controls and interval definition. Shading denotes paired gene-cluster 95% intervals conditional on fitted models, not donor-level uncertainty. The Frangieh tasks share a cohort. Fractions do not measure intervention count or experimental cost.

In the fixed-dimension pipeline, revealing all selected entries raises mean held-out Spearman correlation by 0.442 for CD4 activation, with a paired gene-cluster 95% interval of [0.419, 0.465]. External K562 fitness gains 0.116 [0.101, 0.133]. Schmidt activation also shows a positive conditional contrast. These effects demonstrate useful information in measured source responses under controlled estimation; they are not model-independent bounds on unseen-intervention performance (Table 2).

**Table 2.**
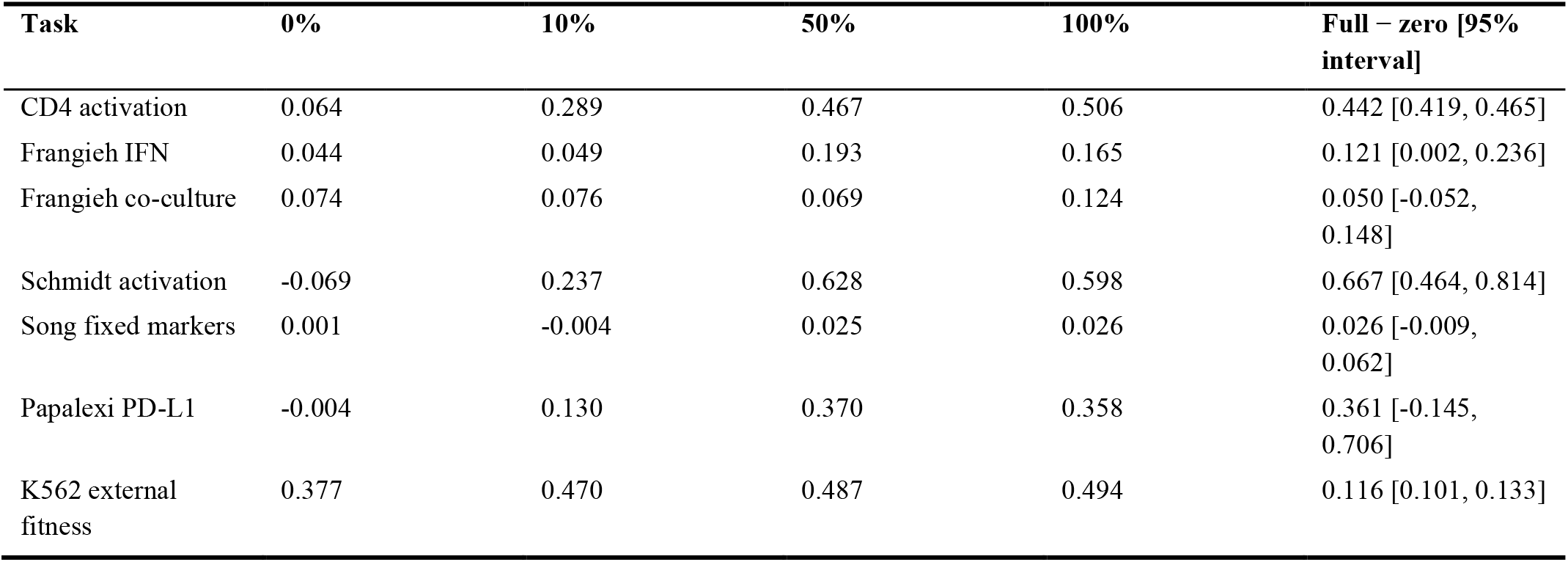
Controlled response-entry availability under the fixed-dimension ridge pipeline. Entries are mean test Spearman correlations over ten outer splits. The final column is the paired full-minus-zero availability effect with a gene-cluster 95% interval, conditional on fixed fitted models. Intervals are exploratory and unadjusted for multiple comparisons. All tasks, including weak and uncertain contrasts, are retained.

The added biological endpoints prevent interpreting these gains as universal. Song’s fixed CD69/IL2RA endpoint produces a gain of 0.026 [−0.009, 0.062]. The primary PD-L1 protein comparison has a gain of 0.361 [−0.145, 0.706], with only 24 eligible interventions. Its secondary top-decile AUROC contrast is positive, but cannot replace the uncertain primary correlation result. Performance need not improve monotonically with availability: extra features can impair finite-sample estimation. Removing PCA preserves positive full-minus-zero response-entry effects for CD4 activation (0.374 [0.359, 0.389]) and external fitness (0.129 [0.117, 0.140]), while the Song contrast remains near zero (Supplementary Note 10).

Eligibility for the two weaker endpoints explains much of their uncertainty and is therefore stated here rather than deferred. The protein destination is lower surface PD-L1 in replicate 2 of a multimodal screen, measured as negative mean log1p ADT abundance against non-targeting controls; direct CD274 perturbation and its RNA readout are excluded, and requiring at least 20 cells in each arm leaves 24 gene interventions. The marker destination is mean activated CD69 and IL2RA expression, with both readouts and their direct interventions excluded from features and candidates, requiring at least ten cells per arm and leaving 1,883 genes. Neither count is a guide count. The marker cohort had already been analysed with an unsupervised axis, so it is not a prospectively untouched validation set, and its weak contrast is retained rather than attributed to a noise ceiling.

### A matched proxy comparison changes the interpretation of accuracy

To isolate endpoint construction, we paired K562 CRISPRi transcriptomic signatures [29] with a separately measured pooled-CRISPR knockout growth endpoint from DepMap 24Q4 [30]. The destination was fixed as stronger growth suppression, represented by negative Chronos gene effect. The verified K562 model is ACH-000551. This adds an independent assay and intervention modality, rather than a new cell type or clinical outcome (Supplementary Note 11).

Both tasks use exactly 7,354 eligible genes, source inputs, outer partitions and nested ridge selection. A fixed readout-gene partition separates source features from the genes defining a PC1 proxy. The proxy axis is fitted on anchor interventions only; neither task uses held-out outcomes to select models. Mean test Spearman correlation is 0.959 [0.956, 0.962] for the proxy and 0.445 [0.425, 0.466] for external fitness (Fig. 3a). Corresponding mean AUROCs are 0.999 and 0.863.

**Figure 3.**
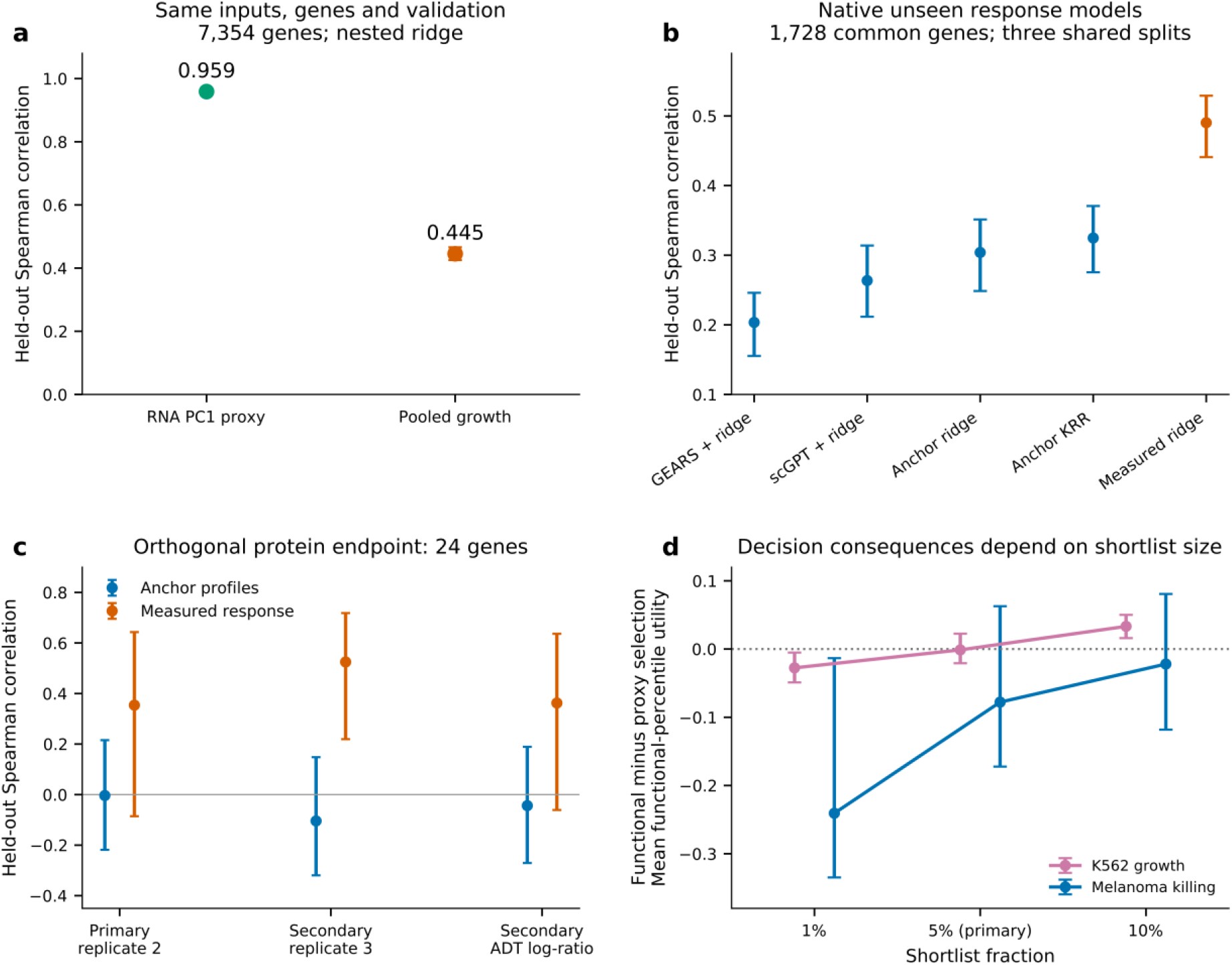
Biological endpoints and stronger unseen comparators. a, Same-input, same-candidate comparison of anchor-fitted RNA PC1 and independent pooled-growth fitness for 7,354 K562 genes, with nested ridge selection. b, Native GEARS and scGPT functional readouts versus anchor-profile ridge/kernel ridge and measured-response ridge on 1,728 common genes from the raw K562 essential-gene assay, across three shared splits. Response models train on anchor interventions only; functional heads use fitting outcomes only. Native recipes differ in readout coverage and inference averaging. This candidate population differs from the 7,354-gene comparison in a. c, Primary PD-L1 and secondary technical/normalization checks on 24 genes, rescoring the same predictions without refitting. d, Functional-minus-proxy selection differences in mean functional percentile for 1%, 5% (primary) and 10% shortlists in K562 growth and melanoma/TIL killing. Five families are selected by inner validation for each endpoint. Neither primary interval establishes a penalty from proxy selection; K562 ordering reverses across secondary cutoffs. Secondary intervals are exploratory and unadjusted for multiple comparisons. Error bars in a–d denote conditional gene-cluster 95% intervals; technical replicate results are not independent biological replication.

The gap is a property of the endpoint rather than of the selected estimator. A covariance score fitted on the same anchors reaches 0.928 [0.922, 0.933] on the proxy and 0.403 [0.381, 0.424] on fitness, and a fixed ridge penalty gives 0.959 and 0.444, within 0.001 of the selected values, so the ordering survives removing model selection entirely. The functional endpoint is the negative Chronos gene effect from DepMap 24Q4, pinned by DOI, with K562 identified as model ACH-000551; higher values mean stronger loss of pooled growth fitness. Eligibility follows from named intervention and readout overlap, a finite gene effect and unique-gene grouping, with repeated transcriptomic records averaged per gene and no model or outcome threshold applied, which is what leaves exactly 7,354 candidates. Growth suppression is not therapeutic selectivity, and neither endpoint was chosen by comparing held-out candidate performance.

Thus a nearly perfect transcriptomic ranking does not establish nearly perfect functional prioritization, even with identical candidate and validation populations. This comparison does not identify a biological ceiling: assay noise, intervention modality and endpoint content all change. Its contribution is to expose a conclusion that a high proxy score alone cannot support. Near-ceiling principal-component tasks are retained as reconstruction controls rather than counted as independent evidence of difficult biological ranking.

We next tested whether this endpoint discrepancy changes a practical intervention shortlist. Five learner families were tuned separately for each endpoint, and the family was selected by inner validation only. For the predefined top-5% shortlist, mean functional percentile was 0.841 for proxy-selected models and 0.839 for functional-selected models. The paired functional-minus-proxy difference was −0.001 [−0.021, 0.023]. Top-functional-decile precision at the same cutoff is 0.501 for both selections, so the two shortlists agree on a second, rank-free measure as well. Thus the large predictability difference did not produce a detectable functional shortlist penalty in this system, although absence of a detected difference does not establish equivalence. The prespecified secondary cutoffs reverse the ordering: functional-minus-proxy utility is −0.028 [−0.049, −0.005] at 1% and 0.033 [0.016, 0.050] at 10% (Fig. 3d). These exploratory, unadjusted comparisons show cutoff-dependent consequences rather than a uniform downstream penalty (Supplementary Note 11).

A separate melanoma/TIL co-culture viability assay provides a second functional context on 226 shared genes. The endpoint averages depletion against IFNγ-treated controls across all three reported TIL ratios, fixed before fitting. Predictability again separates sharply from utility: inner-selected models reach mean test correlation 0.913 for the RNA proxy but −0.057 for the functional endpoint, so the functional outcome is essentially unpredictable here. Despite that, at the top-5% cutoff mean functional percentile is 0.518 for proxy selection and 0.440 for functional selection, a difference of −0.078 [−0.172, 0.063], again providing no evidence that proxy selection worsens the functional shortlist. Training directly on a functional endpoint therefore does not guarantee a better shortlist when that endpoint is itself poorly predicted. With five genes per split in the test shortlist, this context constrains interpretation rather than settling it. This assay shares the previously analyzed Frangieh cohort and does not constitute a new independent source study (Supplementary Note 11).

The small multimodal protein screen supplies a complementary, explicitly limited check. Using the same predictions trained for primary replicate-2 PD-L1 outcomes, rescoring on technical replicate 3 yields response-informed ridge correlation 0.525 [0.219, 0.718]. The primary replicate-2 estimate is 0.354 [−0.086, 0.643]; a within-cell ADT normalization sensitivity remains uncertain (Fig. 3c). These technical checks characterize assay sensitivity and do not provide additional biological replication.

### Stronger unseen comparators test the scope of the information effect

Track B is evaluated with ridge, radial-basis kernel ridge, histogram gradient boosting, an MLP and an inductive graph-neighbour model. Each family receives six candidate specifications and the same three-fold validation objective. A frozen Geneformer token embedding [31] provides an external gene prior, combined with the common anchor profiles. It is a token-prior baseline, not native foundation-model response inference. Unknown tokens are retained with a missingness indicator, preserving candidate coverage. Pretraining overlap with evaluated cohorts is unverified.

Full-profile controls retain every nonconstant anchor coordinate instead of reducing profiles to 30 principal components. They distinguish the consequences of unavailable candidate responses from an imposed representation bottleneck (Supplementary Note 10). On external fitness, unseen kernel ridge reaches correlation 0.383 [0.363, 0.404], exceeding response-informed ridge at 0.373 in this representation; validation-selected response-informed boosting reaches 0.445 [0.426, 0.463]. Thus comparator choice can reverse a simple comparison between tracks. The selection protocol that makes this comparison fair was fixed before any model fit and is given in the Methods; in particular the graph comparator builds messages from fitting nodes only, so test-node features cannot alter fitted representations.

We additionally trained native GEARS and pretrained scGPT response models on raw K562 essential-gene CRISPRi cells, with 1,728 common candidates and three shared splits (Fig. 3b). Only anchor interventions and controls train or validate response prediction; both fitting and test interventions remain unseen. A common nested ridge head learns growth fitness from fitting genes. Mean test correlations are 0.203 for GEARS, 0.264 for scGPT, 0.304 for anchor-profile ridge and 0.325 for anchor-profile kernel ridge; measured-response ridge reaches 0.490. However, scGPT improves primary top-5% shortlist utility over anchor ridge by 0.076 [0.001, 0.132], despite a lower point estimate of overall correlation. This conditional, exploratory result further separates global ranking accuracy from shortlist utility. Candidate coverage and head tuning are matched; native architecture, readout coverage and inference budgets differ, and three splits do not establish a universal model ordering (Supplementary Note 11).

### The nonnegative set selector recovers measured constituents

Ranking single interventions and choosing a combination are different operations, so we evaluated FSTN’s set selector on its own terms. Under measured single-perturbation effects, 131 two-gene Norman targets were reconstructed from a 105-gene single library, and the selector is scored on whether it recovers the two constituents of the measured combination. Nonnegative orthogonal matching pursuit with a joint refit after every pick (Algorithm 1, Methods) recovers 0.725, 0.840 and 0.916 of the constituent pair within two, three and five picks, against 0.714, 0.771 and 0.855 for an unconstrained greedy residual and 0.546, 0.615 and 0.706 for independent cosine ranking. Paired per-target bootstrap intervals put the advantage over the unconstrained selector at +0.069 [+0.038, +0.099] at three picks and +0.061 [+0.034, +0.092] at five, while at two picks it is +0.012 [+0.000, +0.027] and not distinguishable from zero. The nonnegativity constraint therefore earns its place at larger budgets rather than on the headline pair. This is measured constituent-pair retrieval, not minimal intervention design: the generating pair is an answer key for identity, not proof that no other combination reaches the same phenotype, and 129 of the 131 targets share a constituent gene with another target, so the intervals are optimistic (Supplementary Note 3).

### Conclusion stability differs from numerical identity

The reference grid these checks are run on is the seven-column, eight-model-class benchmark retained in Supplementary Note 10. Its pattern motivates the information contract. On the five single-condition principal-component columns the projection reference spans 0.953 to 1.000 and matches or beats the continuous-target ridge on every one, so those columns cannot show learning helping and are retained as reconstruction controls. They still separate the weaker generative and disentangled-latent methods, which fall as low as 0.724. Only the cross-condition columns can show learning helping. On the grid’s seven columns FSTN’s linear ridge significantly exceeds neural transport in six and the disentangled-latent method in six, with one significant deficit, on Frangieh control-to-IFN (−0.122 [−0.195, −0.046]); widening to all twelve systems that carry paired candidate-cluster intervals gives nine and eight significant exceedances respectively, and that same single deficit. That deficit tracks the objective, not the model class: the continuous-target ridge reaches 0.639 there against CellOT’s 0.761, while rank-transforming the training outcome to match the top-decile metric reaches 0.766, a difference from CellOT of [−0.022, 0.029] that establishes neither superiority nor equivalence. A grid that looked like a model-class comparison is therefore largely a target-construction and objective comparison.

Across the seven reference configurations, the signs of mean Ridge-minus-CellOT, Ridge-minus-PAIRING and kernel-ridge-minus-Ridge AUROC differences are unchanged across four computational settings (Fig. 4a–c). These settings include the reference computation, newer and frozen environments, and preservation of the reference PCA input layout. These settings are not spot checks: all seventy shared-space splits and sixty native-tool evaluations were rerun, and twenty independent repeat splits reproduce the current score arrays exactly under the recorded settings, and the unresolved discrepancy is in the neural arms. Preserving the historical principal-component input layout reduces the largest mean AUROC discrepancy from 0.0191 to 0.0138, so input layout accounts for part of it and most of it remains unexplained. Exact historical neural score recovery remains unresolved; sign stability does not establish practical superiority when differences are small.

**Figure 4.**
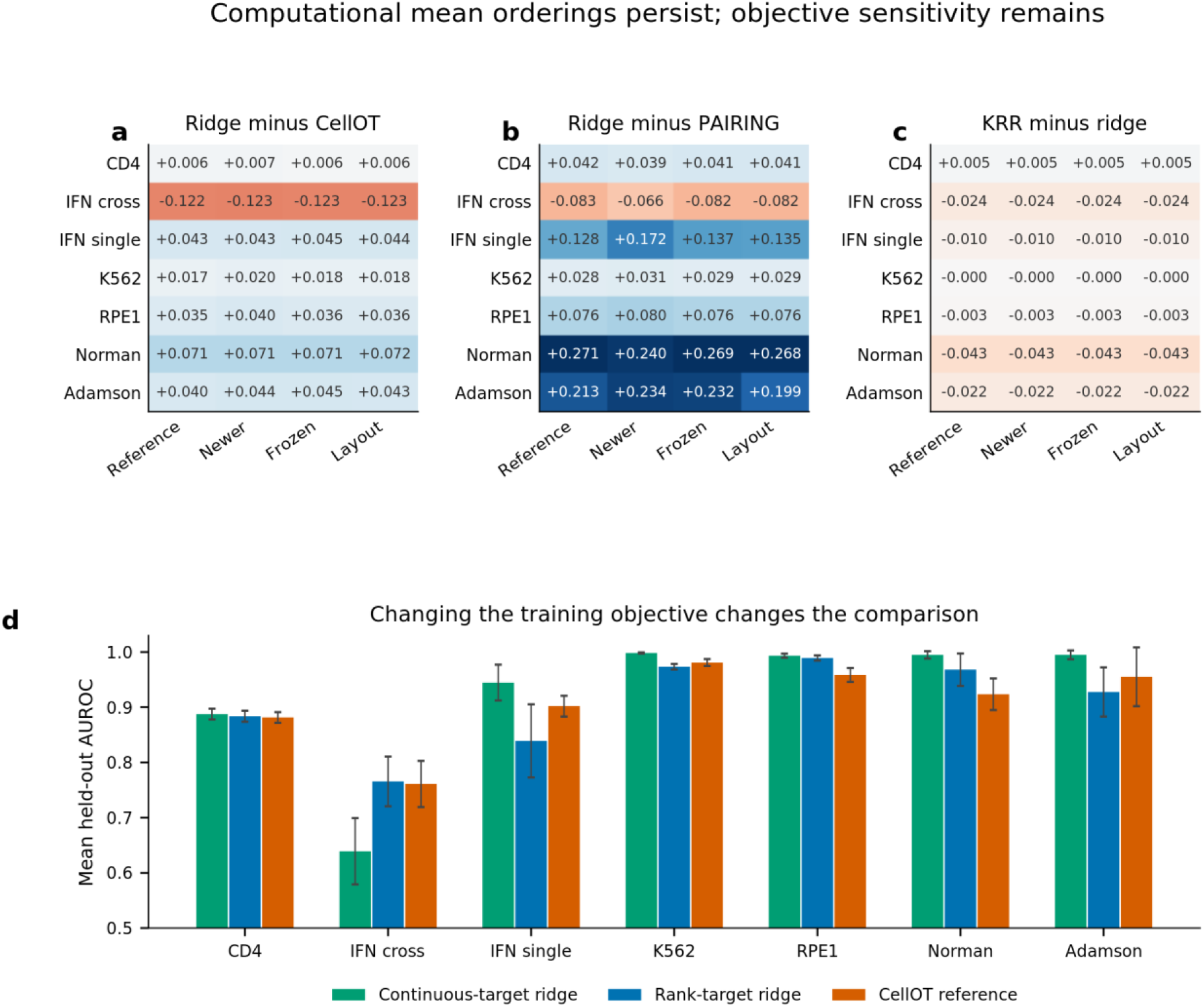
Numerical stability and objective sensitivity. a-c, Mean AUROC differences for Ridge versus CellOT, Ridge versus PAIRING and kernel ridge versus Ridge across four computations of seven reference configurations. No sign reversal occurs in these contrasts; small differences do not establish practical superiority. d, Continuous-target ridge, rank-target ridge and CellOT on the same reference systems. Bars show means; error bars where shown are s.d. across ten overlapping splits, not confidence intervals. Changing the training objective alters the Frangieh comparison, whereas computational variation preserves the displayed mean orderings. The near-equal rank-ridge and CellOT means do not establish equivalence.

Training objectives can change conclusions more than these numerical differences (Fig. 4d), and the rank objective is not uniformly better elsewhere. Actual-target PC2 retraining likewise differs sharply from merely rescoring PC1-trained predictions against PC2 (Supplementary Note 7). Numerical reproducibility and scientific robustness therefore require separate reporting.

A Gaussian eigenvector condition characterizes when population covariance projection and optimal linear ranking coincide (Supplementary Note 2). Training-only diagnostics track fitted gains in controlled simulation: across 4,320 configurations spanning training sizes, dimensions, anisotropies and signal-to-noise ratios, predicted and observed gains correlate 0.846 with mean absolute error 0.041 under a Gaussian population, and 0.815 with error 0.055 under a Student-t(3) population for which the condition is not assumed to hold. That calibration does not survive contact with the real studies: the study-level correlation is −0.143 across six groups, with a bootstrap interval spanning −1 to 1, and leave-study-out prediction has mean absolute error 0.093. The failures are not uniform shrinkage but sign errors. Schmidt has the largest predicted gain of the seven tasks at 0.335 and the most negative observed gain at −0.102, and the protein task is predicted at 0.063 against an observed −0.042. We retain this negative calibration result and treat the theory as conditional supporting analysis, rather than a predictor of real biological task difficulty (Supplementary Note 12).

## Discussion

An information contract is useful when it changes an inference, not merely a reporting checklist. Here it separates a near-perfect transcriptional reconstruction result from a much weaker external functional ranking on the same genes and inputs. Controlled response-entry experiments then locate useful source-response information under specified learners, while negative marker and uncertain protein results delimit generalization. These outcomes support a benchmark-validity claim rather than a claim that one model family solves inverse intervention design.

Independent evidence remains finite, and the specific shape of that limit matters more than the general caveat. Seven tasks span six study groups, because the two Frangieh conditions share a cohort; paired endpoints, technical replicates and repeated partitions multiply measurements without adding biological contexts. The two added functional contexts are both assay extensions of cohorts already analysed here, not independent source studies. Conditional gene-cluster intervals preserve repeated-candidate dependence but do not estimate donor-level transfer, carry no refitting uncertainty and are unadjusted across tasks and cutoffs.

Three comparisons are confounded in ways no interval expresses. The external fitness contrast crosses both an assay boundary, CRISPRi transcriptional against CRISPR-knockout growth, and an intervention modality, so assay noise, modality and endpoint content all change together and the 0.959 against 0.445 gap cannot be read as a biological ceiling. The protein experiment rests on 24 interventions, and its primary interval includes zero; a positive secondary contrast on a technical replicate does not resolve that, since replicate outcomes correlate 0.617 and 0.768 with the primary and so supply no independent biological replication. The marker cohort permits as few as ten cells per arm and had already been examined with an unsupervised axis.

The supporting theory is reported as a negative calibration rather than a tool. It predicts simulated gains well and real-study gains not at all, with sign errors on the two tasks where it is most confident, so it explains one population comparison and should not be used to anticipate which real task will reward learning. Equally, the unseen-perturbation frontier is bounded by what was tested: five learner families, six settings each and a frozen token prior whose pretraining overlap with the evaluated cohorts is unverified. No conclusion here requires every unseen-perturbation algorithm to underperform, and no universal information-theoretic limit follows from these comparisons.

The melanoma application remains an illustration: its coordinate is an expression-derived program score, not a trajectory or an independent functional validation, and its supporting screens do not establish tumour specificity or successful reprogramming (Supplementary Note 1).

What follows for anyone building or reading such a benchmark is concrete. A reported score should name three things: which candidate responses the ranker could see, where the destination came from and whether it was measured by the same assay as the features, and what was permitted to touch held-out outcomes. Absent those, two numbers from different papers are not comparable, and a near-ceiling value may index reconstruction of a dominant axis rather than useful prioritization. The corollary for interpretation is that a benchmark can be internally valid and still answer a question other than the one a reader cares about, which is why we report the proxy and functional endpoints side by side rather than selecting whichever supports a stronger claim.

Future benchmarks can extend the same contract to prospective destinations, donor-held-out evaluation and measured combinations. For current datasets, the actionable conclusion is narrower: state exactly which candidate responses and endpoint definitions a score uses, and test whether the biological inference survives changing them. A high inverse-ranking score is evidence for its declared task, not an interchangeable measure of intervention utility.

## Methods

Task construction and provenance. Cross-condition source signatures and supplied target axes were retained for CD4 activation, Frangieh IFN/co-culture and Schmidt activation [32]. Eligible named genes must have both a source intervention signature and a source readout, enabling identical candidate populations in both tracks. Song’s destination is the mean activated CD69 and IL2RA control-referenced expression; those readouts and their direct interventions are excluded. Papalexi source features are replicate-1 RNA signatures, and the primary outcome is negative replicate-2 mean log1p surface PD-L1 relative to non-targeting controls. Barcode order and guide identities were verified; direct CD274 intervention and its RNA readout were excluded. At least 20 cells per required protein-screen arm and 10 per Song arm were required. External K562 outcomes are negative DepMap 24Q4 Chronos gene effects, pinned to dataset DOI 10.25452/figshare.plus.27993248.v1. Source signatures with repeated eligible gene identities were averaged before splitting; no outcome-guided eligibility was used. Accessions, hashes and cohort dependence are documented in Supplementary Notes 10–12 and Supplementary Table 1.

Information and feature fitting. Ten seeded 60/40 outer partitions divide training genes into disjoint anchor and fitting halves. Co-response profiles use only anchor intervention rows, never fitting/test interventions. Source readouts are selected by anchor-only variance (up to 512), and anchor means and variances standardize response entries. Seeded nested entry masks are independent of outcomes. Zero availability removes all candidate-source values; explicit missingness indicators accompany revealed entries. Common profile features remain available at every fraction. Inner-fold StandardScaler and PCA use fitting-fold candidates only, with at most 30 components; final transformations are refitted on all fitting candidates. Full-profile controls remove PCA and drop only constant coordinates. Candidate responses are intentionally supplied in Track A and are withheld in Track B; test outcomes are never model inputs.

Matched learners and selection. For each of the three information arms, the five equally tuned families compete on inner cross-validation alone, the winning family is then scored once on outer test, and no outer-test result is permitted to influence any choice. Test results for every family are reported rather than only the selected one, so a reader can see the spread the selection was drawn from. The external-prior arms are reported separately rather than folded into the frontier, since a token prior is a different kind of input from an anchor profile. A constant prediction is scored as zero correlation rather than discarded, which prevents a degenerate model from being silently excluded. Three-fold inner validation maximizes Spearman correlation. Ridge penalties are 0.1, 10 and 100; RBF kernel ridge uses three training-distance-based bandwidths and penalty 1. Histogram boosting uses 150 iterations and 7, 15 or 31 leaves. MLP and inductive graph-neighbour models use 16, 32 or 64 hidden units, 200 epochs and Adam learning rate 0.001 with weight decay 0.001. Every family crosses its three settings with raw or rank-transformed training targets, giving six settings per arm. Target scaling and ranks use training rows only. Model-family selection, where reported, uses inner scores across the same five core families. The graph model uses up to ten fitting neighbours and a mean-neighbour aggregation; test queries cannot alter fitted-node features. Geneformer V1-10M word embeddings are frozen, matched to the checkpoint vocabulary and never fine-tuned. Missing genes remain eligible. Full architecture, token coverage and hyperparameters are in Supplementary Note 10

### Algorithm 1

Response ranking and nonnegative set selection.

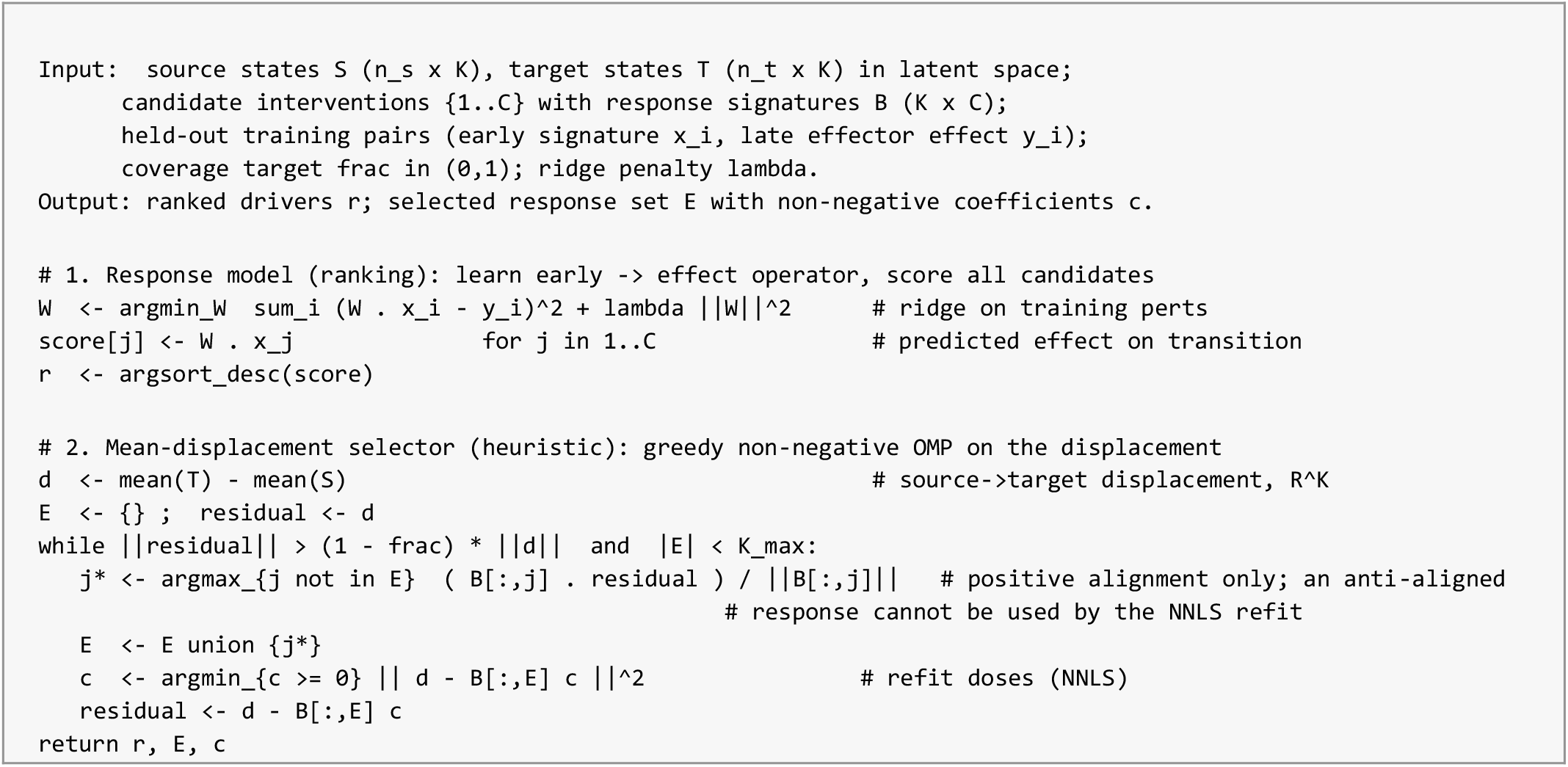

Endpoints and statistics. Every interval reported here is a 1,000-draw paired gene-cluster bootstrap that resamples candidate genes with their shared multiplicities across repeated test appearances, then rebuilds rankings and shortlists within each resample. It therefore respects repeated-candidate dependence across overlapping splits, which a naive within-split interval does not. It is conditional on the fitted models, the processed matrices and the endpoint definitions: it carries no retraining uncertainty, no donor-level transfer and no multiplicity adjustment across the seven tasks and three cutoffs. Intervals that narrowly exclude zero should be read accordingly. Mean outer-test Spearman correlation is primary; constant predictions receive zero correlation. AUROC and average precision use a top-decile outcome definition and are secondary; class-ineligible splits remain missing with their counts reported. Uncertainty uses 1,000 paired gene-cluster bootstrap draws, sharing each sampled gene’s weight across its repeated appearances in overlapping splits. Intervals condition on fitted models, processed matrices and endpoint choices. They are exploratory, unadjusted for multiple comparisons and do not represent independent-donor inference. In the paired-target experiment, readout genes are split once with seed 20261017; proxy PC1 uses label genes from anchor interventions only. Both endpoints share source feature genes, candidates and split indices. Primary protein predictions are rescored on replicate 3 and an ADT log-ratio without model refitting.

Scope and computation. Analysis plans and input hashes were fixed before their respective fits; functional-fitness, paired-target and full-profile extensions followed initial six-task findings and are explicitly exploratory. The reference grid retains its original preprocessing, which includes partly transductive feature construction; it is separated from the new training-only comparisons. Seven interface tests check information masking, held-out-label isolation, inductive graph behavior and statistical implementations. Full shared-space, native-tool and raw-atlas reruns and their limits are detailed in Supplementary Note 9.

Melanoma’s expression program score and scVI embedding are distinct quantities; latent-projection sensitivity is a separate analysis, not a validation of the expression score. Full supplementary methods retain the application and reference procedures.

## Data availability

All primary data are public and are available through the accessions listed in Supplementary Table 1, including GSE153056, GSE249595, GSE133344 and the pinned DepMap 24Q4 release (10.25452/figshare.plus.27993248.v1, CC BY 4.0). No restricted or controlled-access data were used in any reported analysis. Prepared matrices are checksummed and their provenance is recorded with the software; external matrices and pretrained weights require the documented source downloads.

## Code availability

InverseBench, the FSTN pipeline, the protocol interface and the analysis scripts are available from the corresponding author on request, and will be released publicly with an archival DOI minted from a tagged release before publication. Supplementary Table 6 records the frozen environment.

## Author contributions

Conceptualization: J.L. and C.T. Methodology: J.L. and Y. Luo. Software: J.L. Validation: J.L., L.F.H., Y. Luo and W.S. Formal analysis: J.L. and Y. Luo. Investigation: J.L. and Y. Luo. Data curation: J.L. and Y. Luo. Writing—original draft: J.L. Writing—review and editing: J.L., L.F.H., W.S., Y. Li, J.W.J., H.Q., Y. Luo and C.T. Visualization: J.L. and Y. Luo. Supervision: C.T. Clinical interpretation of the prioritization task and the nominated targets: J.W.J. and H.Q.

## Acknowledgements

We thank the authors of the public perturbation atlases and screens analysed here, listed with their accessions in Supplementary Table 1, for releasing the data that makes this benchmark possible.

