## Supplementary Information for "Information contracts determine what inverse perturbation benchmarks measure"

#### Supplementary Note 1: Validation of the melanoma demonstration system

These analyses support the melanoma demonstration and are, like the demonstration itself, a plausibility check rather than validated biology. FSTN is the name of the linear state-transition pipeline used throughout this note and in the released code: its response model is the continuous-target ridge of the main text, and the no-fitting comparator it is scored against is the projection reference.

##### FSTN's predicted effects match held-out perturbation experiments

Beyond ranking drivers, we asked whether FSTN's predicted effects match measured perturbation outcomes it never saw. Using a genome-scale CRISPRi Perturb-seq atlas of primary human CD4+ T cells  (11,526 genes perturbed across resting and stimulated conditions) as held-out ground truth, we defined an effector axis empirically as the resting-to-stimulated transcriptomic direction, estimated from training perturbations only, and measured each perturbation's effector effect as the projection of its differential-expression signature onto this axis.

On strictly held-out perturbations (60/40 split, 8,017 perturbations, axis from training only), FSTN's response model, mapping a perturbation's resting-state signature to its effector effect, achieved Spearman r = 0.60 (AUROC 0.86, precision@10 = 1.0, top-25 fold enrichment 8.4, permutation p < 5 x 10^-4); using the early, 8-hour signature improved this to r = 0.68 (AUROC 0.89, precision@10 = 1.0, fold enrichment 9.6, p < 5 x 10^-4; Extended Data Fig. 3b). Against a strong projection reference baseline that predicts each perturbation's late effect from its own early-timepoint shift along the effector axis , the learned linear response operator won decisively (AUROC 0.89 versus 0.62; Δ = +0.28 ± 0.02, ten of ten splits), the gain reflecting learning the multivariate early-to-late response rather than model capacity. This is a forward fidelity gain: it establishes that the response model learns real early-to-late structure, but does not translate into a driver-ranking advantage over other methods, since in the fair inverse benchmark the same supervised map added essentially nothing over a label-informed shared representation (0.888 versus 0.887). Equivalent held-out fidelity was obtained with a mechanistic backend that learns a regulatory response operator directly from the perturbation measurements (held-out Spearman 0.68); because it is estimated from actual interventions, it is a more direct causal object than a correlation-inferred network.

Because the genome-scale atlas is CD4+, we validated FSTN's reactive regulators against an independent CD8+ resource: pooled CRISPR knockout screens of transcription factors in human CD8+ T cells (GSE241933). FSTN's melanoma reactive transcription factors showed significantly larger CD8+ screen phenotypes than other genes (median absolute effect 0.30 versus 0.19; Mann-Whitney p = 4.7 x 10^-6), with FOXO1, GATA3, RUNX3, PRDM1, TBX21 and BATF3 among the strongest hits, confirming that the program FSTN recovers is functional in the cytotoxic lineage.

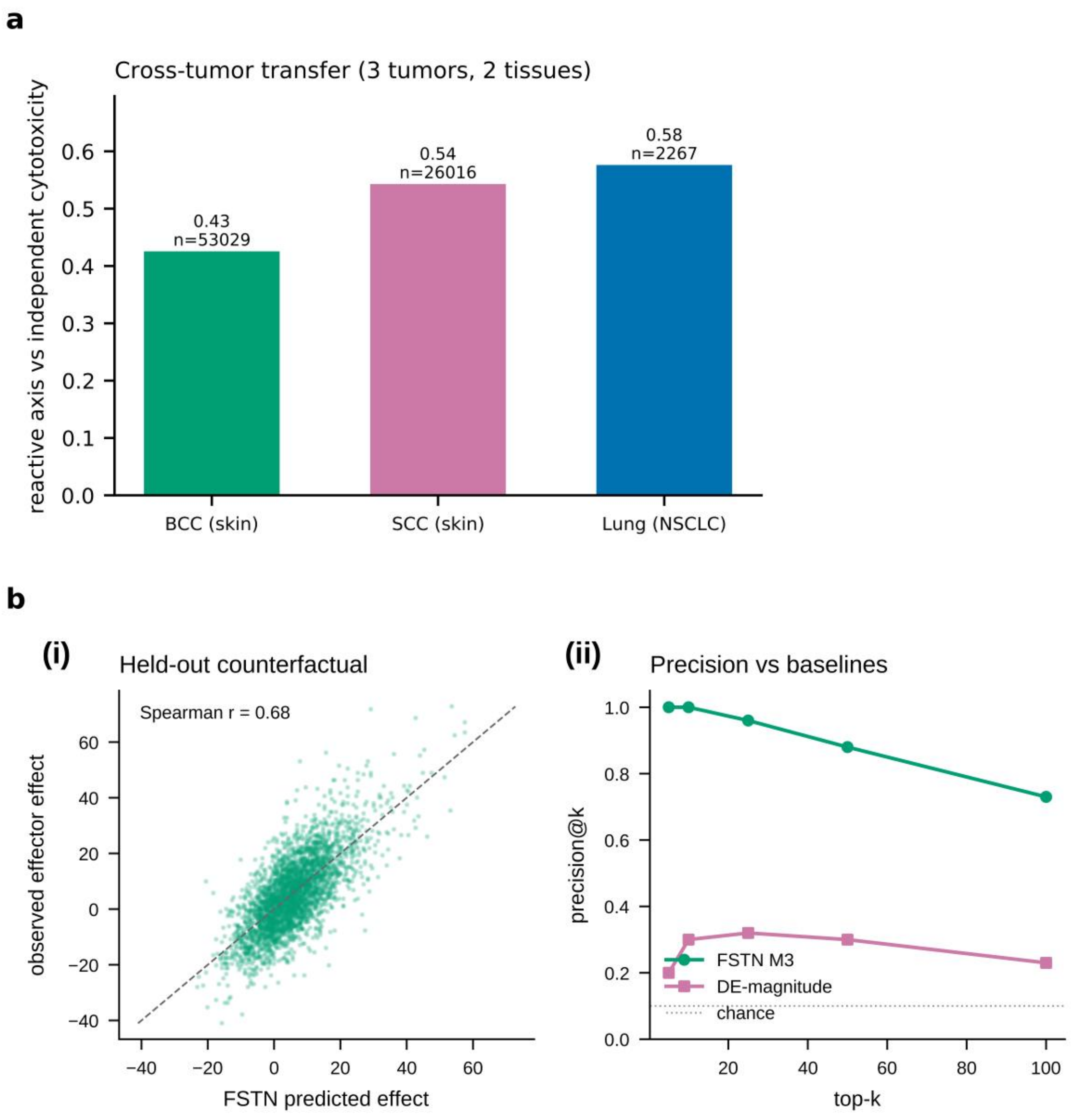


#### Under measured effects, a selected edit set moves real cells toward the target

As a controlled demonstration in the measured-effect regime, applying the selected edits to non-reactive source cells at fixed intensity using each edit's independently measured perturbation effect moved the source distribution onto the reactive attractor, overshooting the target mean by about 20% (fraction closed 1.21, where 1.0 lands on the target mean), whereas random edit sets of the same size and intensity closed only 0.11 (permutation $p < 0.002$). The five-edit set combined coherent T-cell effector regulators (VAV1, CD3E, MALT1, PLCG1) with one pleiotropic transcriptional adaptor (TADA2B). Because this loop uses measured effects as the actuator, it is an in-silico comparison of selection strategies in the measured-effect regime, not a forward experiment, and consistent with the decisive test above is not evidence that the pipeline prescribes unmeasured edits well; prospective wet-lab validation is the natural next step. Among the highest-ranked non-canonical edits were druggable signaling nodes such as ITK and MALT1, and the very top of the list is enriched for pleiotropic transcription and translation machinery, so these are hypotheses for follow-up rather than validated targets.

#### Transition programs generalize across patients and tumors

We tested generalization along two axes. Splitting melanoma samples into disjoint plate sets, the reactive-regulator ranking learned on one half was almost perfectly preserved in the held-out half (rank correlation +0.99, shuffled-label $p < 0.005$), providing a plate-split reproducibility check, without excluding all batch or donor effects (Extended Data Fig. 4a). Transferring the melanoma reactive-minus-naive signature to three unrelated tumors across two tissue types, it correlated with an independent cytotoxicity readout in each: basal cell carcinoma (53,029 cells, r = +0.43), squamous cell carcinoma (26,016 cells, r = +0.54; both GSE123813) and, from a separate study, non-small-cell lung cancer (2,267 cells, r = +0.58; GSE218262), all shuffled-label $p < 0.005$ (Extended Data Fig. 3a; summarised for basal cell carcinoma in Extended Data Fig. 4b). The regulatory logic FSTN learned in melanoma tracks the analogous state across skin and lung tumors.

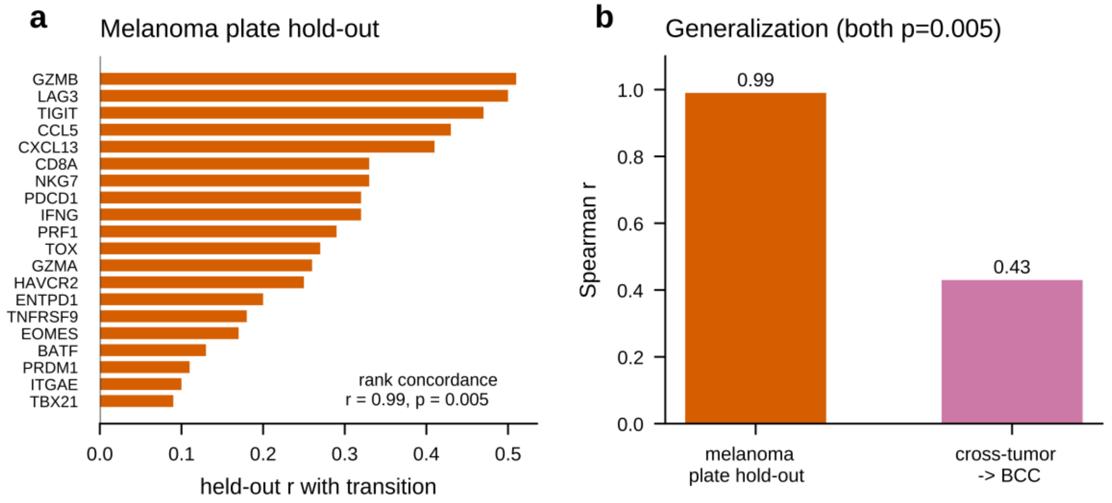


#### FSTN generalizes to a different cell type and transition

To test whether FSTN is a workflow on another transcriptional readout, we applied the identical machinery to an unrelated transition with its own perturbation ground truth: CRISPRa Perturb-seq in K562 leukemia cells driving erythroid differentiation (Norman et al. , 111,445 cells, 236 perturbations). Defining differentiation by an erythroid program score and fitting FSTN's response model on training perturbations, FSTN predicted the differentiation-driving effect of held-out perturbations (Spearman r = 0.99, AUROC 0.99, fold enrichment 3.8, permutation p < 5 x 10^-4). Integrity controls established genuine predictive signal rather than circularity: a shuffled target collapsed prediction to r = 0.02, and the differentiation readout was predicted consistently better than arbitrary gene-program scores (the high absolute correlation partly reflects the coherence of the differentiation program; the shuffle control is what establishes validity).

We also delineated where the approach does not transfer. Applying the identical two-condition design to a dendritic-cell LPS-activation Perturb-seq (Dixit et al. , 25 transcription-factor knockdowns), a perturbation's resting-state signature did not predict its activation outcome (in-sample Spearman approximately 0, held-out negative), indicating that the specific resting-to-activated predictive is transition-dependent rather than universal, and additionally limited by that dataset's small perturbation library. We report this null alongside the positive results.

#### Figures referenced in this note

The panels supporting this note are Extended Data Fig. 3a (cross-tumor transfer),

Extended Data Fig. 3b (held-out predictive fidelity) and Extended Data Fig. 4a,b

(plate hold-out generalization), whose legends are given with the Extended Data figures.

#### Supplementary application figures

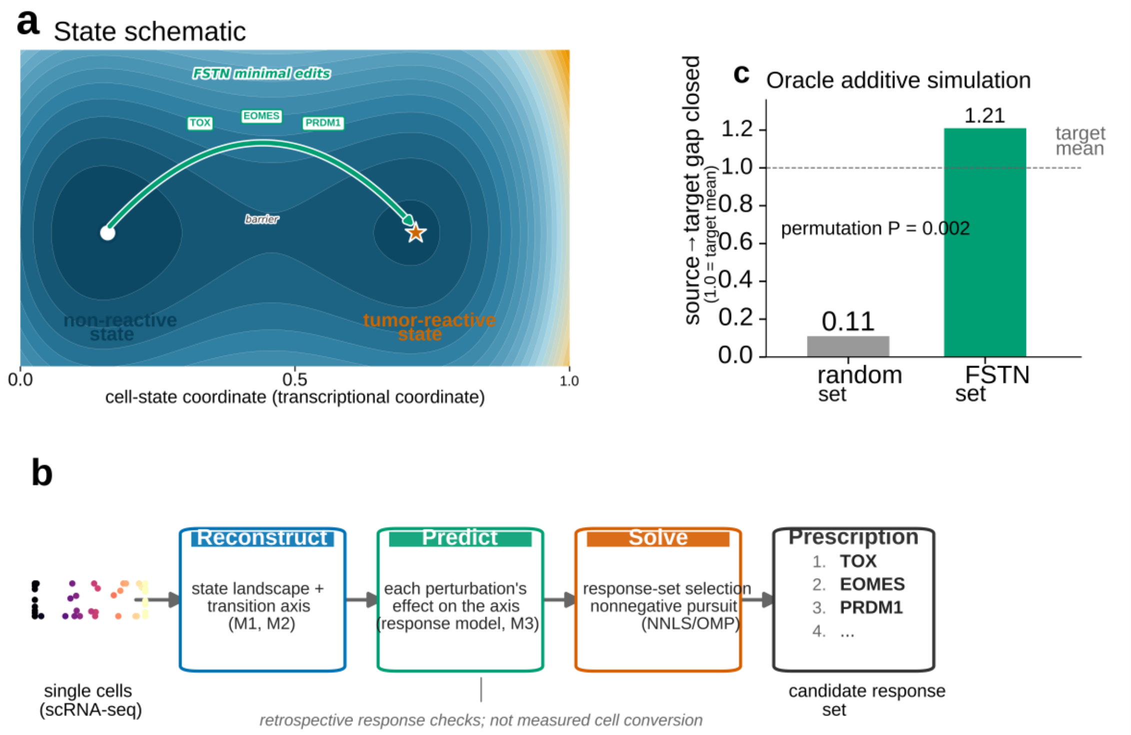


Supplementary Figure 1. Scope of response-informed prioritization. a, Schematic landscape illustrates a transcriptionally anchored coordinate; valleys, barrier and path are not inferred dynamical attractors or an observed conversion. Gene labels are illustrative. b, Workflow separates coordinate reconstruction, response ranking and heuristic NNLS/OMP selection. c, Oracle additive simulation under measured response effects: the selected set closes 1.21 of the target-axis gap versus 0.11 for random sets (permutation p = 0.002). This is modeled displacement, not an observed intervention or functional reprogramming result. Predicted-effect selection is evaluated separately in Supplementary Figure 3i.

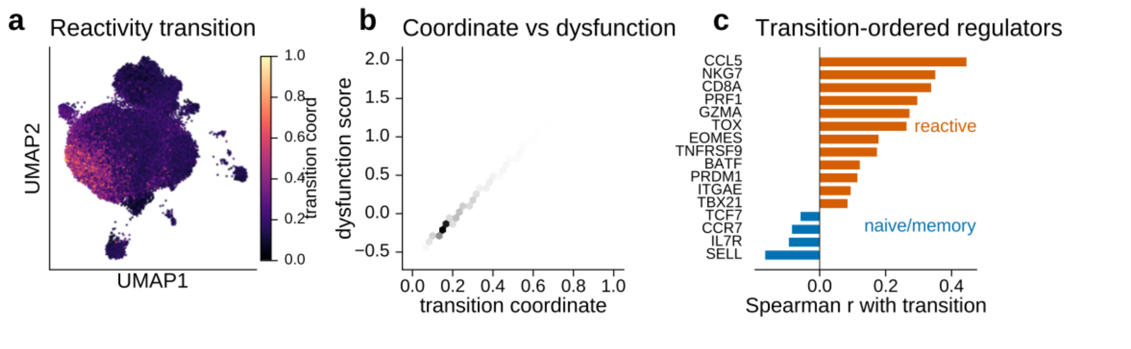


Supplementary Figure 2. Expression-defined melanoma program and visualization. a, UMAP of the scVI latent embedding of all 65,035 post-quality-control melanoma TILs (GSE123139), each cell coloured by its transition coordinate from 0 (naive and memory pole) to 1 (cytotoxic tumor-reactive pole). b, The same 65,035 cells as a hexagonal density plot of transition coordinate against the dysfunction and tumor-reactivity score, showing the defining expression score after rescaling; this is not an independent validation; shading is cell density, not a fitted model. c, Transition-ordered regulators, reactive versus naive and memory poles. The panel shows a canonical CD8 differentiation marker set, not the top of the ranking: three regulators correlate more strongly with the transition than the highest plotted gene (GZMB +0.50, LAG3 +0.49, TIGIT +0.46 versus CCL5 +0.45) and appear in the ranked candidate list instead. The complete ranking of all 36 regulators is Supplementary Table 3.

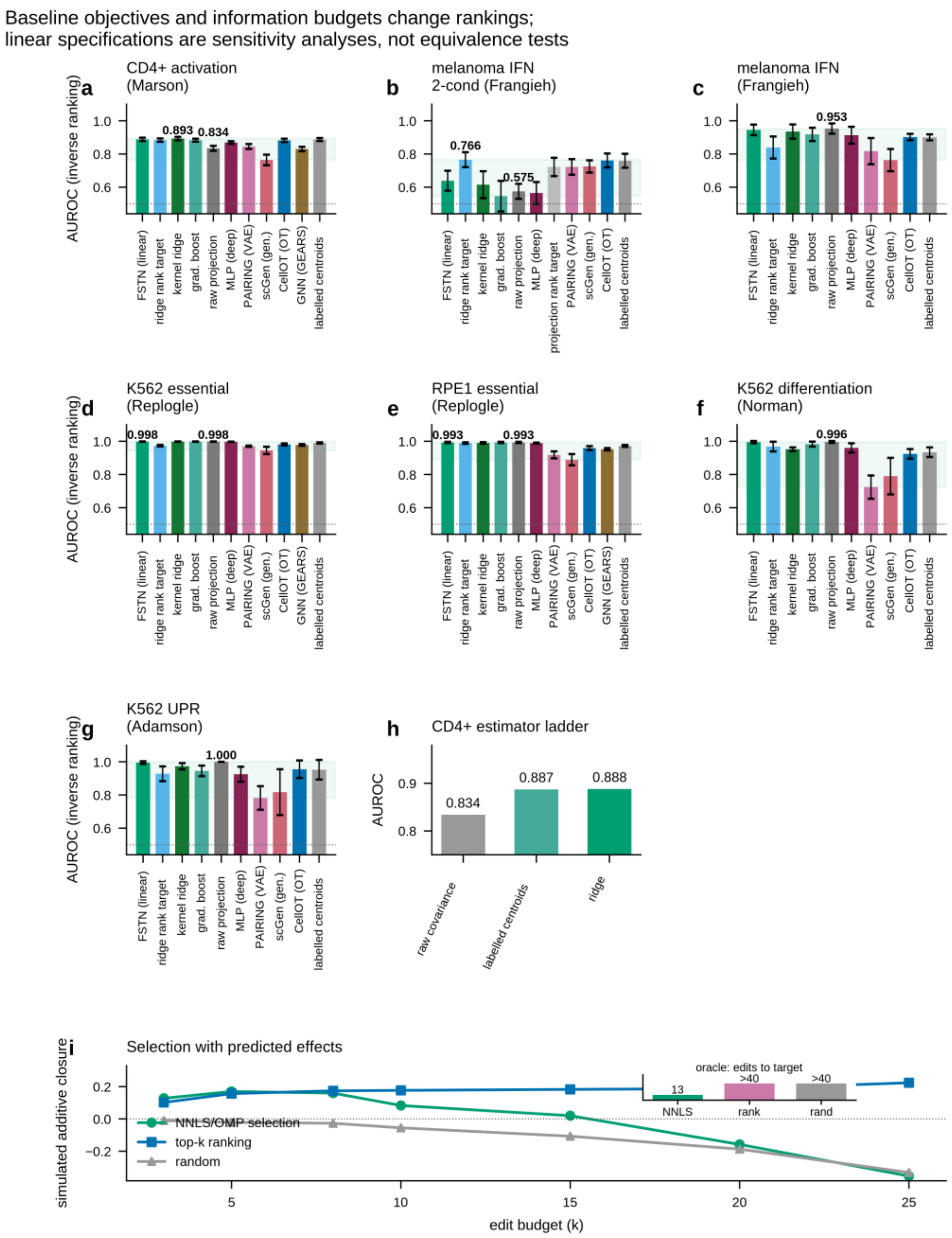


Supplementary Figure 3. Shared-space ranking references and selection sensitivities. a-g, Mean AUROC and s.d. over ten held-out-perturbation splits for CD4+, Frangieh cross-condition, Frangieh single-condition, K562, RPE1, Norman and Adamson. Single-condition tasks separate feature and label genes; cross-condition tasks separate measurements without a gene split. Both continuous-target and rank-target linear specifications are shown; the added rank-target bars show mean AUROC. Raw projection is feature-PC based on single-condition tasks and training-label covariance based on cross-condition tasks. No equivalence test was performed. Native tools are separate checks (Supplementary Note 5). h, CD4+ estimator ladder: raw covariance projection, label-informed centroid contrast and ridge. Source/target clouds use training outcomes. i, Simulated additive closure versus edit budget under predicted effects, with an oracle measured-effect inset. Negative closure at larger budgets is specific to the tested prediction/selection pipeline and is not evidence that all combination design fails.

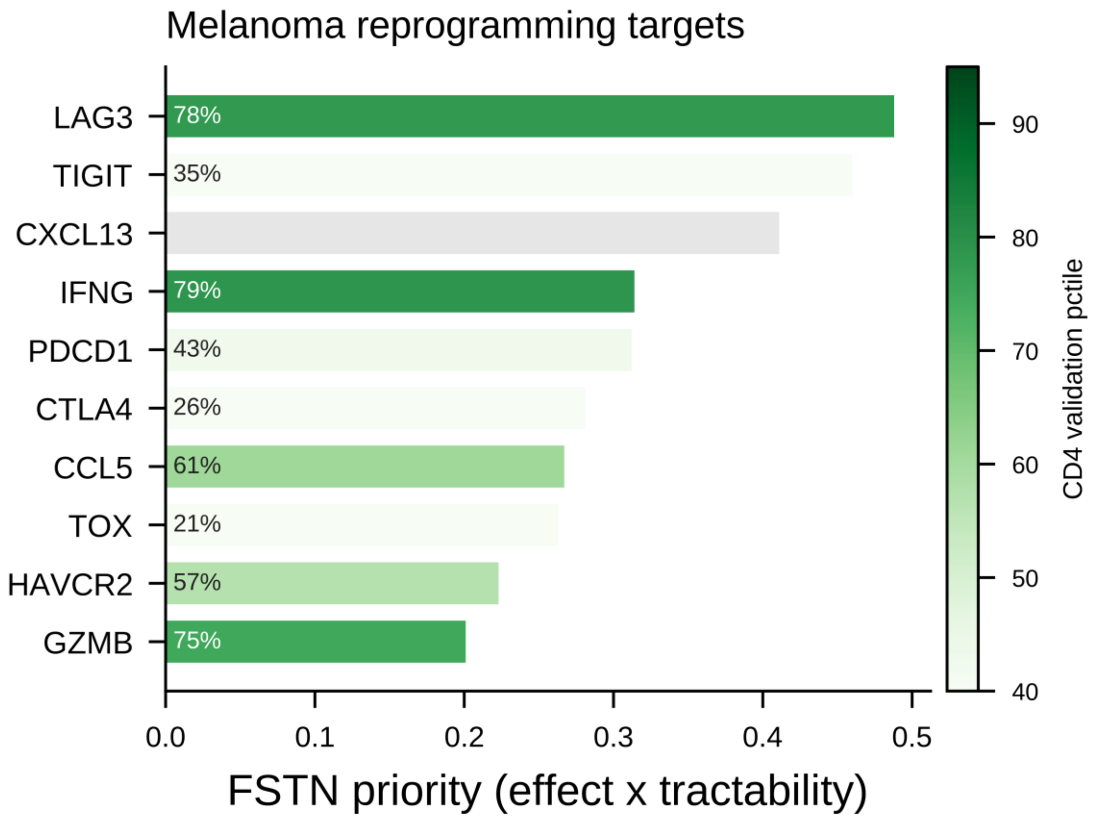


Supplementary Figure 4. Ranked melanoma reprogramming target list, with independent CD4+ cross-support. Bars are the FSTN priority score, the predicted effect on the melanoma reactive axis multiplied by an experimental tractability weight; bar colour and the inset percentage give each gene's percentile in the independent CD4+ perturbation atlas, so colour is a cross-dataset check and not an input to the ranking. Grey marks a gene not measured in that atlas (CXCL13). The ten highest-priority candidates of twelve are shown; the full list, including the two below the cut, is Supplementary Table 5. Note that one omitted candidate, ENTPD1/CD39 at rank 11, carries the strongest cross-support of all twelve (95th percentile, CD4+ effect +27.8), so the display cut is by priority and not by validation.

The published-tool illustration uses non-matched candidate sets and cannot establish comparative accuracy (Extended Data Fig. 1a,b).

#### Supplementary Note 2 | A conditional Gaussian result for covariance projection

##### Population setup

Let centered features x and scalar outcome y be jointly Gaussian, with positive-definite feature covariance S, positive outcome variance q and feature-outcome covariance vector v. For example, $x = A z + eps_x$ and $y = b^T z + eps_y$ give $v = A\ Sigma_z\ b$ under independent centered measurement noise. We do not require A to have full column rank; otherwise the no-signal case would be excluded for a nonzero b under positive-definite latent covariance.

The conditional mean is linear: $E[y|x] = w_{star}^T x$, where $w_{star} = S^{\{-1\}} v$. Compare it with the unwhitened covariance projection $s0 = v^T x$. The latter uses outcome covariance. In finite data, estimating v from paired training examples is a supervised procedure, even though no regression inverse is computed. A fixed PC projection is a different estimator unless an additional alignment condition identifies its direction with v.

##### Proposition and proof

If v = 0, joint Gaussianity implies independence of x and y and neither score contains outcome information. If v is nonzero, among linear scores the conditional-mean direction maximizes Pearson correlation. The squared correlations of the two scores are:

$$\text{optimal squared correlation} = (v^T S^{\{-1\}} v) / q$$
$$\text{projection squared correlation} = (v^T v)^2 / (q \cdot v^T S v)$$

Cauchy–Schwarz applied to $S^{\{-1/2\}}v$ and $S^{\{1/2\}}v$ shows the first is at least the second. Equality holds exactly when $S^{\{-1\}}v$ is a positive scalar multiple of v, equivalently when v is an eigenvector of S. Positive proportionality also gives identical rankings. Otherwise the optimal score is strictly better in population Pearson correlation. For Gaussian linear scores, Spearman correlation is a strictly increasing function of Pearson correlation, so the same ordering applies. This statement concerns these population linear directions; it is not a finite-sample guarantee for a fitted ridge or an AUROC equivalence test.

For a Gaussian threshold outcome with a fixed threshold on y, the conditional exceedance probability is monotone in $w_{star}^T x$ because conditional variance is constant. This motivates the direction for threshold ranking under the stated model. It does not show that least squares or a rank transform optimizes finite-sample top-decile AUROC under a non-Gaussian response model.

##### Measurement count is neither necessary nor sufficient

A single measurement may contain feature and label genes with different covariance geometry. Consider jointly Gaussian gene measurements (x1, x2, y) with covariance:

```
[2,28,1]
[28,1]
[1,3]
```

With x1 and x2 as features, v = (1,1) and S = diag(2,1). Covariance projection is x1+x2, whereas the optimal score is 0.5x1+x2. Their population correlations are 0.667 and 0.707. Thus one measurement with a gene split can admit an improvement. Conversely, separate source and target measurements can preserve the relevant eigenvector, yielding no population gain from whitening.

The single-condition PC1 tasks in this benchmark have strongly correlated feature- and label-gene axes. This is an empirical property of their construction, not a theorem about all single-condition tasks. The benchmark also compares different feature spaces, truncations and target transformations; their differences are not exhausted by a whitening factor in one fixed coordinate system.

#### Empirical scope

CD4+ and Frangieh IFN show gains for selected supervised estimators over their projection references. Frangieh co-culture has weak measured predictivity, while Schmidt shows no detected gain against its supplied target axis. Weak predictivity is not zero mutual information, and a null comparison on a small library is not evidence that a projection is Bayes-optimal. Schmidt's curated-target result in Supplementary Note 7 additionally demonstrates target dependence.

Rest-to-8 h, rest-to-48 h and 8 h-to-48 h comparisons reuse the CD4+ atlas, perturbations and donors. They are within-atlas sensitivities rather than independent replications of a structural law. Exploratory screens rejected for sparse measurements or small libraries are dataset-suitability observations, not scored evidence for the proposition.

#### Synthetic illustrations and validation limits

Let L denote the target measurement matrix, and define the scalar phenotype score as $y = u^T Lz$, where u weights the measured features. Then $y = (L^T u)^T z$, so the phenotype coefficient in latent space is $b = L^T u$. The analytical examples use this definition; replacing the transpose by a pseudoinverse generally defines a different phenotype. The result does not establish a general measurement-count criterion.

Neither readout divergence nor angular misalignment alone determines the gain: eigenvalues, signal strength and noise also matter. A monotone divergence sweep in one simulation would not prove a general monotonicity result. Deterministic numerical checks verify the eigenvector identity, a positive-definite same-measurement counterexample and a different-measurement equality case. These checks validate the population algebra, not fitted-model performance on real systems.

We have not tested a training-only geometry diagnostic that predicts gains on independent real systems. That experiment is required before treating covariance geometry as an established real-data mechanism rather than a conditional explanatory result.

#### Supplementary Note 3 | Measured constituent-pair retrieval

This separate retrospective task asks whether measured single-response signatures identify the interventions that generated a measured double-response target. It evaluates constituent retrieval, not deployment of predicted combinations or a proof of minimal intervention design.

##### Answer key

The Norman 2019 K562 combinatorial CRISPRa screen measures 131 two-gene perturbations together with all their constituent single perturbations (105 single-gene signatures with at least 20 cells each). For a two-gene target state produced by genes A and B, the known generating pair is {A, B}. This does not establish that no other single intervention or combination could produce a similar phenotype. The task: given the control-referenced target signature and the single-gene library, select a small set of singles whose combined (additive) effect reaches the target, and score whether the selected set recovers {A, B}.

##### Selectors

Three selectors are scored under measured single-perturbation effects. Algorithm 1, nonnegative orthogonal matching pursuit with a joint NNLS refit after every pick, is run directly. It is compared with an unrestricted greedy residual, which takes signed scalar steps and never refits earlier coefficients, so four of 262 coefficients in a repeat check of its first two picks were negative and could not be realised by a nonnegative dose model, and with independent cosine ranking.

Independent ranking scores each single g by cosine($s_g$, $s_{target}$) and takes the top k. It has no redundancy awareness, so it can pick two singles pointing the same way.

Redundancy-aware (greedy residual, the paper's set-selection idea) iteratively adds the single that most reduces the residual target − sum(selected singles), so each pick must explain a direction not already covered.

##### Result

Mean over the 131 two-gene targets (chance recall@2 = 2/105 = 0.019):

| selector | recall@2 | recall@3 | recall@5 | exact-pair@2 |
|---|---|---|---|---|
| independent ranking | 0.546 | 0.615 | 0.706 | 0.198 |
| redundancy-aware | 0.714 | 0.771 | 0.855 | 0.511 |

recall@k is the fraction of the true two-gene set recovered within the top k selected singles; exact-pair@2 is the fraction of targets for which both constituents are recovered in the first two picks. Redundancy-aware selection recovers the true constituent pair 2.6-fold more often than independent ranking (exact-pair 0.511 versus 0.198), and dominates at every budget. Genetic interactions can make the double signature differ from the sum of its singles ($s_{AB}$ != $s_A$ + $s_B$). This complicates response reconstruction but does not establish a theoretical ceiling on constituent identification. The reported means are descriptive; reused genes across targets induce dependence.

These measured-effect means support a constituent-retrieval advantage for this signed residual heuristic over independent cosine ranking on this library. They do not validate the nonnegative solver, establish unique minimality or demonstrate cell conversion.

#### Algorithm 1 scored against the alternatives

131 two-gene Norman targets, 105-gene single library, measured effects.

| selector | recall@2 | recall@3 | recall@5 | exact-pair@2 |
|---|---|---|---|---|
| independent cosine ranking | 0.546 | 0.615 | 0.706 | 0.198 |
| unrestricted greedy residual | 0.714 | 0.771 | 0.855 | 0.511 |
| Algorithm 1 (nonnegative OMP + NNLS) | 0.725 | 0.840 | 0.916 | 0.527 |

Paired per-target bootstrap, 2,000 draws, Algorithm 1 minus the unrestricted greedy: recall@2 +0.0115 [+0.0000, +0.0267]; exact-pair@2 +0.0153 [+0.0000, +0.0382]; recall@3 +0.0687 [+0.0382, +0.0992]; recall@5 +0.0611 [+0.0344, +0.0917]. The constraint earns its place at larger budgets; at two picks the advantage is not distinguishable from zero. 129 of the 131 targets share a constituent gene with another target, so resampling targets does not remove gene-level dependence and these intervals are optimistic.

#### Supplementary Note 4 | A gene-regulatory-network-inversion method arm

The eight-method grid covers linear, kernel, tree, generative, optimal-transport and graph-neural classes, but not the gene-regulatory-network (GRN) inversion class that a large part of the field uses to prescribe transcription-factor edits. This note adds that class and shows the two-regime finding extends to it.

##### Method and dataset

TFcomb inverts a base GRN to rank the transcription factors (TFs) whose activation best drives a source cell state toward a target state (a Tikhonov-regularised inversion of the regulatory map; from the scDirect/TFcomb codebase, github.com/Chen-Li-17/TFcomb). We ran it on the tool's shipped real paired scRNA+scATAC demo, human fibroblast-to-iPSC reprogramming (figshare article 28138355; 7,841 cells x 1,838 genes; base GRN 31,844 peaks over 1,100 genes built from the scATAC via CellOracle). The scATAC provides the base regulatory network; the scRNA provides the source (fibroblast) and target (iPSC) states. The ground-truth minimal driver set is the Yamanaka reprogramming factors {POU5F1, SOX2, KLF4, MYC, NANOG}. KLF4 is absent from the base-network TF pool, so the maximum achievable recall of the five-factor key is 0.80.

##### Comparison

We scored TFcomb head-to-head against two linear baselines on the same TF pool and answer key:

linear regulon projection, a one-step linear read over the same base GRN (rank TFs by the alignment of their regulon with the source-to-target expression shift); isolates what TFcomb's inversion adds over a single graph step.

linear self-DE (no network): rank TFs purely by their own up-regulation in the target state; a no-network, projection reference analogous to the additive projection in the main benchmark.

##### Result

| method | recall@10 | recall@20 | transition-improvement (top 20) | median driver rank |
|---|---|---|---|---|
| TFcomb inversion | 0.40 | 0.60 | 0.48 | 8.5 |
| linear regulon projection (same GRN) | 0.40 | 0.60 | 0.42 | 11.5 |
| linear self-DE (no network) | 0.60 | 0.80 | 0.57 | 6.5 |

Per-TF ranks under TFcomb: POU5F1 = 1, MYC = 2, SOX2 = 15, NANOG = 22 (KLF4 not in the pool).

Two things hold at once. TFcomb genuinely works as a GRN inverter: it places the two master regulators POU5F1 and MYC at ranks 1 and 2, and it beats a one-step linear projection over the same network (transition-improvement score 0.48 versus 0.42), so its inversion extracts signal the single graph step does not. But it does not beat the no-network baseline that ranks TFs by their own target up-regulation, which recovers the driver set earlier on every aggregate (recall@20 0.80 versus 0.60, median rank 6.5 versus 8.5, transition-improvement 0.57 versus 0.48).

#### Reading

This is the two-regime finding in an independent method class and an independent transition. Fibroblasts and iPSCs are read out in the same single-cell RNA measurement, a single-readout system, so the transition axis is directly legible in the features and a projection reference is near-optimal, just as on the five single-condition Perturb-seq systems in the main benchmark. The GRN inversion adds value over a same-network linear step, but not over reading the transition directly from a single measurement. Consistent with the paper's central point: elaborate machinery earns its keep only where the features and labels come from different measurements.

#### Supplementary Note 5 | Released tools in their own native feature space

The benchmark ranks every method in a shared neutral latent, imposed for fairness: it removes the feature-space asymmetry that would otherwise let a method with a richer representation appear to win. A skeptic could ask the opposite: does the shared latent instead handicap the deep methods, so that the linear-beats-deep result is an artifact of the protocol rather than of the methods? This note provides limited checks by running the authors' own released code in each tool's native feature space and scoring it on the identical held-out driver labels.

The numbers in this note are the historical native-tool runs. Supplementary Note 9 reports the completed reruns, revised numerical values and GEARS comparisons on matched covered candidates.

##### Protocol

For each tool we reproduce the benchmark's family-specific label definition and held-out-perturbation-row split and top-decile driver labels byte-for-byte (asserted equal to the stored per-split label dumps before any score is computed), and change only one thing: the tool ranks in its OWN native feature space rather than in the shared PCA-30 latent. Ten splits per system; AUROC on the held-out drivers.

##### CellOT (released code, native gene space)

We ran the released CellOT (cellot.networks.icnns.ICNN + cellot.models.cellot) to learn the source-to-target optimal-transport map in the full train-gene signature space, then ranked candidates by the alignment of their native signature with the learned transport displacement. Run on SLURM (gen-cpu, array over splits, fresh env torch 1.11.0 / py3.9.5).

| system | regime | native CellOT | linear FSTN | projection reference |
|---|---|---|---|---|
| CD4+ | two-measurement (learnable map) | 0.837 +/- 0.015 | 0.888 | 0.834 |
| Norman | single-condition (saturated) | 0.997 +/- 0.004 | 0.995 | 0.996 |
| Adamson | single-condition (saturated) | 1.000 +/- 0.000 | 0.995 | 1.000 |

On CD4+, the one system where a learnable cross-measurement map exists and methods separate, native CellOT scores 0.837, below linear FSTN (0.888) and at the projection reference (0.834). On the saturated single-condition systems it ties the additive ceiling (this is the saturation regime, not an edge: the additive projection matches it).

##### GEARS (released code, native gene space)

We ran the released GEARS (cell-gears 0.1.2) in its own native ~5,000-HVG log-normalized gene space on Norman 2019 (CRISPRa, the native-fit genetic-perturbation system), training a full

GEARS model per split and ranking held-out perturbations by the projection of GEARS' predicted delta-expression onto the oriented held-out driver axis. Because CPU training was infeasible (~2.7 h per epoch), we trained properly on an NVIDIA H100 (one GPU per split, 20 epochs), with the GO graph pre-built once to avoid a relative-path write in the released code. AUROC is computed on the GEARS-predictable subset of held-out rows (coverage reported).

| system | regime | native GEARS (H100, 20 ep) | linear FSTN | projection reference |
|---|---|---|---|---|
| Norman | single-condition (saturated) | 0.922 +/- 0.039 | 0.995 | 0.996 |

Per-split AUROC: 0.966, 0.943, 0.901, 0.966, 0.921, 0.888, 0.882, 0.980, 0.913, 0.857 (coverage 92-95 of 95 held-out rows; 5-14 positives per split). The fully trained released GEARS, in its native representation, scores 0.922, below linear FSTN (0.995) and below the projection reference (0.996).

#### Reading

Native CellOT is below the ridge reference on CD4+ but has slightly larger means on Norman and Adamson. These checks do not establish a representation-independent capacity theorem. Native GEARS is scored on a predictable subset, so its displayed comparison to full-library reference means is contextual rather than a strictly matched-candidate inferential comparison. Common-subset rescoring and matched tuning budgets remain required before making a definitive native-tool ordering claim.

#### Single-cell foundation-model arm (Geneformer)

The same holds for a single-cell foundation model: Geneformer, scored by the identical leakage-free displacement in its pretrained embedding rather than in the shared latent, does not beat the linear model on either single-condition system it can be run on (0.991 versus additive 0.996 on Norman; 0.901 versus 0.953 on Frangieh, losing on 9 of 10 splits), reproducing the recent finding that transcriptomic foundation models do not beat linear baselines on perturbation tasks.

A gene-regulatory-network inversion arm, which solves the prescriptive task from an inferred network rather than from measured responses, is reported separately in Supplementary Note 4.

#### Supplementary Note 6 | Canonical reference protocol and uncertainty

##### Reference task families

The reference benchmark uses one canonical protocol module, shared by the registry, evaluator and reference methods. Its two task families declare distinct preprocessing and label populations:

Two-measurement systems: CD4 activation, Frangieh IFN/co-culture, Schmidt and additional within-atlas CD4 pairs use source-condition signatures and a supplied target-condition outcome, without a readout-gene split.

Single-condition systems: Frangieh, K562, RPE1, Norman and Adamson use a 60/40 readout-gene partition and a 60/40 candidate-row partition. Feature and label axes are computed from distinct readout genes, with top-decile outcomes defining drivers.

These reference procedures are separate from the disjoint-anchor, training-only transformations of Supplementary Note 10.

##### Declared preprocessing populations and method boundary

In two-measurement systems PCA fits on training feature rows and scaling uses all feature rows. The external target-condition axis is supplied by the loader and is not a per-split trained axis. In single-matrix systems the feature-PC axis, label-PC axis and shared PCA fit on all rows; scaling also uses all feature rows. The label axis is signed against the feature axis over the candidate population. Thus the protocol is not strictly inductive in preprocessing. Model parameters fit only on training perturbations.

The evaluator retains held-out labels outside the method context. A method receives features, training outcomes, training-defined directions and evaluation indices only. Interface tests exercise both task families. This is an information contract, not an operating-system sandbox against arbitrary Python reading public files.

##### Exact reproduction, not agreement of a rounded mean

Two different protocols can land on the same third decimal of a mean AUROC, so agreement of a summary statistic is not evidence that a harness reproduces a published run. The reconciliation is therefore tested against the stored per-candidate score vectors of the published grid, split by split, with three criteria required to hold exactly: the scored-row count, the driver labels, and the induced ranking, the last of these because AUROC is a function of the ranking alone and those three conditions are together sufficient for the checked comparisons for the published AUROC to be recovered. On all seven systems whose arrays are available, the canonical protocol reproduces the reference baseline ranking exactly on all ten splits, with residual value drift of order 1e-5 attributable to the float32 precision at which the systems' arrays and score dumps are stored (the one system stored at double precision agrees to 3e-14). The baseline self-test tolerance is 0.002. This baseline check does not claim exact reproduction of neural score vectors; their numerical sensitivity is reported separately in Supplementary Note 9.

#### The frozen split manifest

The release ships one row per system and split recording the candidate count, the training and scored row sets as hashes of their sorted index arrays, the number of drivers among the scored candidates, the feature and label gene sets where a gene split exists, the axis-fit population, the orientation rule, and the preprocessing-fit population. Identical hashes mean identical held-out sets, so a reproduction can be verified without shipping index arrays and a mismatch localises to a split rather than to a disagreeing average.

| system | family | candidates | train rows | scored rows | drivers among scored |
|---|---|---|---|---|---|
| CD4+ | two-measurement | 8,345 | 5,007 | 3,338 | 349 |
| Frangieh control-versus-IFNγ | two-measurement | 235 | 141 | 94 | 8 |
| Frangieh | single-condition | 246 | 147 | 99 | 11 |
| K562 | single-condition | 11,258 | 6,754 | 4,504 | 466 |
| RPE1 | single-condition | 2,679 | 1,607 | 1,072 | 113 |
| Norman | single-condition | 236 | 141 | 95 | 11 |
| Adamson | single-condition | 107 | 64 | 43 | 5 |

The last column is worth reading directly: Adamson contributes five drivers among forty-three scored candidates per split, which is the honest reason its intervals are wide and a reason not to lean on small differences there.

#### Baseline objective sensitivity

The continuous-outcome ridge scores 0.639 on Frangieh IFN while CellOT reaches 0.761. The label-informed centroid contrast scores 0.759. Rank-transforming training outcomes raises ridge to 0.766 and raw covariance projection from 0.575 to 0.722. This is an exploratory specification analysis prompted by the original discrepancy, applied to all grid systems and reported alongside the original estimates.

| comparison (linear minus comparator) | mean gap | perturbation-level 95% CI |
|---|---|---|
| rank-target linear vs CellOT | +0.005 | [−0.022, +0.029] |
| rank-target linear vs PAIRING | +0.044 | [−0.005, +0.093] |
| rank-target linear vs scGen-style | +0.040 | [−0.015, +0.089] |

These intervals do not establish equivalence. Rank-transformed squared loss is not direct optimization of top-decile AUROC. No independent validation set was used to prospectively select between the two specifications; reporting both is transparency about sensitivity, not an out-of-sample selection rule. Both the raw covariance reference and the centroid direction use training outcome labels and are supervised.

The rank-target specification is worse on several single-condition tasks, including Adamson (0.928 versus 0.995) and Frangieh (0.839 versus 0.945). A fair capacity comparison would permit comparable training-only objective and hyperparameter selection for all learners. No universal claim that linear models suffice follows from this analysis.

#### Uncertainty with repeated candidates

The intervals first reported resampled candidates independently inside each of the ten splits and then averaged the ten gaps. The ten splits are overlapping random 40% row subsets of one perturbation pool, so a perturbation recurs across several of them, and independent within-split resampling treats those recurrences as independent observations. That understates the variance of the averaged gap, which makes the intervals too narrow. Two estimators that do not assume independence are now reported. The first takes the perturbation as the resampling unit: each draw resamples perturbation identities with replacement from the pooled candidate set, and a drawn identity carries all of its occurrences across every split, preserving the cross-split dependence. The second resamples split-level paired gaps when candidate-level reconstruction is unavailable. Splits overlap in training and evaluation populations, so this is not guaranteed conservative or an independent-sample estimator. It is a conditional sensitivity summary; the graph-network comparison also uses it.

Recovering perturbation identity required reconstructing each split's held-out indices. The row split is deterministic given its seed, but the held-out size does not determine the pool size uniquely, since 234 and 235 candidates both leave 94 rows held out, so every admissible pool size is tried and only a reconstruction that passes two checks is used: a perturbation held out in two splits must carry the same stored effect value in both, and across all splits every candidate labelled a driver must have an effect above every candidate labelled a non-driver. A system that fails those checks is reported as unreconstructible and falls back to the split-level estimator rather than being resampled on a guess.

The correction widens the intervals in the expected direction and changes some interval interpretations; main claims use the explicitly stated estimator. On CD4+ the headline gap of a learned model over the projection reference is +0.053 with a perturbation-level interval of [+0.043, +0.064], against the within-split interval [+0.048, +0.059], so it is about 1.8 times wider and still excludes zero. The small margins survive too: the linear model over the neural transport model is +0.006 with interval [+0.003, +0.009], and the kernel-ridge edge over the linear model is +0.005 with interval [+0.003, +0.007]. On the boundary system that fails the learnability condition every comparison spans zero under both estimators, without proving either a lack of information or an optimal projection. The table states which estimator applies. Candidate resampling conditions on the fitted models and does not capture donor variation. Historical adjusted p-values and the new cluster intervals should not be read as a newly corrected, exhaustive family of tests across all exploratory targets and specifications.

#### Canonical execution and environment sensitivity

The main compute worker and the objective-sensitivity runner now call the canonical split functions instead of maintaining independent implementations. The actual worker entry point is regression-tested under both protocol families, and every new score dump retains training rows, held-out rows, driver labels and its protocol manifest. Ninety additional control splits cover actual-target PC2 training and the Track B asymmetry experiments described in Supplementary Note 7.

The expanded historical reference check covers ten systems with available score dumps, including all seven main-grid configurations. Datlinger has no published dump and is reported as unverified. Labels and the rankings of ridge, raw projection and centroid contrast match the historical scores on all checked splits. Row identities are also compared directly where the historical dump retains them; the new dumps preserve those identities in every case.

All nine worker scorers have additionally been rerun across the seven main-grid configurations under both the frozen-model environment and a newer environment, with explicit thread settings. The shared-space GNN has been rerun across its three evaluated systems and now retains individual candidate scores. These reruns support the main comparison pattern, but not every neural score vector reproduces the historical floating-point trajectory. Numerical differences remain even when using the recorded frozen-model package versions. The comparison files retain every split's mean metrics and differences; no claim of bitwise neural reproduction is made. Reference Table N10a retains the historical frozen results, rather than silently substituting a selectively chosen rerun.

The newer- and frozen-environment runs are computational sensitivities on the same datasets, not independent replications or evidence identifying a unique cause of numerical drift. The software release records environments, outputs, protocol tests and complete comparison reports. Four-system baseline execution has also been checked from a fresh standalone installation. Completed native-tool fits, full raw-atlas reconstruction, scVI training and count-based sensitivities are reported in Supplementary Note 9. These project-environment computations do not establish complete clean-host reproduction of every external tool or historical figure.

#### Supplementary Note 7: The information budget, and what the benchmark does and does not measure

This Note addresses a scope question that applies to the whole resource: what is each candidate allowed to be described by, and what task does that make this. It separates the benchmark into two tracks, measures the harder one rather than asserting that the easier one stands in for it, and tests whether the single-condition conclusions depend on how the target was defined.

##### The budget in the published grid, stated plainly

In the grid of Table 1 a candidate perturbation is represented by its own already-measured signature: in the two-measurement systems its differential-expression profile in the source condition, and in the single-condition systems its profile in the train-gene half of the one signature matrix. Everything the benchmark rewards is therefore conditional on a screen having already been performed on every candidate it ranks.

The deployment scenario this matches is cross-context prioritization. A screen exists in an accessible context, and the question is which of those measured candidates moves the cell along a transition defined in a context that was not screened, or along an axis read from genes held out from the features. This is a genuine and common problem, because the accessible screen is usually cheaper than the target-condition experiment one actually wants. It is nonetheless weaker than the strong reading of inverse design, in which an intervention is nominated from an unperturbed source state and a desired target alone, having never been performed on anything.

We therefore distinguish, and name, two tracks:

Track A, response-informed prioritization. The candidate carries its own measured source-condition signature. This is the published grid. A projection reference is well defined here, because there is a signature to project onto the transition axis, which is why the central comparison of this paper is possible at all.

Track B, unseen-perturbation ranking. The candidate may be described only by information obtainable without perturbing it. No projection baseline exists, because there is no signature to project. Non-trained priors and heuristics remain possible; the own-response projection from Track A is unavailable.

##### Track B: the construction, and why it is not a strawman

A candidate gene g is represented by its behaviour as a readout gene across the training perturbations, that is by column g of the training-row signature matrix: how g's own expression moves when other genes are perturbed. Observing that never requires perturbing g. For a held-out candidate its own row is in the test set by construction, so a held-out candidate never enters its own feature vector. Training candidates do have their own row among the training interventions, an asymmetry in the historical arm, now tested by masking and disjoint-anchor controls below. The label is unchanged from Track A, the effect of perturbing g on the target axis, so the two tracks are directly comparable on the same candidates and the same labels.

This is the class of side information that released methods use to generalise to unseen perturbations: a GEARS-style gene graph exploits exactly this co-response and co-expression structure. Track B is restricted to candidates whose perturbed gene is also among the measured readout genes, which is 96% to 100% of the library on the systems where it could be built, and that restriction is reported with each result.

Four representations/heuristics were explored, and the best observed score is reported descriptively. This limited search does not establish intrinsic task difficulty or eliminate representation misspecification:

1. the co-response profile in the source condition;

2. the same profile in the target condition, which is still within budget, since observing how g

responds to other perturbations never requires perturbing g, and which lives in the condition the label is read from;

3. the candidate's overall response amplitude, a no-learning Track B floor;

4. a k-nearest-neighbour predictor assigning a candidate the mean measured effect of the training

candidates whose co-response profiles most resemble its own.

The source profile is tested with ridge and kernel ridge; the target profile is tested with ridge. These are the implemented arms, not the full native graph/deep-model suite.

#### Target-direction robustness on the single-condition systems

A second scope question concerns the label rather than the features. On the five single-condition systems the target is the top principal component of the held-out genes, described as differentiation, immune response or fitness. PC1 of a response matrix is not an independently specified biological destination, so a near-perfect AUROC there may partly reflect that the benchmark reads its target off the dominant direction of the very variation it then asks methods to rank.

The scope of that objection is worth stating precisely: it reaches the single-condition family only. In the two-measurement family the target is a control-derived condition-contrast axis, fixed before any candidate is ranked, so the label is not a principal component of the ranked matrix at all.

Two controls test the single-condition family on identical splits, candidates and methods:

A subdominant target. The label axis becomes the second principal component of the held-out genes. The existing code retains the feature-PC1 training target, so this arm only rescores the original rankers; it does not ask whether a learner trained against PC2 can improve on an appropriate PC2 reference.

A residualized analysis. The systematic (average-perturbation-effect) direction is projected out of every signature, features and labels alike, and the feature axis, the shared latent and the label are all recomputed from the residualized matrices. This is the direct control for the axis-alignment concern. The systematic axis is the mean of the uncentered signature matrix, since the signatures are already differential effects relative to control; taking the mean of a column-centred matrix returns zero and makes the control silently vacuous, which is a trap we fell into and caught by noticing that it changed nothing at all.

These historical arms are descriptive perturbations of target and representation. The target-trained PC2 experiment below supplies the actual target on training rows and fits its corresponding covariance reference.

#### Track B results under the tested representations

| system | candidates | best Track B | Track A projection | Track A learned | random floor |
|---|---|---|---|---|---|
| CD4+ activation | 8,326 | 0.585 | 0.833 | 0.888 | 0.493 |
| Frangieh control-versus-IFNγ | 226 | 0.582 | 0.585 | 0.639 | 0.499 |
| Frangieh control-versus-co-culture | 225 | 0.566 | 0.586 | 0.534 | 0.510 |
| Schmidt rest-versus-restimulated | 72 | 0.589 | 0.828 | 0.820 | 0.509 |

The tested baselines score 0.566 to 0.589. These are limited baseline results, not an upper bound on unseen-perturbation methods. Different library sizes are different biological systems and do not constitute a controlled learning curve or power analysis. Taking the best observed arm is exploratory selection, not a performance estimate of a prospectively chosen method.

On matched candidate sets, CD4+ projection scores 0.833 versus 0.585 for the best Track B arm, and Schmidt scores 0.828 versus 0.589. These gaps compare different representations and estimators and have not been assigned a model-independent causal interpretation or paired uncertainty interval. Track B also permits target-condition training profiles in one arm. Both candidate coverage and this budget must be stated when comparing results.

#### Target-direction robustness: results

| system | target | learned | no-learning | learning gain | systematic variance removed |
|---|---|---|---|---|---|
| Frangieh | PC1 | 0.945 | 0.953 | -0.008 (4/10) | |
| Frangieh | PC2 | 0.585 | 0.583 | +0.002 (5/10) | |
| Frangieh | residualized | 0.964 | 0.956 | +0.008 (5/10) | 0.166 |
| K562 | PC1 | 0.998 | 0.998 | -0.000 (3/10) | |
| K562 | PC2 | 0.705 | 0.705 | -0.000 (2/10) | |
| K562 | residualized | 0.995 | 0.995 | +0.000 (4/10) | 0.073 |
| RPE1 | PC1 | 0.993 | 0.994 | -0.000 (2/10) | |
| RPE1 | PC2 | 0.813 | 0.813 | -0.000 (7/10) | |
| RPE1 | residualized | 0.832 | 0.832 | +0.000 (6/10) | 0.334 |
| Norman | PC1 | 0.995 | 0.996 | -0.001 (2/10) | |
| Norman | PC2 | 0.791 | 0.791 | +0.000 (4/10) | |
| Norman | residualized | 0.996 | 0.998 | -0.002 (2/10) | 0.519 |
| Adamson | PC1 | 0.995 | 1.000 | -0.005 (0/10) | |
| Adamson | PC2 | 0.802 | 0.793 | +0.010 (6/10) | |
| Adamson | residualized | 0.990 | 1.000 | -0.010 (0/10) | 0.141 |

Absolute performance depends on which axis is scored. The PC2 result does not establish that learning gains are invariant to target definition, because both fitted rankers retain feature-PC1 training targets. Residualization removes systematic response direction and changes representation as well as label construction; its reported estimator comparison is similarly conditional. The new target-trained and matched-candidate uncertainty controls are reported below. They retain explicit limits on representation, target construction and uncertainty.

#### Literature-fixed targets and sensitivity to gene-set definition

The targets above are still data-derived. We therefore replaced the target with a literature-fixed marker program on the three two-measurement systems, which are the systems that carry the paper's positive results and the ones whose caches retain gene names. The programs are a 29-gene interferon-gamma response set for the two Frangieh systems and a 20-gene T-cell activation set for Schmidt, both fixed before any result was computed and used only to define the target.

Interpreting this arm requires a target-definition sensitivity check. A ranking metric is computed against a top decile, so what matters is not whether the target correlates with the published axis but how its top-decile membership changes under gene-subset definitions. The curated programs have lower split-half top-decile overlap:

| system | curated: split-half score r | curated: top-decile Jaccard | published axis: top-decile Jaccard |
|---|---|---|---|
| Frangieh control-versus-IFNγ | 0.939 | 0.296 | 0.561 |
| Frangieh control-versus-co-culture | 0.912 | 0.251 | 0.492 |
| Schmidt | 0.756 | 0.345 | 0.779 |

The curated program has correlated split-half continuous scores, splitting half to half at r = 0.76 to 0.94, yet its top decile reproduces at a Jaccard of only 0.25 to 0.35, well below the 0.49 to 0.78 of the published axes. These gene-set comparisons do not establish a ceiling on achievable AUROC or identify why an estimator scores near chance. Curated and published scores correlate at r = +0.96 on Frangieh control-versus-IFNγ, but agreement of continuous scores does not ensure agreement of top-decile membership.

With that caveat, the gains and their perturbation-level intervals are:

| system | target | learning gain | 95% CI | scored candidates | drivers | verdict |
|---|---|---|---|---|---|---|
| Frangieh control-versus-IFNγ | curated | +0.043 | [−0.012, +0.104] | 94 | ~8 | null |
| Frangieh control-versus-IFNγ | published axis | +0.044 | [+0.007, +0.079] | 94 | ~8 | learning helps |
| Frangieh control-versus-co-culture | curated | +0.073 | [−0.001, +0.148] | 94 | ~10 | null |
| Frangieh control-versus-co-culture | published axis | +0.011 | [−0.034, +0.056] | 94 | ~10 | null |
| Schmidt | curated | +0.088 | [+0.010, +0.171] | 29 | ~3 | learning helps |
| Schmidt | published axis | +0.011 | [−0.047, +0.067] | 29 | ~3 | null |

These exploratory point estimates and intervals show target sensitivity. Frangieh IFN's estimated gain is similar under the two targets, but its curated-target interval is wider. This does not identify measurement noise as the cause. Schmidt's curated-target result differs from its supplied-axis result; small samples and exploratory multiplicity limit the strength of that observation. Gene-subset overlap should be reported as sensitivity to target construction, not a validated performance ceiling.

#### Interpretation limits

Splitting a curated program's genes measures sensitivity to biological feature choice, not independent replicate noise. Comparing halves of 20–29 curated genes with halves of up to 2,000 axis genes changes aggregation depth and gene content. No ceiling on AUROC follows from Jaccard alone. Independent cell, guide or donor replicates and matched gene counts would be needed to support a measurement-reliability explanation. The Schmidt curated-target gain is exploratory evidence of target dependence, subject to small-sample and multiplicity limitations, rather than a result to discard because it differs from the supplied-axis comparison.

A related endpoint question, whether inverse ranking is the more robust endpoint than forward prediction under held-out gene families, was tested and returned a null; the controlled gene-family holdout analysis is reported in Supplementary Note 8.

#### Target-definition and information-access controls

These exploratory controls were specified before model fitting. Ridge penalty is 10; there is no selection among test-performing arms. All results use ten canonical splits. Intervals are 95% percentile intervals from 1,000 paired candidate-cluster bootstrap draws, applying one weight per candidate across overlapping splits. They condition on fitted models and target axes and do not include model refitting or independent-system uncertainty. Intervals are unadjusted for the multiple systems and specifications.

##### Training on the actual PC2 outcome

For each single-condition system, features, row splits and feature/label gene halves follow the canonical protocol. The label is PC2 of the label-gene matrix fitted on all rows; its sign is chosen by its correlation with feature-PC1 scores on training rows only. Thus axis construction remains transductive, while orientation does not inspect held-out candidate outcomes. The ridge is trained on that PC2 score on training candidates. Its comparator is a supervised covariance projection estimated from the same training PC2 scores and raw feature genes. This comparator is different from the outcome-free feature-PC1 projection. A rank-target ridge and rank-target covariance reference are also both fitted.

| System | PC1-trained ridge rescored on PC2 | PC2-trained ridge | PC2 covariance | Ridge minus covariance, 95% CI |
|---|---|---|---|---|
| Frangieh | 0.585 | 0.983 | 0.989 | -0.0063 [-0.0254, +0.0063] |
| K562 | 0.705 | 0.996 | 0.995 | +0.0006 [+0.0002, +0.0009] |
| RPE1 | 0.814 | 0.998 | 0.997 | +0.0011 [+0.0005, +0.0018] |
| Norman | 0.731 | 0.999 | 0.994 | +0.0047 [+0.0012, +0.0105] |
| Adamson | 0.731 | 0.944 | 0.985 | -0.0414 [-0.1112, +0.0055] |

| System | PC2 rank-target ridge | PC2 rank-target covariance | Difference, 95% CI |
|---|---|---|---|
| Frangieh | 0.963 | 0.989 | -0.0266 [-0.0450, -0.0102] |
| K562 | 0.983 | 0.994 | -0.0117 [-0.0138, -0.0095] |
| RPE1 | 0.991 | 0.965 | +0.0256 [+0.0198, +0.0313] |
| Norman | 0.986 | 0.990 | -0.0041 [-0.0154, +0.0053] |
| Adamson | 0.929 | 0.981 | -0.0512 [-0.1252, -0.0045] |

Training for PC2 substantially raises absolute performance compared with rescoring PC1-trained rankers. The small positive ridge-minus-covariance differences on K562, RPE1 and Norman are specific to this representation, target and reference; they do not reinstate a target-invariant claim about learning. For the smaller libraries, sign choices can differ from the historical all-row orientation. The paired comparison above always scores both methods against identical labels.

#### Track B masking and disjoint anchor-row controls

Candidates are restricted to perturbed genes with a measured readout column, as in the historical Track B analysis. The canonical 60/40 row split is then applied to that restricted library. The unmasked and masked controls both fit scaling and PCA using training candidates only, isolating diagonal masking from the historical all-candidate scaling. For each training candidate, masking replaces its own-intervention entry with the mean for other fitting candidates in that coordinate, before scaling and PCA. Target-condition profiles are reported separately because they have an additional measurement budget.

| System | Source unmasked | Source masked | Masked minus unmasked, 95% CI | Target unmasked | Target masked |
|---|---|---|---|---|---|
| CD4 | 0.585 | 0.586 | +0.001 [-0.001, +0.003] | 0.575 | 0.576 |
| FrangiehIFN | 0.584 | 0.571 | -0.014 [-0.074, +0.048] | 0.509 | 0.491 |
| FrangiehCoc | 0.525 | 0.505 | -0.020 [-0.087, +0.041] | 0.561 | 0.550 |
| Schmidt | 0.545 | 0.514 | -0.030 [-0.228, +0.156] | 0.604 | 0.649 |

The disjoint anchor control uses the first half of the canonical training rows only to measure candidate readout profiles. The other half fits the model; held-out test rows are unchanged. Neither fitting nor test candidates contribute their own intervention to any anchor profile. The Track A covariance reference is refitted on exactly the same reduced fitting candidates. This is a symmetric information control, with fewer fitting examples and fewer anchor measurements than the full training-row analysis.

| System | Anchor source ridge | Anchor target ridge | Matched Track A covariance | A minus source B, 95% CI |
|---|---|---|---|---|
| CD4 | 0.575 | 0.573 | 0.834 | +0.259 [+0.234, +0.282] |
| FrangiehIFN | 0.511 | 0.482 | 0.579 | +0.068 [-0.069, +0.200] |
| FrangiehCoc | 0.516 | 0.533 | 0.517 | +0.000 [-0.109, +0.126] |
| Schmidt | 0.550 | 0.486 | 0.837 | +0.287 [+0.150, +0.425] |

The controls estimate the behavior of these ridge representations, not an upper bound on unseen-perturbation performance. Masking and anchor sampling can improve or reduce a score. Differences between Track A and Track B continue to combine information and representation/estimator changes; no causal information-budget ratio is reported. All individual split predictions, candidate indices, fit populations and bootstrap summaries are retained with the analysis code.

#### Common baseline specifications across the complete grid

The canonical rerun applies raw- and rank-target specifications to both sides of each ridge-minus-projection comparison on all seven grid configurations. Single-condition PC1 projection has no fitted outcome transform and is unchanged between columns. These are exploratory sensitivities, not prospectively chosen winners. No ratio combines a maximum selected across model classes with a different projection objective.

| System | Raw ridge | Raw projection | Raw gap | Rank ridge | Rank projection | Rank gap |
|---|---|---|---|---|---|---|
| CD4 | 0.888 | 0.834 | +0.053 | 0.884 | 0.834 | +0.050 |
| FrangiehIFN | 0.639 | 0.575 | +0.064 | 0.766 | 0.722 | +0.044 |
| Frangieh | 0.945 | 0.953 | -0.008 | 0.839 | 0.953 | -0.114 |
| K562 | 0.998 | 0.998 | +0.000 | 0.973 | 0.998 | -0.025 |
| RPE1 | 0.993 | 0.994 | -0.000 | 0.989 | 0.994 | -0.004 |
| Norman | 0.995 | 0.996 | -0.001 | 0.968 | 0.996 | -0.028 |
| Adamson | 0.995 | 1.000 | -0.005 | 0.928 | 1.000 | -0.072 |

#### Supplementary Note 8: Is inverse ranking the more robust endpoint? A tested null

The main text reports that inverse ranking accuracy is flat across test-train signature distance, the stratification on which forward prediction is reported to degrade. That invites a stronger claim, that inverse ranking is the more robust of the two endpoints, and this Note records the two experiments we ran to test it and why we do not make that claim.

##### A distance-binned comparison is not interpretable, and we report it as a failed design

The obvious test scores both endpoints on the same splits and compares their degradation from the near to the far distance tercile. We ran it on CD4+ over ten splits with an identical latent, rows and model class, varying only what is predicted. The result was uninterpretable:

| endpoint | near | mid | far | far minus near |
|---|---|---|---|---|
| inverse (AUROC) | 0.875 | 0.851 | 0.879 | +0.4% |
| forward (response correlation) | 0.221 | 0.270 | 0.356 | +60.7% |
| forward, mean-centred | 0.218 | 0.261 | 0.344 | +57.7% |

Forward accuracy rises by 61% with distance from the training set, the opposite of the inter-heterogeneity finding the design was meant to reproduce. This does not invalidate prior cross-context studies: distance within one atlas is a different generalization axis. We examined two possible confounds. The far tercile carries 1.18-fold larger late responses, and a response correlation is far easier to score on a large structured response than on a near-null one, so response magnitude may contribute to endpoint difficulty. Stratifying by response magnitude does not remove the trend (+0.060, +0.040 and +0.077 within low, mid and high magnitude strata), so effect size is a real confound but not the whole story.

The general lesson is the one that matters: a distance binning does not hold difficulty fixed across two endpoints scored by different metrics, so a near-to-far contrast between them cannot be read as one degrading and the other not. We report this because the failed design is informative, and because the same trap is available to anyone repeating the analysis.

##### A gene-family holdout, with explicit composition limitations

We replaced the distance axis with a held-out structure defined outside the data. Whole Gene Ontology biological-process families are withheld from training, so membership is fixed by curators with no reference to the responses being scored but can still correlate with effect size and other aspects of difficulty. Families contain 40 to 400 perturbed genes and are greedily deduplicated so no two share more than 25% of their members, since an overlapping term would leak the held-out biology back through training. Each family is paired with a size-matched random holdout, because withholding a family removes both biology and training data while the random control removes only training data; the difference combines training-set exclusion and evaluation-composition effects, because the family and random controls score different perturbations. Both endpoints use the identical fold, latent, rows and model class.

| GO term | process | held out | inverse, family vs random | forward, family vs random | magnitude ratio |
|---|---|---|---|---|---|
| GO:0016567 | protein ubiquitination | 352 | 0.931 vs 0.888 (+0.043) | 0.296 vs 0.276 (+0.020) | 1.01 |
| GO:0045892 | negative regulation of transcription | 347 | 0.848 vs 0.898 (-0.051) | 0.294 vs 0.280 (+0.014) | 1.04 |
| GO:0035556 | intracellular signal transduction | 267 | 0.952 vs 0.879 (+0.073) | 0.304 vs 0.288 (+0.016) | 1.00 |
| GO:0043066 | negative regulation of apoptosis | 239 | 0.863 vs 0.894 (-0.031) | 0.293 vs 0.298 (-0.005) | 1.02 |
| GO:0006338 | chromatin remodeling | 218 | 0.786 vs 0.888 (-0.101) | 0.300 vs 0.282 (+0.018) | 1.08 |
| GO:0006974 | DNA damage response | 217 | 0.817 vs 0.877 (-0.059) | 0.298 vs 0.272 (+0.026) | 1.05 |
| GO:0008284 | positive regulation of cell proliferation | 190 | 0.886 vs 0.904 (-0.017) | 0.291 vs 0.292 (-0.002) | 1.00 |
| GO:0055085 | transmembrane transport | 190 | 0.728 vs 0.904 (-0.176) | 0.269 vs 0.292 (-0.023) | 0.96 |
| GO:0006886 | intracellular protein transport | 183 | 0.865 vs 0.882 (-0.017) | 0.259 vs 0.297 (-0.038) | 0.98 |
| GO:0043065 | positive regulation of apoptosis | 174 | 0.926 vs 0.874 (+0.051) | 0.284 vs 0.285 (-0.001) | 0.99 |
| GO:0006955 | immune response | 169 | 0.929 vs 0.848 (+0.081) | 0.288 vs 0.273 (+0.015) | 0.99 |
| GO:0008285 | negative regulation of cell proliferation | 165 | 0.832 vs 0.904 (-0.071) | 0.290 vs 0.286 (+0.004) | 1.01 |

The mean response-norm ratios are close, but matching means does not establish equal response distributions, driver prevalence or task difficulty. Family folds share most training data and can share genes; they are not independent biological replications. The script selects families by size and overlap and skips folds with fewer than three drivers, which limits the reported population.

#### No established endpoint advantage, with family-dependent changes

Across the twelve reported families, inverse ranking has mean family-minus-random difference −0.023 AUROC and forward response correlation has mean +0.004. Individual ranking losses are substantial in some families, including −0.176 for transmembrane transport and −0.101 for chromatin remodeling. A standard deviation larger than the mean is not evidence that these differences are noise.

The exploratory Wilcoxon comparison of relative changes has $p = 0.27$. It does not establish either endpoint's equivalence or stability. AUROC and response correlation have different reference levels and scales, so dividing by their respective random-control scores does not create a common effect scale. We retain that calculation as exploratory provenance, not a definitive endpoint-comparison test. No prespecified noninferiority margin was used.

We report heterogeneous family behavior and no detected average asymmetry under this analysis. We do not claim that inverse ranking is uniformly stable or more robust than forward prediction. A stronger design would evaluate the same family under size-matched training exclusions, quantify family-specific uncertainty and define practical tolerances separately for each endpoint.

The null must not be read as contradicting the forward-prediction benchmarks we cite. Those measure degradation across cellular contexts, holding out cell types a model never saw. Our holdout withholds gene families within a single cell type. Those are different axes of generalization, and a cell-type holdout is untested here, so our result bears on novel biology within a context rather than on transfer between contexts.

One scope limit applies to the null as much as it would have to a positive result. This is one atlas and one family definition. A donor holdout, which a reader may reasonably prefer, is not available to us: the differential-expression atlas every published number here is computed from is already pooled across the three donors and retains only donor correlation summaries, so recovering per-donor effects would mean re-deriving differential expression from roughly 1.4 TB of raw files and would yield an atlas no longer matching the one the rest of the paper uses.

### Supplementary Note 9 | Rerun validation and remaining numerical uncertainty

#### Shared-space numerical sensitivity

All 70 system/split combinations were rerun after preserving the historical feature-matrix memory layout during the single-condition PCA fit. Fancy indexing had converted the original Fortran-contiguous gene slice to a C-order copy, changing float32 PCA roundoff. The corrected implementation reproduces the historical preprocessing expressions without changing the gene split, held-out candidate rows or fitting populations. This correction addresses one source of numerical sensitivity; it does not identify the cause of every historical discrepancy.

The maximum absolute difference in system/method mean AUROC from historical scores is 0.0138 (Adamson PAIRING), compared with 0.0191 in the earlier frozen-version rerun. Frangieh PAIRING differs by -0.0074. Every label vector and retained historical row identity was checked. Independent repeats of all ten Adamson and all ten Frangieh splits reproduced the corrected PAIRING, MLP, scGen-style and gradient-boosting score arrays exactly under the recorded environment and thread settings. This is observed repeatability of the current workflow, not recovery of historical neural score arrays or a cross-platform determinism guarantee. Reference Table N10a retains the reference comparison; complete rerun score files are provided rather than selectively replacing methods.

#### Native-tool reruns with matched-candidate references

Thirty native CellOT fits, ten released GEARS fits on Norman and two Geneformer raw-cell re-embedding runs (ten scoring splits each) were completed. CellOT uses the canonical split; GEARS verifies its split against both the canonical evaluator and historical dump. GEARS training is explicitly seeded and uses twenty epochs on an H100 with CUDA-enabled PyTorch 2.3.1. Geneformer reuses the pretrained V1-10M checkpoint; no foundation-model pretraining or fine-tuning is claimed. Raw cells were converted and tokenized afresh.

The following means use identical covered candidates for each native method and its two references. GEARS coverage is restricted to predictable candidates; its references are rescored on that same subset. These controls address the former candidate-coverage mismatch but do not equalize representation, supervision, validation or tuning budgets.

| System | Native method | Rerun AUROC | Ridge, same candidates | Projection, same candidates |
|---|---|---|---|---|
| Adamson | CellOT | 1.0000 | 0.9950 | 1.0000 |
| CD4 | CellOT | 0.8358 | 0.8876 | 0.8342 |
| Frangieh | Geneformer | 0.9011 | 0.9447 | 0.9528 |
| Norman | CellOT | 0.9881 | 0.9949 | 0.9959 |
| Norman | GEARS | 0.9178 | 0.9949 | 0.9959 |
| Norman | Geneformer | 0.9907 | 0.9949 | 0.9959 |

The historical native CellOT Norman mean was 0.997; the rerun is 0.9881. Historical native GEARS was 0.922; the rerun is 0.9178. These differences are retained explicitly. Native CellOT reaches 1.000 on Adamson and therefore exceeds the continuous-target ridge mean there; the projection also reaches 1.000. The results do not establish universal linear superiority or a capacity theorem. Full candidate-level outputs, split-specific coverage and paired candidate-cluster bootstrap intervals are retained with the analysis code. The exploratory intervals use 1,000 shared candidate-weight replicates across overlapping splits, conditional on fitted scorers, without model refitting or multiplicity adjustment.

##### Full raw-atlas reconstruction and count-based sensitivity

The complete GSE123139 raw archive was reconstructed from 204 plates: 78,336 input cells, 65,035 cells and 33,456 genes after QC, with 2,000 highly variable genes. The thirty-dimensional scVI model completed all 200 training epochs on CPU using scvi-tools 1.1.3 and seed zero; finite latent values and the saved model were verified. Cell identities and order match the historical atlas. The expression-derived dysfunction coordinate has Spearman correlation 1.000 with the historical coordinate. This coordinate is calculated from marker expression independently of scVI, so its agreement is not evidence of identical latent embeddings or temporal ordering.

The archived sensitivity script incorrectly interpreted A.raw.X as integer counts; inspection showed that field is log-normalized in the saved atlas. Main-model training already used the correct layers['counts']. The sensitivity script now uses that integer-count layer. Six scVI fits on the same 12,000-cell subsample, each with up to sixty epochs, yield absolute coordinate Spearman correlations against the thirty-dimensional seed-zero reference of 0.9352 (ten dimensions), 0.9503 (twenty), 0.9497 (fifty), 0.9450 (seed one) and 0.9387 (seed two). These are corrected results and supersede the archived count-misinterpreted sensitivity run. Sensitivity coordinates project scVI onto an expression-defined high-minus-low contrast.

#### Supplementary Note 10 | Matched information contracts and model selection

This note documents the InverseBench evaluation protocol and the matched information contracts used throughout the paper. FSTN is the reference pipeline evaluated under it; its response model is the continuous-target ridge of the main text and its combination step is Algorithm 1.

The seven-task design fixes common eligible gene candidates, disjoint anchor interventions and training-only selection. All performance and uncertainty tables are derived from retained candidate-level predictions.

Table 1. Tasks, eligible gene interventions and endpoint provenance. Seven tasks represent six study groups. Counts are after shared candidate eligibility, not independent biological sample counts.

| Task | Genes | Destination | Scope |
|---|---|---|---|
| CD4 activation | 8,326 | Supplied 48 h axis | Same atlas; candidate-conditional inference |
| Frangieh IFN | 226 | Supplied IFN axis | Shared Frangieh cohort |
| Frangieh co-culture | 225 | Supplied co-culture axis | Shared Frangieh cohort |
| Schmidt activation | 72 | Supplied activation axis | Small candidate library |
| Song fixed markers | 1,883 | CD69/IL2RA expression | At least 10 cells per arm |
| Papalexi PD-L1 | 24 | Surface PD-L1 protein | 24 genes; technical replicates |
| K562 external fitness | 7,354 | Negative Chronos gene effect | Same cell line; independent assay and modality |

Table 2. Controlled response-entry availability under the fixed-dimension ridge pipeline. Entries are mean test Spearman correlations over ten outer splits. The final column is the paired full-

minus-zero availability effect with a gene-cluster 95% interval, conditional on fixed fitted models. All tasks, including weak and uncertain contrasts, are retained.

| Task | 0% | 10% | 50% | 100% | Full – zero [95% interval] |
|---|---|---|---|---|---|
| CD4 activation | 0.064 | 0.289 | 0.467 | 0.506 | 0.442 [0.419, 0.465] |
| Frangieh IFN | 0.044 | 0.049 | 0.193 | 0.165 | 0.121 [0.002, 0.236] |
| Frangieh co-culture | 0.074 | 0.076 | 0.069 | 0.124 | 0.050 [-0.052, 0.148] |
| Schmidt activation | -0.069 | 0.237 | 0.628 | 0.598 | 0.667 [0.464, 0.814] |
| Song fixed markers | 0.001 | -0.004 | 0.025 | 0.026 | 0.026 [-0.009, 0.062] |
| Papalexi PD-L1 | -0.004 | 0.130 | 0.370 | 0.358 | 0.361 [-0.145, 0.706] |
| K562 external fitness | 0.377 | 0.470 | 0.487 | 0.494 | 0.116 [0.101, 0.133] |

#### Reference benchmark and application methods

#### Methods

Datasets and preprocessing. The primary system is human melanoma tumor-infiltrating lymphocytes (GSE123139, Li et al. [33]), profiled by MARS-seq; we used the raw count matrix and retained 65,035 of 78,336 loaded cells, the cell-level quality-control criterion being a minimum of 200 detected genes per cell (Extended Data Fig. 5a), with the sequencing plate (204 plates) retained as a batch covariate. Genes detected in fewer than three cells were removed, leaving 33,456 of 55,857 genes, and counts were kept unnormalized for the negative-binomial embedding (below). For cross-tumor generalization we used GSE123813 (Yost et al. [34]), which provides matched 10x scRNA-seq and T-cell-receptor sequencing for basal and squamous cell carcinoma, and GSE218262 for non-small-cell lung carcinoma; each was processed with the same QC and restricted to T cells. The genome-scale perturbation ground truth is a CRISPRi Perturb-seq atlas of primary human CD4+ T cells ([20]; GSE314342): we used the per-perturbation differential-expression signatures provided in the zscore layer, comprising 33,983 perturbation-by-condition profiles across 11,526 perturbed genes measured in three culture conditions (resting, 8 h and 48 h stimulation), and restricted analyses to single-gene perturbations with sufficient cells per the resource's inclusion criteria. CD8+ validation used pooled CRISPR transcription-factor screens in human CD8+ T cells (GSE241933; the related superseries GSE218988 was retrieved but not used in any reported analysis), building on genome-wide CRISPR screens that map regulators of primary T-cell function [35], from which we computed a per-gene phenotype as the mean log2 fold change of a gene's guides between the sorted high and low populations. The second, independent transition is K562 CRISPRa Perturb-seq (GSE133344, Norman et al.), for which we computed per-perturbation expression signatures and an erythroid-differentiation score; the negative control is a dendritic-cell LPS-activation Perturb-seq (GSE90063, Dixit et al. [36]). For the multi-system fair benchmark we additionally used two genome-wide Replogle CRISPRi Perturb-seq screens [29]: K562 (11,258 single-gene perturbations) and RPE1 (2,679 perturbations); a melanoma CRISPR-KO Perturb-seq of the IFN response (Frangieh et al. [37], 246 perturbations), which we analyse both as a single-condition system and, exploiting its control/IFNγ/co-culture design, as a two-condition system (features from each perturbation's control-condition signature, labels from its IFNγ-condition effect; 235 perturbations have at least 20 cells in both conditions) that provides a second cross-condition map like CD4+; and a K562 CRISPRi Perturb-seq of the unfolded-protein response (Adamson et al. [38], 107 perturbations). Single-condition labels are defined by PC1 of held-out label genes. Two-measurement labels use the supplied target-condition axis and no gene split. Both families hold out perturbation rows; their preprocessing populations differ as specified below. Where a harmonized release was available we used it [39]. All accessions and URLs are listed in the bibliography.

M1, expression coordinate and embedding. The melanoma coordinate is the min-max scaled expression-derived dysfunction/reactivity score computed with scanpy score_genes. It is not a projection of the scVI embedding. The scVI embedding uses the count layer, plate batch covariate, 2,000 variable genes, 30 latent dimensions and 200 epochs. The independently implemented latent-projection sensitivities answer a different question; agreement of the expression coordinate with its own defining score is a consistency check, not independent validation.

M2, transition-ordered regulators. For each candidate regulator we computed the Spearman correlation of its expression (or, where applicable, its inferred activity) with the transition coordinate, giving a signed association with the reactive pole; the full ranking is Supplementary Table 3. We additionally estimated an activation-order statistic as the coordinate value at which a regulator's expression crosses its upper quartile, yielding a coordinate-crossing order, not a measurement of temporal activation. Recovery was scored as recall of a literature-curated tumor-reactivity transcription-factor set (Supplementary Table 2) among the transcription factors ranked by their association with the transition coordinate, over a standard human transcription-factor catalog (1,731 factors present in the data; SCENIC allTFs union the curated set); the curated set was used to report recall rather than to fit the response ranker; transcriptional overlap with the axis-defining programs limits its independence. Ten of the fifteen curated factors fell within the top fifty (recall@50 = 0.67), with the exhaustion and reactivity regulators TOX, EOMES and ID2 the three top-ranked factors of the entire catalog; the un-recovered factors (including TCF7 and ZEB2) mark the opposite naive and memory pole and rank accordingly (Extended Data Fig. 6c).

M3, part 1: response model (predictive and mechanistic variants). M3 has two components, a response model that scores single edits and a transport layer that selects a set from them; this subsection covers the first and the next the second. The core learner maps a perturbation's transcriptional signature to its effect on the transition. In the predictive variant we reduced each perturbation's early (8 h) or resting signature to 50 principal components and fit ridge regression [40] (L2 penalty alpha = 10) to the perturbation's effector-axis effect, defined as the projection of its late (48 h) differential-expression signature onto the effector axis. Regression parameters were fitted on training perturbations under a 60/40 split. Early features and late outcomes come from different conditions. The supplied condition-contrast axis is fixed by the loader; the release manifest distinguishes that external axis from per-split fitted model parameters. The mechanistic variant instead learns a reduced-rank linear response operator (ridge, alpha = 50, 100 components on both input and output) that maps a perturbation's early signature to its full downstream expression shift, from which the effector effect is read by projection; the operator is fitted to measured interventions, but interpretation as causal transport across contexts requires additional assumptions [41]. It achieved held-out accuracy comparable to the predictive variant under the evaluated settings.

M3, part 2: nonnegative response-set selection. Given a response dictionary and a target mean displacement, the selector greedily adds candidate directions and refits nonnegative coefficients by NNLS [42]. It uses a PCA representation of measured or predicted response signatures, greedy decorrelation (cosine threshold 0.7, at most 80 representatives), and an 0.85 displacement-coverage target. The auxiliary entropically regularized Wasserstein readout uses Sinkhorn regularization epsilon = 0.5 [43,44]; it does not make the mean-displacement selector an optimal-transport solver. Algorithm 1 selects only positively aligned responses, since an anti-aligned response cannot be used by the nonnegative refit, and is stated in the Methods. Greedy recovery is not a general proof of minimum cardinality or submodularity; Extended Data Fig. 7 illustrates behavior on one synthetic problem. Algorithm 1 is evaluated directly on the Norman constituent-retrieval task; a signed matching-pursuit variant without joint refitting is retained there as the comparison (Supplementary Note 3).

Complexity. Ridge fitting in K shared dimensions costs $O(PK^2 + K^3)$, with $O(CK)$ scoring for C candidates. Greedy selection scans the dictionary and solves restricted NNLS problems repeatedly; its runtime depends on the active-set solver and selected set size. These downstream stages can run on CPU. This excludes scVI training, PCA and data preparation, and does not describe neural comparator training or native GEARS, which used a GPU. We do not infer an end-to-end runtime advantage without a matched timing experiment.

Deployment scenario and information budget. Track A provides every candidate's measured source-condition signature, or feature-gene signature subset, before ranking. The source-to-target task is response-informed prioritization, not fully unseen intervention nomination. Track B describes candidate genes by co-response columns of training perturbations. A target-condition co-response variant is allowed and reported explicitly. In the historical Track B arm a held-out candidate contributes no own intervention row, but training candidates do. Masking and disjoint-anchor controls now test that asymmetry; the anchor arm excludes both fitting and test candidates from the profile-generating intervention rows. Neither track validates functional conversion in an unmeasured tumor. Supplementary Note 7 records the tested estimators, candidate coverage and target-definition sensitivities.

Shared-space benchmark and information separation. Eight learned model classes are evaluated: continuous-target ridge, CellOT-style neural transport, a PAIRING-style VAE adaptation, a scGen-style latent-shift model [45], a GEARS-style GNN, an MLP, kernel ridge and gradient boosting. Shared-space adaptations are distinguished from released native tools. Single-condition systems use a 60/40 feature/label gene split, axes fitted over all perturbation rows, and a 60/40 perturbation-row split. PCA and feature scaling use all feature rows. The target axis is signed to agree with the feature axis over all rows, so preprocessing is transductive. In two-measurement systems the source signature is the feature and the target-condition outcome supplies the label. There is no gene split. PCA fits on training perturbations, feature scaling uses all feature rows, and source/target clouds are chosen from training outcomes. Learned methods fit on training rows and score held-out rows in both families. Driver labels are the top decile of the defined outcome across the candidate pool. The registry exposes a label-free method context; evaluator labels and split records remain outside it. This contract is not a security sandbox for arbitrary submitted Python. Reference-ranking reproduction and per-family manifests are described in Supplementary Note 6. The generic and registry paths delegate split construction to the canonical module; legacy analysis scripts must be checked against that protocol before adding results.

Metrics and paired comparisons. AUROC is the primary held-out-candidate metric. AUPRC, precision@25, precision@100 and Spearman score correlation are reported for the stored configurations in Supplementary Table 4, without extrapolating uncomputed metrics to the rank-target sensitivity analysis. Paired AUROC intervals resample perturbation identities jointly across their repeated held-out occurrences when identities can be reconstructed and verified. This conditions on fitted models. Single-condition and GNN summaries use resampled split-level paired differences where candidate-level information is not available to that procedure; overlapping splits limit their interpretation as independent uncertainty estimates. Published multiple-testing summaries cover the stated FSTN-versus-comparator family rather than every exploratory analysis, baseline specification and target construction. We distinguish stored adjusted p-values from the newer candidate-cluster intervals. No equivalence or noninferiority margin was specified, so intervals spanning zero are not labeled equivalence.

Systematic-variation diagnostic. Following the concern raised by Systema [9], axis alignment is the cosine between the target direction and the mean uncentered perturbation-response vector. Supplementary Table 7 and Extended Data Fig. 8 report it descriptively. It does not by itself establish whether ranking is biologically specific or whether a supervised model can improve on projection. Feature-PC1 reconstruction, systematic-response alignment and cross-condition predictivity are different properties.

Cross-condition and target sensitivities. We compare multiple condition pairs from the CD4+ atlas, Frangieh control-to-IFN-gamma/co-culture and Schmidt rest-to-restimulation. These configurations differ in sample size, noise and outcome construction. Additional pairs within an atlas are not independent cohorts. Ridge trained on rank-transformed outcomes is an exploratory specification check, applied to both regression and covariance-projection estimates and reported alongside the original continuous-target estimates. The separate synthetic Gaussian illustration concerns covariance geometry and does not establish a necessary condition based on measurement count. Alternative-target and residualization scripts are described with their actual training targets in Supplementary Note 7.

Inter-heterogeneity (out-of-distribution) stratification. To test robustness to test-train dissimilarity, we defined each held-out CD4+ perturbation's inter-heterogeneity as the minimum Euclidean distance in the shared latent from that perturbation to the nearest training perturbation, and partitioned held-out perturbations into near, mid and far terciles by this distance, reporting FSTN's ranking AUROC within each tercile (Extended Data Fig. 2).

Predicted-effect selection test. For fixed budgets k, response-set selection and top-k ranking are evaluated using additive held-out response vectors. Predicted effects have mean predicted-versus-measured cosine 0.594 across ten splits. Closure measures source-to-target mean-axis displacement under this simulated additive dose model; it is not measured survival, function, full-distribution matching or actual cell conversion. The measured-effect regime is an oracle analysis. The conclusions apply to these predictors and selectors, not all possible combination strategies.

Head-to-head implementations. CellOT was run as its own neural optimal-transport model, using the released input-convex neural-network architecture [46] and an adversarial optimal-transport objective [47,48]: two input-convex networks (potentials f and g) with two hidden layers of 64 units, LeakyReLU activations and softplus-constrained weights, trained with Adam (learning rate 1e-3, betas 0.5/0.9) for 2,000 outer iterations with 10 inner updates of g per outer step and batch size 128, to transport the non-reactive (source) latent distribution onto the reactive (target) distribution; candidate perturbations were then ranked, in the shared latent space, by the alignment of their held-out early signature with the learned transport direction (the mean displacement under g.transport). PAIRING was adapted from its published mechanism, a variational autoencoder with adversarial basal/perturbation disentanglement and latent transition-vector alignment, rather than run from the authors' release, because that release operates in a fixed 978-gene landmark space with a small-molecule and shRNA perturbation vocabulary that would confound a comparison on our CRISPRi gene set; the encoder and decoder used 64-unit hidden layers, the latent was split into basal (8) and perturbation (8) components, a discriminator enforced realistic reconstructions and an adversary was trained to predict the source/target group from the basal latent while the encoder was trained so it could not (isolating the transition in the perturbation latent), optimized with Adam (1e-3) for 2,500 steps; candidates were ranked by cosine alignment of their held-out early perturbation-latent with the source-to-target transition vector. The deep GEARS-style graph neural network was a model-class adaptation of the GEARS mechanism [49] evaluated in the shared latent: a k-nearest-neighbour graph whose nodes are perturbation candidates in the shared latent, with neighbours searched among training candidates and test nodes attached by bidirectional edges, followed by a two-layer graph convolutional network (torch_geometric) and an MLP head, trained for 150 epochs on the training-row (latent to effect) pairs, and scored on held-out perturbation rows in the same shared latent. We used a graph-model class adaptation, a shared-latent CellOT implementation and a PAIRING-style adaptation because the benchmark's shared-representation condition requires every method to rank in the same neutral latent; the native pipelines each operate in their own feature space, which would reintroduce the feature-space asymmetry the benchmark controls for. The releases confirm this is not merely a convenience: the released CellOT and GEARS install and run (GEARS only under a specific dependency-pinning recipe), each in its own gene-graph or transport feature space, while the released PAIRING is locked to a 978-gene L1000 landmark space and a fixed 8,532-entry small-molecule/shRNA/CRISPR perturbation catalog and ships no training data, so its provided configuration does not directly implement our candidate universe without additional input mapping and adaptation. As a robustness check that the shared latent does not itself handicap the deep methods, we additionally ran the released CellOT code in its own native gene feature space (full train-gene signatures rather than the shared PCA-30 latent), keeping the gene split, held-out-row split and driver labels byte-for-byte identical to the benchmark (asserted against the stored label dumps before scoring); it scored AUROC 0.837 +/- 0.015 on CD4+ (below linear FSTN's 0.888 and at the additive projection's 0.834) and tied the additive ceiling on the saturated single-condition systems (Norman 0.997, Adamson 1.000); the released GEARS graph network, trained for 20 epochs on a GPU in its native gene space, likewise scored 0.922 +/- 0.039 on Norman, below linear FSTN (0.995) and the additive projection (0.996), these are limited native-space checks, not a representation-independent capacity theorem (Supplementary Note 5). The shared-space GNN was run only on the three systems with at least 500 perturbations, and its sign convention was aligned to the training axis by correlation (identical to the labelling axis, so no independent orientation heuristic could flip it); a shuffled-target control returned AUROC near 0.5 on every split, providing a negative control, without proving that every possible information leak is absent. Four additional learned classes were fit in the identical shared latent on training rows only and scored on held-out rows: a deep multilayer perceptron (two 64-unit ReLU hidden layers, Adam, 300 epochs) to test whether depth rather than graph structure helps; a kernel-ridge regressor (RBF kernel, alpha and gamma chosen by 3-fold cross-validation on training rows) and a gradient-boosting regressor (histogram-based, 200 trees, depth 3) to test whether nonlinearity helps; and a scGen-style generative model, a variational autoencoder over the shared latent from which a perturbation vector is formed as the latent difference between low- and high-effect training rows, and candidates scored by projection onto it, an adaptation of scGen's latent-arithmetic mechanism (the released scGen operates in its own gene space). Each of the four passed a shuffled-target leakage control (AUROC ~0.5) and produced scores distinct from every other method. CellOracle was run in its official container (kenjikamimoto126/celloracle_ubuntu:0.18.0) via apptainer [50] using its built-in human promoter base gene-regulatory network, and scored on the transcription factors present in its inferred network. Dynamo was run in an isolated environment at documented defaults and scored on highly variable genes. All baselines were recomputed on the same held-out splits rather than quoted from their source papers (Extended Data Fig. 9). The projection reference is family-specific: feature-PC projection for single-condition tasks and a raw-feature covariance projection estimated from training target outcomes for cross-condition tasks. The latter uses supervision; it is not the same estimator as an outcome-free additive response baseline [7]. Each published tool was run in its own environment on identical inputs and the same metric and scored on its native candidate coverage, which we report explicitly rather than pooling across a tool's non-overlap.

Generalization and negative control. For the second transition (K562, Norman) we scored erythroid differentiation from a canonical marker set, computed per-perturbation effects, fit the response model on training perturbations, and validated on held-out perturbations with shuffled-target and random-gene-set controls to exclude circularity. For the negative control (dendritic-cell LPS activation, Dixit) we applied the identical resting-to-activated two-condition design and report the honest null. Generalization within melanoma was tested by learning the reactive-regulator ranking on one disjoint set of plates and measuring rank concordance on the held-out plates, and across tumors by transferring the melanoma reactive-minus-naive signature to independent skin and lung tumors; both used shuffled-label permutation nulls.

Orthogonal validations. We validated the reactive regulators in the cytotoxic lineage using CD8+ screens (Mann-Whitney comparison of the recovered transcription factors' screen phenotypes against other genes; Extended Data Fig. 6b), and assessed the reactive axis against an orthogonal clonal-expansion readout, T-cell-receptor clonal expansion, testing whether the axis separates expanded clones from singletons (AUROC 0.72; Extended Data Fig. 6a). A same-condition circular fit, which yields a near-perfect but invalid result, was excluded and is shown explicitly as a leakage control (Extended Data Fig. 10).

Uncertainty and computation scope. We report means and standard deviations across ten evaluation splits, permutation controls and paired comparisons. Two interval estimators are used and the choice is determined by whether a split's candidate identities can be recovered and verified, not by the result. Candidate-cluster intervals, which resample candidates across all their occurrences and so preserve repeated-candidate dependence, are available for five systems: CD4+ activation, the Frangieh IFN and co-culture two-condition analyses, Schmidt and Song. The remaining seven, the Frangieh single-condition analysis, K562, RPE1, Norman, Adamson and the two CD4+ condition pairs, have split-level resampling only, because their candidate identities could not be reconstructed and verified against the stored per-candidate scores. Identity-verification status is recorded for each comparison in the accompanying analysis records. Candidate-cluster intervals condition on the trained models; split resampling resamples overlapping splits, is not guaranteed conservative and cannot supply donor-level inference. Historical GNN intervals use per-split AUROC summaries. The new canonical rerun retains candidate indices and GNN scores. Its bidirectional attachments permit test-node features to affect training-node messages, so graph structure is transductive even though only training outcomes are supervised. CD4+ and CD8+ validations are separate. CPU-only claims apply to downstream linear ranking and NNLS selection, not scVI, neural comparator fitting or native

GPU GEARS. Supplementary Table 6 records the frozen environment, whose platform-specific dependencies may require adjustment for a new compute host.

Hyperparameters and settings. The original reported configuration uses 30 shared latent dimensions and ridge alpha = 10; demonstration response variants use 50 or 100 components and the stated penalties. Comparator settings are documented above. KRR selects alpha and gamma by training-only cross-validation. Rank-target fitting and penalty sweeps are exploratory analyses motivated by observed results. They were not specified a priori. Competitors were not all tuned under a matched objective-selection budget, which limits conclusions about model-class capacity. Embedding and seed sensitivities are reported as robustness checks rather than evidence that every hyperparameter choice is optimal.

Rerun validation. All seventy shared-space splits and sixty native-tool evaluations were rerun. Preserving the historical PCA input layout reduces the largest mean AUROC discrepancy from 0.0191 to 0.0138; exact historical neural reproduction remains unresolved. Twenty independent repeat splits reproduce current score arrays exactly under the recorded settings. Full raw-atlas reconstruction and scVI training were completed, and count-based embedding sensitivities were corrected (Supplementary Note 9). Reference Table N10a retains the reference-computation values.

Code and data availability. The software is available from the corresponding author on request, and will be released publicly with an archival DOI before publication. Four main-grid processed signature caches and two additional boundary-control caches accompany the standalone Python package, with source-data provenance and checksums. The full seven-system run additionally requires the external CD4 cache and Replogle matrices documented in the release instructions. The release includes the canonical protocol, method interface, benchmark source, control analyses and retained validation records; arbitrary Python methods must honor the declared information contract. Public archival deposition is pending and no archival DOI has been assigned. A clean-environment four-system harness test is documented; the completed native-tool and raw-atlas reruns are documented in Supplementary Note 9; complete clean-host reproduction of every historical figure and external tool is not claimed. All primary datasets and accessions are listed in Supplementary Table 1.

---

#### Reference benchmark tables

Reference Table N10a. Inverse-ranking benchmark across eight learned model classes and six transition systems. Seven columns represent six systems because Frangieh has both single- and two-condition configurations. Values are mean AUROC ± s.d. over ten held-out-perturbation splits where s.d. was retained. Single-condition labels use held-out genes; cross-condition labels use a separate measurement without a gene split. Projection references are fixed feature-PC projections in single-condition tasks and supervised covariance projections in cross-condition tasks. Rank-target rows are sensitivity analyses, not a prospectively selected estimator.

| Model class | CD4+ activation (Marson) | melanoma IFN 2-condition (Frangieh) | melanoma IFN (Frangieh) | K562 essential (Replogle) | RPE1 essential (Replogle) | K562 differentiation (Norman) | K562 UPR (Adamson) |
|---|---|---|---|---|---|---|---|
| FSTN, linear (ours) | 0.888 ± 0.010 | 0.639 ± 0.060 | 0.945 ± 0.032 | 0.998 ± 0.001 | 0.993 ± 0.004 | 0.995 ± 0.007 | 0.995 ± 0.008 |
| FSTN, rank-matched target (ours) | 0.884 | 0.766 | 0.839 | 0.973 | 0.989 | 0.968 | 0.928 |
| Projection reference | 0.834 ± 0.015 | 0.575 ± 0.045 | 0.953 ± 0.031 | 0.998 ± 0.001 | 0.993 ± 0.004 | 0.996 ± 0.006 | 1.000 ± 0.000 |
| Projection, rank-target covariance | 0.834 | 0.722 | 0.953 | 0.998 | 0.993 | 0.996 | 1.000 |
| CellOT (neural OT) | 0.882 ± 0.010 | 0.761 ± 0.042 | 0.902 ± 0.019 | 0.981 ± 0.006 | 0.959 ± 0.012 | 0.924 ± 0.029 | 0.955 ± 0.053 |
| PAIRING (disentangled-latent) | 0.845 ± 0.015 | 0.722 ± 0.047 | 0.817 ± 0.079 | 0.970 ± 0.004 | 0.918 ± 0.021 | 0.724 ± 0.070 | 0.782 ± 0.071 |
| scGen-style (generative) | 0.764 ± 0.032 | 0.725 ± 0.037 | 0.763 ± 0.067 | 0.945 ± 0.022 | 0.889 ± 0.034 | 0.790 ± 0.110 | 0.817 ± 0.138 |
| Kernel ridge (RBF, nonlinear) | 0.893 ± 0.010 | 0.615 ± 0.081 | 0.935 ± 0.043 | 0.998 ± 0.001 | 0.990 ± 0.005 | 0.952 ± 0.011 | 0.973 ± 0.019 |
| Gradient boosting (nonlinear) | 0.883 ± 0.009 | 0.546 ± 0.092 | 0.918 ± 0.040 | 0.998 ± 0.001 | 0.993 ± 0.004 | 0.983 ± 0.015 | 0.945 ± 0.032 |
| MLP (deep, non-graph) | 0.869 ± 0.008 | 0.565 ± 0.066 | 0.913 ± 0.051 | 0.997 ± 0.001 | 0.989 ± 0.003 | 0.960 ± 0.028 | 0.925 ± 0.045 |
| GEARS-style GNN (deep) | 0.830 ± 0.013 | — | — | 0.978 ± 0.005 | 0.952 ± 0.007 | — | — |
| Label-informed centroid contrast | 0.887 ± 0.009 | 0.759 ± 0.042 | 0.900 ± 0.018 | 0.990 ± 0.004 | 0.972 ± 0.006 | 0.934 ± 0.029 | 0.952 ± 0.059 |
| axis-alignment (systematic-variation diagnostic) | 0.58 | 0.22 | 0.44 | 0.82 | 0.98 | 0.88 | 0.51 |

Continuous-target FSTN leads the tested shared-space deep references in six columns, except the Frangieh cross-condition column where CellOT reaches 0.761 versus 0.639. A rank-target linear reference reaches 0.766 there; no equivalence test was performed. GNN results were paired over the same ten splits; the gaps are +0.058, +0.021 and +0.042, every interval excluding zero and FSTN ahead in 10 of 10 splits. These split-resampled intervals are conditional summaries of overlapping splits. The alignment row reports cosine with the mean perturbation response, not an independent guarantee of task specificity.

Reference Table N10b. CD4+ estimator ladder, mean AUROC over ten held-out-perturbation splits.

| Configuration | AUROC |
|---|---|
| Shared latent + continuous-target ridge | 0.888 |
| Shared latent + label-informed centroid contrast | 0.887 |
| Raw-feature covariance projection | 0.834 |

The centroid contrast selects source and target clouds using training outcome labels. Its proximity to ridge shows that regression adds little under this configuration; it does not show that outcome supervision is unnecessary.

#### Complete matched comparison results

The fixed-dimension core suite evaluates five families in the B-only, A-only and combined full-response arms. Every family has six settings: three structural/regularization options crossed with raw versus rank-transformed training targets. The full-profile suite evaluates five B families and three A and B-plus-prior families (ridge, histogram boosting and MLP). Within-arm family selection therefore compares 30 settings for B and 18 for A or B-plus-prior in that suite; it is not an equal search budget between those selected arms. Intermediate availability fractions use ridge alone with six settings each. The controlled availability contrast, rather than selected-family cross-track differences, holds the learner and selection budget fixed.

Kernel bandwidth uses fitting-fold distances only. Neural fitting uses fixed epochs rather than test-informed early stopping. The frozen Geneformer V1-10M checkpoint SHA256 is a5e33a757431643b3697de7ef6127950cdc49e06e58d4266b3a3ab191b683f14. Only its 256-dimensional gene-token vectors and a missing-token indicator are used. No attention pass, cell embedding or response fine-tuning is performed. This baseline supplies a portable external prior; it is not a claim about native Geneformer inference. Token coverage and unknown-gene eligibility are recorded in the release evidence.

All entries below are mean outer-test Spearman correlations. Methods are not selected by these test values. Candidate counts and outer splits are identical within a task. The source-derived JSON tables also retain standard deviations, candidate-conditional intervals, secondary AUROC/AP counts and training-only selected-family frequencies.

#### Full-profile unseen comparators

| Task | Ridge | KRR | Boosting | MLP | Graph | Inner-selected |
|---|---|---|---|---|---|---|
| CD4 | -0.001 | 0.059 | 0.028 | 0.019 | 0.011 | 0.057 |
| FrangiehIFN | 0.058 | -0.017 | -0.037 | 0.046 | 0.048 | 0.017 |
| FrangiehCoc | 0.003 | 0.120 | 0.040 | 0.041 | 0.008 | 0.047 |
| Schmidt | -0.069 | -0.056 | -0.051 | -0.057 | -0.012 | -0.062 |
| Song_TCR | 0.005 | 0.019 | 0.014 | 0.008 | -0.004 | -0.001 |
| Papalexi_PDL1 | -0.004 | 0.007 | 0.194 | 0.018 | -0.051 | 0.028 |
| K562_Fitness | 0.178 | 0.383 | 0.335 | 0.274 | 0.290 | 0.383 |

#### Full-profile prior comparators

| Task | Ridge + prior | Boosting + prior | MLP + prior | Inner-selected |
|---|---|---|---|---|
| CD4 | 0.009 | 0.022 | 0.031 | 0.031 |
| FrangiehIFN | -0.009 | -0.009 | -0.039 | -0.036 |
| FrangiehCoc | 0.067 | 0.044 | 0.079 | 0.069 |
| Schmidt | 0.086 | -0.039 | -0.015 | -0.017 |
| Song_TCR | -0.008 | -0.018 | -0.006 | -0.009 |
| Papalexi_PDL1 | -0.143 | -0.128 | -0.021 | 0.006 |
| K562_Fitness | 0.212 | 0.374 | 0.313 | 0.374 |

#### Full-profile response-only comparators

| Task | Ridge | Boosting | MLP | Inner-selected |
|---|---|---|---|---|
| CD4 | 0.590 | 0.597 | 0.555 | 0.596 |
| FrangiehIFN | 0.175 | 0.136 | 0.144 | 0.170 |
| FrangiehCoc | 0.091 | 0.060 | 0.104 | 0.111 |
| Schmidt | 0.599 | 0.645 | 0.613 | 0.624 |
| Song_TCR | 0.010 | -0.012 | 0.025 | -0.009 |
| Papalexi_PDL1 | 0.354 | 0.319 | 0.328 | 0.261 |
| K562_Fitness | 0.373 | 0.445 | 0.359 | 0.445 |

#### Controlled full-profile availability

| Task | 0% | 10% | 25% | 50% | 100% |
|---|---|---|---|---|---|
| CD4 | -0.001 | 0.196 | 0.274 | 0.327 | 0.373 |
| FrangiehIFN | 0.058 | 0.074 | 0.076 | 0.173 | 0.187 |
| FrangiehCoc | 0.003 | 0.068 | 0.060 | 0.083 | 0.116 |
| Schmidt | -0.069 | 0.237 | 0.486 | 0.628 | 0.598 |
| Song_TCR | 0.005 | -0.003 | 0.017 | 0.032 | 0.005 |
| Papalexi_PDL1 | -0.004 | 0.130 | 0.310 | 0.370 | 0.358 |
| K562_Fitness | 0.178 | 0.276 | 0.295 | 0.303 | 0.307 |

#### Paired full-profile response-entry contrasts

| Task | Full minus zero correlation [95% interval] |
|---|---|
| CD4 | 0.374 [0.359, 0.389] |
| FrangiehIFN | 0.129 [0.013, 0.249] |
| FrangiehCoc | 0.114 [0.021, 0.213] |
| Schmidt | 0.667 [0.464, 0.814] |
| Song_TCR | -0.001 [-0.033, 0.035] |
| Papalexi_PDL1 | 0.361 [-0.145, 0.706] |
| K562_Fitness | 0.129 [0.117, 0.140] |

The independent zero-availability implementation reproduces full-profile B ridge predictions to relative/absolute tolerance 1e−7 in every split. Seven boundary tests verify zero-availability masking, held-out target isolation, training-only PCA, inductive graph invariance, the eigenvector condition, weighted bootstrap ranks and constant-score handling. These enforce software boundaries and do not prove that the biological measurements are noise-free.

#### Reference supplementary-table definitions

Supplementary Table 4. Multi-metric benchmark. Stored AUROC, AUPRC, precision@25, precision@100 and score Spearman summaries for the six configurations listed below. The separate Frangieh cross-condition and rank-target sensitivity analyses are not included in this table. These summaries do not establish equivalence or universal dominance.

Supplementary Table 7. Systematic-variation (axis-alignment) diagnostic. Axis alignment is the cosine between the transition axis and the average perturbation-effect axis, following Systema [9]. High alignment indicates overlap with a systematic response direction; it does not by itself establish that ranking is transition-specific or explain its accuracy. The single-condition Frangieh configuration has alignment 0.441, whereas control-to-IFN-gamma has alignment 0.218. Their different targets and information budgets prevent interpreting this contrast as an isolated test of alignment. Neither alignment nor measurement count has been validated as a sufficient predictor of held-out learning gain. The AUROC column reports continuous-target ridge, not the best method selected across specifications.

Supplementary Table 8. Cross-condition ranking configurations and map predictivity. Mean AUROC over ten held-out-perturbation splits, using source-condition features and a separate target-condition measurement. Header rho values summarize held-out source-to-target map predictivity. The three CD4+ pairs share one atlas, perturbations and donors and are within-atlas sensitivities. Frangieh co-culture has weak measured predictivity, and Schmidt shows no detected improvement over its covariance projection under the supplied axis; neither observation establishes zero information or population optimality. Cross-condition covariance projection and centroid contrast use training labels. The table retains continuous-target ridge; the rank-target sensitivity in Reference Table N10a changes the Frangieh comparison. The pert2state adaptation is evaluated under this benchmark's candidate, representation and target choices and does not establish a general ordering of the native methods. Differences between means are descriptive here; uncertainty for selected comparisons is reported separately.

#### Supplementary Note 11 | Independent biological endpoints and matched target construction

##### External functional endpoint

DepMap 24Q4 supplies pooled CRISPR-knockout gene effects independently of the source Replogle CRISPRi RNA signatures. The dataset is pinned to DOI 10.25452/figshare.plus.27993248.v1, under CC BY 4.0. The release metadata identify K562 as ACH-000551. The gene-effect matrix has SHA256 3d8f3ec6dbf2db7ff834b79b508622ec0b226f3518003fe96ecf5a4fcf167e3b. The endpoint is negative Chronos gene effect, so higher means stronger loss of pooled growth fitness. This is not a measure of therapeutic selectivity, clinical response or a new cellular context. Chronos is described by Dempster et al., Genome Biology 22, 343 (2021), DOI 10.1186/s13059-021-02540-7.

Named intervention/readout overlap, finite gene effect and unique-gene grouping define 7,354 eligible candidates without model or outcome threshold selection. Repeated transcriptomic records are averaged per gene before outer splitting. The supplementary interface records source input hashes, the K562 model identifier and exact outer candidate indices.

##### Same-input proxy versus fitness

A fixed random partition assigns 60% of readout genes to source features and the remaining 40% to construction of the transcriptional proxy. PC1 uses anchor-intervention rows of the label-gene matrix only; orientation uses anchor skew. Source inputs and candidate splits are identical for both endpoints. Nested ridge selection crosses three penalties and raw/rank training targets. Table N11a retains the covariance and fixed-penalty references as well as the selected estimator. Intervals are conditional candidate-cluster bootstrap intervals, not a paired causal comparison of assay accuracy.

| Endpoint | Inner-selected ridge | Fixed ridge | Covariance score |
|---|---|---|---|
| RNA PC1 proxy | 0.959 [0.956, 0.962] | 0.959 [0.956, 0.962] | 0.928 [0.922, 0.933] |
| Independent growth fitness | 0.445 [0.425, 0.466] | 0.444 [0.424, 0.465] | 0.403 [0.381, 0.424] |

The comparison is exploratory and was specified after initial six-task results, before any paired-target fits. It does not imply that measurement noise, assay modality or target complexity are controlled between endpoints. It demonstrates that a source-proxy score cannot be interpreted as a functional ranking score. Neither endpoint was chosen by comparing held-out candidate performance.

##### Surface-protein screen

Papalexi et al., Nature Genetics 53, 322–331 (2021), DOI 10.1038/s41588-021-00778-2, GSE153056, provide RNA, guide and ADT data. Replicate-1 RNA signatures supply source inputs.

The primary destination is lower surface PD-L1 in replicate 2, defined as negative mean log1p ADT abundance relative to non-targeting controls. Direct CD274 perturbation and its RNA readout are excluded. At least 20 cells per required arm leaves 24 gene interventions. Guide count is not gene-intervention count. Barcode de-duplication and guide identities were checked against the harmonized RNA matrix.

All primary replicate-2 predictions are retained. Without refitting, the same predictions are scored against replicate 3 and against within-cell log1p PD-L1 minus the mean log1p of the four measured ADTs in replicate 2. The latter is a normalization sensitivity, not a replacement primary endpoint. The outcome correlations with primary replicate 2 are 0.617 and 0.768, respectively. Replicates are technical; no biological replication is inferred.

| Protein endpoint | Anchor-profile ridge | Response-only ridge | Combined full-response ridge |
|---|---|---|---|
| Primary replicate 2 | -0.004 [-0.218, 0.215] | 0.354 [-0.086, 0.643] | 0.358 [-0.086, 0.646] |
| Secondary replicate 3 | -0.104 [-0.319, 0.148] | 0.525 [0.219, 0.718] | 0.535 [0.206, 0.729] |
| Secondary ADT log-ratio | -0.044 [-0.271, 0.189] | 0.362 [-0.061, 0.636] | 0.359 [-0.069, 0.633] |

The primary Spearman intervals are wide and include zero for the response-informed estimates. A positive secondary AUROC contrast does not resolve this uncertainty. Missing class-eligible splits and valid bootstrap counts are reported in the retained metric summaries.

#### Fixed activation markers

Song et al., Nature Cell Biology 27, 493–504 (2025), DOI 10.1038/s41556-025-01626-9, GSE249595, provide untreated and activated T-cell perturbation signatures. The destination is the mean activated CD69 and IL2RA control-referenced log-normalized expression. Both endpoint readouts and direct interventions are excluded from source features and candidates, leaving 1,883 genes. The source cache requires at least ten cells in each arm. The marker endpoint was specified before this model evaluation, but the cohort had already been analyzed with an unsupervised axis and is not a prospectively untouched validation cohort. The weak primary contrast is retained. It does not prove absence of useful information or establish a noise ceiling.

#### Functional shortlist consequences

The analysis specification was frozen before these additional fits, after the preceding ridge endpoint summaries were known. It is exploratory, not prospectively preregistered. The same 7,354 candidate genes, anchor/fitting/test splits and source-feature/label-gene partition were retained. For each endpoint, five families (ridge, kernel ridge, histogram gradient boosting, MLP and inductive GraphSAGE) each received six settings and three-fold inner Spearman selection. Family selection also used inner scores only. Functional test outcomes were used solely for final evaluation. The primary shortlist was the top 5%; 1% and 10% were specified secondary cutoffs.

Functional utility is mean within-test midrank percentile of negative Chronos effect. Secondary measures are precision for the functional top decile, mean functional effect standardized by fitting genes only, and normalized oracle-minus-policy percentile regret relative to random reference 0.5. The oracle is an evaluation ceiling, not a deployable comparator. A paired gene-cluster bootstrap resamples candidate genes with shared multiplicities across repeated test appearances, then reconstructs rankings and shortlists in each resample. The 1,000-draw intervals condition on the fitted models and do not include retraining uncertainty. They are exploratory and unadjusted.

| Shortlist | Proxy-selected utility | Functional-selected utility | Functional – proxy [95% interval] |
|---|---|---|---|
| 5% | 0.841 | 0.839 | -0.001 [-0.021, 0.023] |
| 1% | 0.918 | 0.890 | -0.028 [-0.049, -0.005] |
| 10% | 0.778 | 0.811 | 0.033 [0.016, 0.050] |

At the primary cutoff, top-functional-decile precision is 0.501 for proxy selection and 0.501 for functional selection. The data do not establish a practical shortlist penalty from proxy selection in K562. Absence of a detected difference does not establish equivalence. Growth suppression is not therapeutic selectivity, and these repeated partitions are not independent biological replications. The result prevents interpreting a difference in endpoint predictability as proof of worse functional decisions. Candidate-level predictions, all ten splits, selection records and all specified utility summaries are retained locally.

#### Melanoma co-culture functional validation

A second biological context uses the published Frangieh et al. pooled CRISPR viability screen (Nature Genetics 53, 332–341, 2021; DOI 10.1038/s41588-021-00779-1). The functional endpoint was fixed before fitting as the negative mean MAGeCK gene-level log2 fold change across all three reported TIL:cancer-cell ratios (1:1, 2:1 and 4:1), each contrasted against IFNγ-treated day-17 cells. This controls the IFNγ exposure shared by co-culture conditions. Supplementary Tables 2g, 2k and 2o provide the contrasts; no significance threshold selects candidates. Their intersection with the existing untreated RNA source signatures retains 226 genes. The source cohort has already been analyzed; this is a separate functional assay and a second context relative to K562, not an independent source-study replication. The patient-derived 2686 cancer/TIL system and assay relationship are documented in the original methods.

The same five families, inner validation, ten disjoint anchor/fitting/test splits, readout-gene partition and predefined shortlist metrics were used. The mean test correlations of inner-selected models are -0.057 for the functional endpoint and 0.913 for the RNA proxy. At the primary top-5% cutoff, mean functional percentile is 0.518 for proxy selection and 0.440 for functional selection. The paired functional-minus-proxy difference is −0.078 [−0.172, 0.063]. The small test shortlist (five genes per split), outcome noise and overlapping partitions limit inference. This second context does not establish functional decision harm from proxy selection;

training directly on the functional endpoint does not guarantee improved shortlist utility. All ratios, genes, splits and negative results are retained.

The secondary K562 cutoffs reverse the ordering: functional-minus-proxy utility is −0.028 [−0.049, −0.005] at 1% and 0.033 [0.016, 0.050] at 10%. Melanoma contrasts are −0.241 [−0.335, −0.013] at 1% and −0.022 [−0.118, 0.081] at 10%. These intervals are exploratory and unadjusted; the melanoma 1% shortlist contains only one candidate per split and is particularly unstable. A functional penalty from proxy selection is supported only for the secondary K562 10% comparison under this conditional analysis, and cannot be generalized across cutoffs, studies or retraining environments.

#### Native unseen-perturbation functional comparisons

The raw Replogle–Weissman K562 essential-gene assay from Zenodo record 13350497 contains 310,385 cells and 8,563 measured genes (file MD5 d8cba17576d1a8afc0f7d71b79cad0f7). Eligibility requires at least 20 cells per single perturbation, a measured gene, pinned GO annotation, scGPT vocabulary coverage and finite matching K562 fitness. This retains 1,728 candidates without outcome-based significance or performance filtering. This essential-gene assay subset differs from the preceding 7,354-gene GWPS source comparison.

Three outer splits each contain 518 anchors, 518 fitting genes and 692 test genes. Source cells are capped at 32 actual cells per anchor intervention and 128 controls, giving 16,608–16,627 cells. The top 2,048 highly variable readouts are selected using these anchor/control cells only and combined with measured candidate genes, giving 3,527–3,532 readouts. No fitting- or test-intervention cells enter response-model training, feature selection or validation. Fifteen percent of anchor genes are reserved for RNA validation. Fitting outcomes alone train the functional head; held-out functional outcomes are used only for evaluation.

Released cell-gears 0.1.2 uses hidden size 64, batch size 32, 20 epochs and its native validation top-DE-gene RNA-MSE checkpoint rule. scGPT 0.2.4 uses the released TransformerGenerator, official whole-human checkpoint (SHA256 6cb5d451ab5c4b33eb673adbe4fddc61d2389df1b89b7651a9fe2e557572b922), 12 layers, 512-dimensional embeddings and eight heads. Encoder, value encoder and transformer weights initialize the native response model. It is fine-tuned for 20 epochs with Adam at 0.0001, batch size 4, gradient norm clipping at 1 and bfloat16 computation; anchor-validation RNA MSE selects its checkpoint. The vocabulary-covered scGPT panel contains 3,412–3,428 readouts. Namespace loading avoids importing unrelated downstream task packages; released model computations and pretrained loading utilities are unchanged. Pretraining overlap is unverified.

GEARS predictions average 300 sampled control cells using the released prediction routine; scGPT predictions average 32 controls. Both predict fitting and test interventions as unseen. All functional heads use the same six-setting ridge raw/rank search and three-fold fitting-only Spearman validation, including fold-fitted scaling/PCA. Anchor-profile ridge and kernel ridge use the same observed anchor signatures; measured-response ridge is a response-informed positive control. Native response-model architectures, representation coverage and inference budgets are not identical, and fixed recipes are not exhaustive hyperparameter searches.

| **Method** | **Functional Spearman [95% interval]** | **Top-5% functional-percentile utility [95% interval]** |
|---|---|---|
| Native GEARS + ridge | 0.203 [0.155, 0.246] | 0.716 [0.659, 0.763] |
| Native scGPT + ridge | 0.264 [0.212, 0.314] | 0.791 [0.729, 0.833] |
| Anchor-profile ridge | 0.304 [0.249, 0.351] | 0.715 [0.652, 0.774] |
| Anchor-profile kernel ridge | 0.325 [0.276, 0.371] | 0.743 [0.680, 0.795] |
| Measured-response ridge | 0.490 [0.441, 0.529] | 0.769 [0.711, 0.817] |

Paired gene-cluster resampling uses 1,000 shared draws across repeated test appearances and reconstructs shortlists using candidate multiplicities. Intervals condition on fitted models and are exploratory and unadjusted. Relative to anchor ridge, GEARS correlation differs by −0.100 [−0.157, −0.038], while its primary shortlist utility differs by 0.001 [−0.073, 0.070]. scGPT correlation differs by −0.040 [−0.085, 0.003], while primary shortlist utility improves by 0.076 [0.001, 0.132]. These results do not justify a universal ordering of native and simpler models. The small conditional interval above zero for scGPT shortlist utility should not be treated as donor-level replication or multiplicity-adjusted evidence.

Split-specific working directories isolate ontology graph caches. Graph endpoints were checked against each model node set. All final three GEARS and three scGPT fits completed 20 epochs. Retained predictions, split indices, validation records, environment/source hashes and graph audits support the comparison; exact historical neural reproduction remains a separate unresolved issue.

#### Supplementary Note 12 | Theory calibration and conclusion stability

##### Conditional theory and training-only predictions

Supplementary Note 2 establishes a jointly Gaussian population result for a specified covariance projection. Let v = cov(X,y) and Sigma = cov(X). Projection and optimal linear directions coincide when v is an eigenvector of Sigma. This is not a theorem that one measurement cannot support learning or that two measurements suffice.

We tested incremental explanatory value by deriving all diagnostics from fitting candidates only. The eigenvector misalignment is one minus squared cosine between v and Sigma v. Estimated Gaussian correlations for covariance and whitened linear scores are converted to predicted Spearman correlations by the Gaussian copula identity, 6/pi arcsin(rho/2). Observed gains compare a fixed ridge penalty of ten with the covariance score on outer-test candidates. No held-out gain is used to construct a diagnostic.

The simulation crosses training sizes 60, 200 and 1,000; feature dimensions 5 and 20; four angles; covariance anisotropies 1, 10 and 100; three signal-to-noise ratios; Gaussian and Student-t(3) populations; and ten seeds, giving 4,320 configurations. Independent test samples contain 4,000 observations. The Gaussian condition is not assumed to hold for the heavy-tailed sensitivity.

| Population | Correlation of predicted and observed gain | Mean absolute prediction error |
|---|---|---|
| Gaussian | 0.846 | 0.041 |
| Student_t3 | 0.815 | 0.055 |

##### Real-study calibration includes the negative result

Frangieh configurations are grouped together for study-level evaluation. Training-only predictions do not establish useful real-study calibration: the correlation across six study groups is −0.143, with a study bootstrap interval spanning −1 to 1. Leave-study-out gain prediction has mean absolute error 0.093. Six groups provide very limited calibration power; this result supplies no validated real-data task-difficulty predictor.

| Task | Eigenvector misalignment | Predicted Gaussian gain | Observed held-out gain |
|---|---|---|---|
| CD4 | 0.106 | 0.135 | 0.085 |
| FrangiehCoc | 0.365 | 0.156 | 0.023 |
| FrangiehIFN | 0.315 | 0.164 | 0.047 |
| K562_Fitness | 0.056 | 0.072 | 0.038 |
| Papalexi_PDL1 | 0.036 | 0.063 | -0.042 |
| Schmidt | 0.176 | 0.335 | -0.102 |
| Song_TCR | 0.428 | 0.077 | -0.007 |

#### Computational stability is evaluated on conclusions

Reference, newer-environment, frozen-environment and reference-layout computations retain the same candidate labels and retained-row identities in all seven reference configurations. We compare mean Ridge-minus-CellOT, Ridge-minus-PAIRING and kernel-ridge-minus-Ridge AUROC differences. None changes sign across the four computations. This is descriptive ordering stability for these contrasts, not a claim that every pairwise ordering is unchanged or that small differences establish meaningful superiority. The corrected-layout mean neural discrepancy can still reach 0.0138. Exact reference neural score reproduction remains unresolved.

Target and objective changes answer different questions. Rank-target fitting changes the Frangieh mean comparison, while actual-target PC2 retraining changes conclusions relative to cross-target rescoring. Both the weak and positive target-specific results are retained in Supplementary Note 7. Main Figure 4 places numerical and objective sensitivity together; these checks do not establish equivalence or universal robustness.

#### Extended Data figures

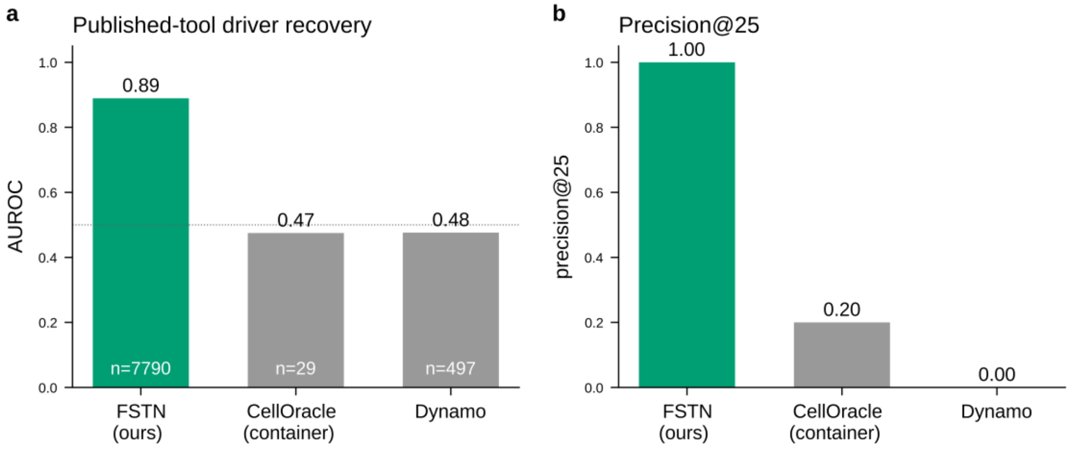


*Extended Data Fig. 1 | Published-tool comparison (CellOracle/Dynamo) on non-commensurable candidate sets. a, driver-recovery AUROC: FSTN 0.89 over 7,790 ranked candidates versus CellOracle 0.47 (29 candidates) and Dynamo 0.48 (497), both at chance (dotted line). b, precision@25 over the same rankings (1.00, 0.20 and 0.00). The candidate sets differ by two orders of magnitude and are set by each tool's own output, so these panels show that the published tools are not commensurable with the shared-space protocol rather than that they are weaker on a like-for-like task.*

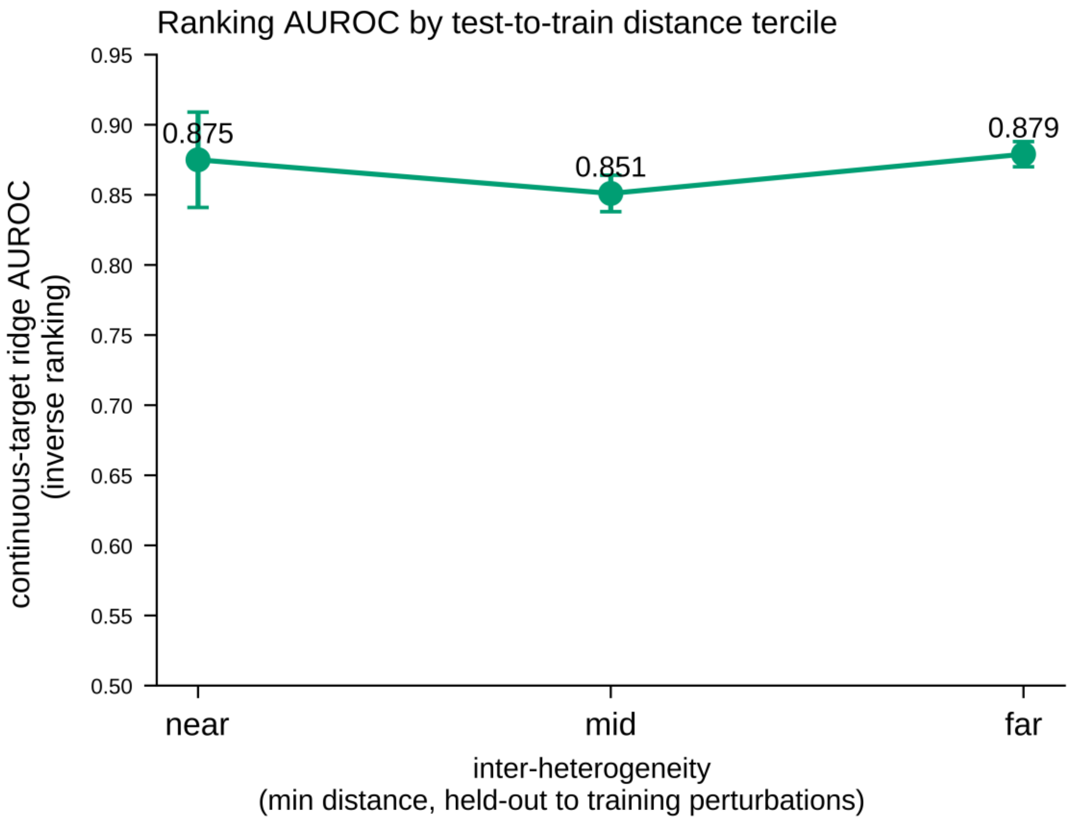


*Extended Data Fig. 2 | Inverse driver ranking across signature-distance terciles (CD4+ near/mid/far 0.875/0.851/0.879).*

**a**

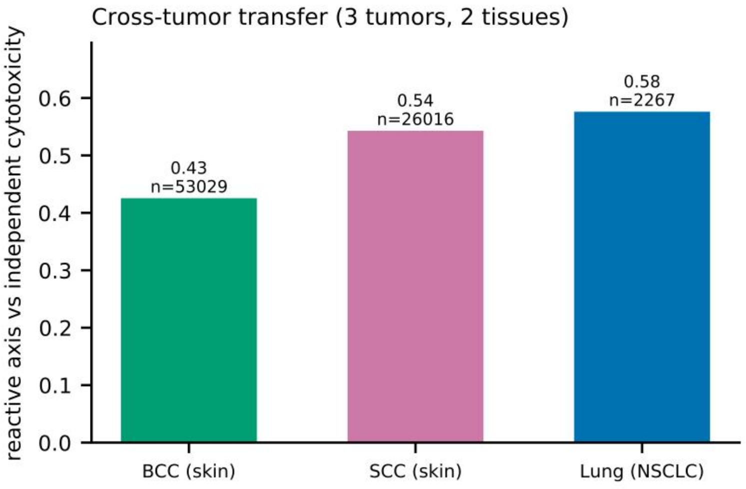


**b**

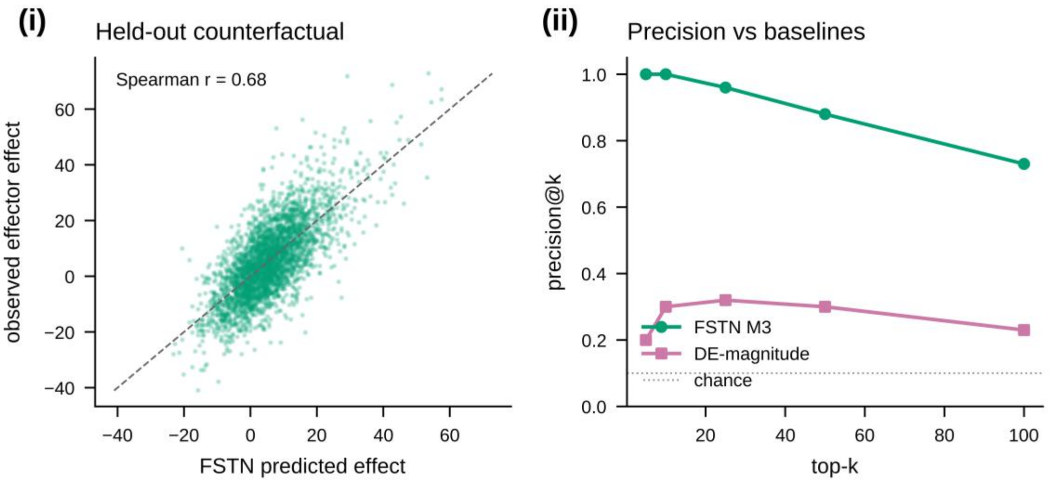


*Extended Data Fig. 3 | Transfer of the reactive program beyond the primary system. a, cross-tumor transfer to BCC, SCC and NSCLC (r = +0.43 to +0.58). b, held-out cytotoxic-lineage comparisons: (i) held-out counterfactual agreement (Spearman r = 0.68) and (ii) precision@k against baselines.*

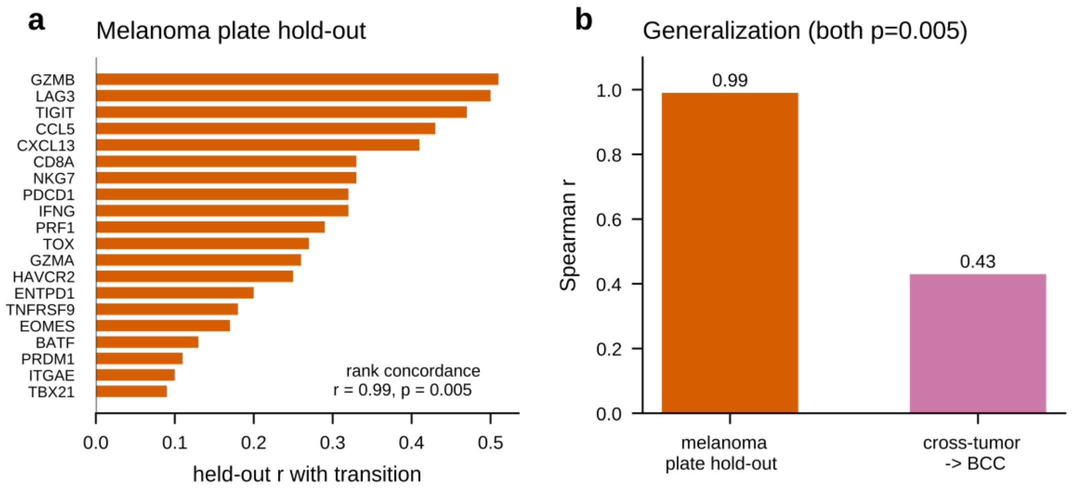


*Extended Data Fig. 4 | Generalization of the reactive-regulator ranking within the primary system. a, melanoma plate hold-out: per-gene held-out correlation with the transition for the top reactive regulators, ranked on one disjoint plate set and scored on the other (rank concordance r = 0.99, shuffled-label p = 0.005). b, the same concordance summarised against one cross-tumor instance (melanoma plate hold-out 0.99; melanoma to basal cell carcinoma 0.43; both shuffled-label p = 0.005). The full three-tumor transfer is Extended Data Fig. 3a.*

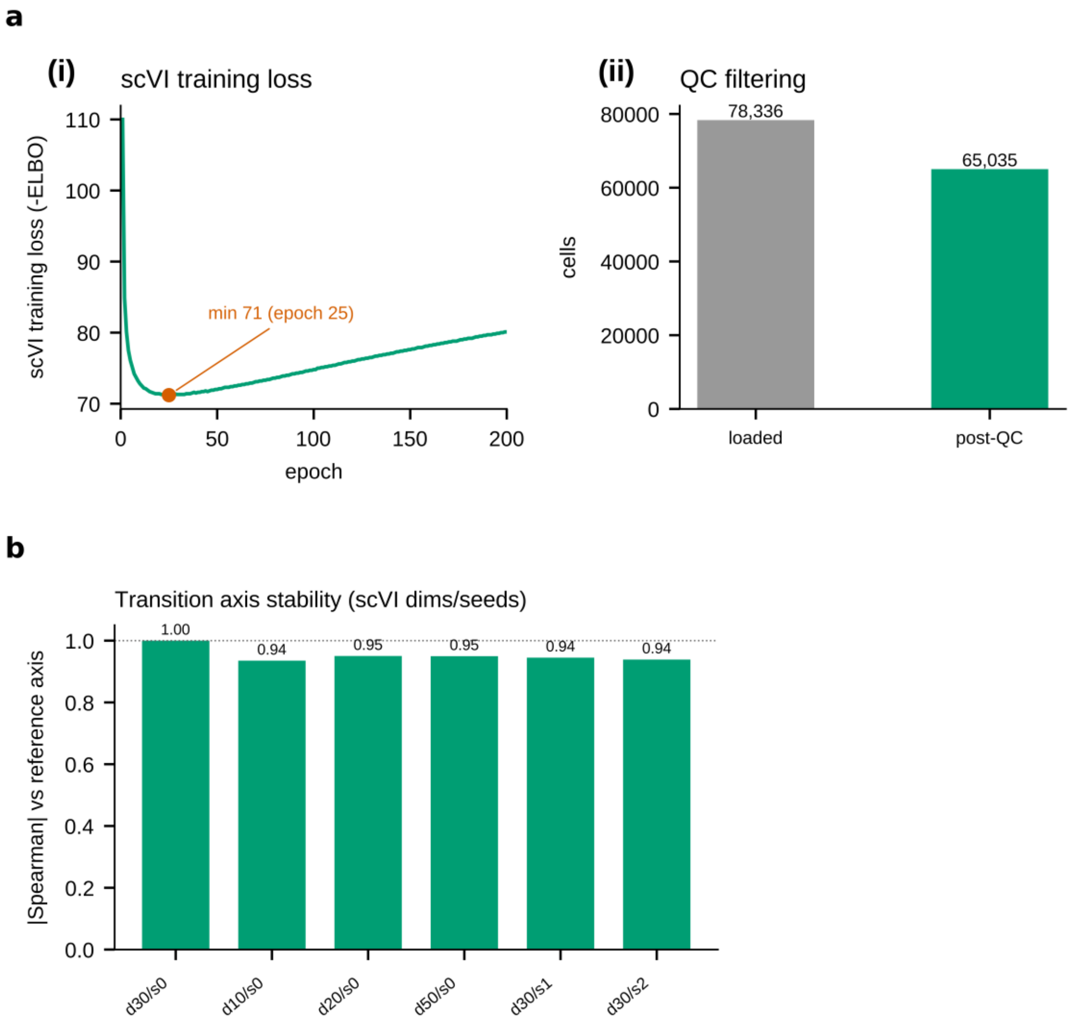


*Extended Data Fig. 5 | Embedding quality control and transition-axis stability. a, (i) historical scVI training loss (negative ELBO) over the full 200-epoch budget: the loss falls steeply, reaches its minimum of 71 at epoch 25 and then rises slowly to 80 by epoch 200, so the budget was not set by convergence of this curve; the embedding is instead qualified by the stability in b. (ii) QC filtering (78,336 loaded to 65,035 post-QC cells). b, integer-count latent-projection sensitivity across dimensions and seeds; this projection is distinct from the expression-defined melanoma coordinate (absolute Spearman correlations 0.9352 to 0.9503 against the reference axis; Supplementary Note 9).*

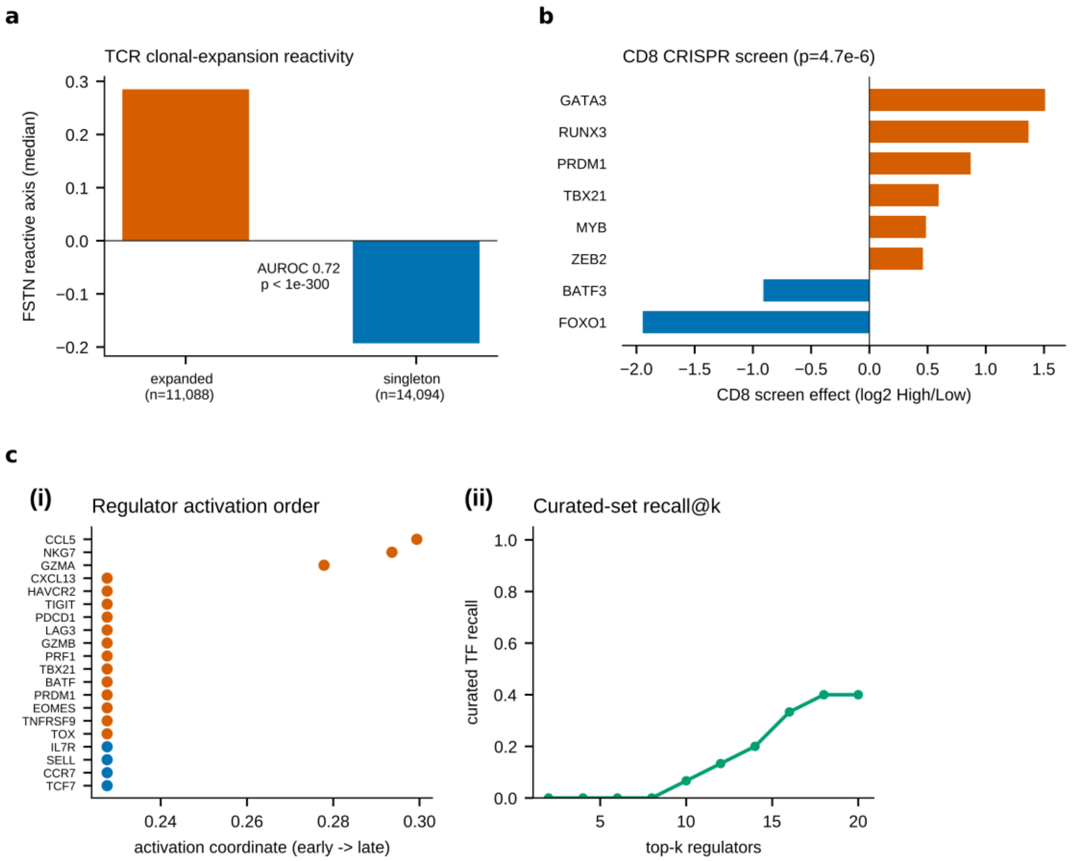


*Extended Data Fig. 6 | Orthogonal validation of the reactivity axis. a, TCR clonal expansion separates expanded clones from singletons (AUROC 0.72). b, CD8+ CRISPR screen validation of the recovered transcription factors (Mann-Whitney p = 4.7e-6). c, (i) regulator-expression threshold crossings along the coordinate: crossings cluster within a narrow range, with CCL5, NKG7 and GZMA at larger coordinates. This is not a temporal activation measurement. (ii) recall of the curated tumor-reactivity TF set over a standard human TF catalog: 10 of 15 curated factors fall within the top 50 (recall@50 = 0.67), with TOX, EOMES and ID2 the three top-ranked of 1,731 factors.*

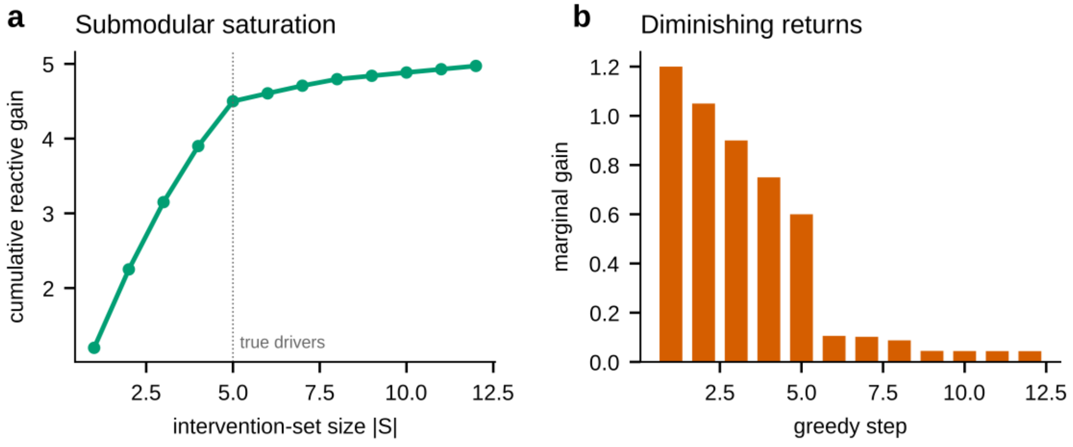


*Extended Data Fig. 7 | M3 counterfactual solver behaviour. a, synthetic saturation example: cumulative reactive gain against intervention-set size, which flattens at the five true drivers (dotted line). b, diminishing returns: the marginal gain of each greedy step falls sharply after the fifth edit.*

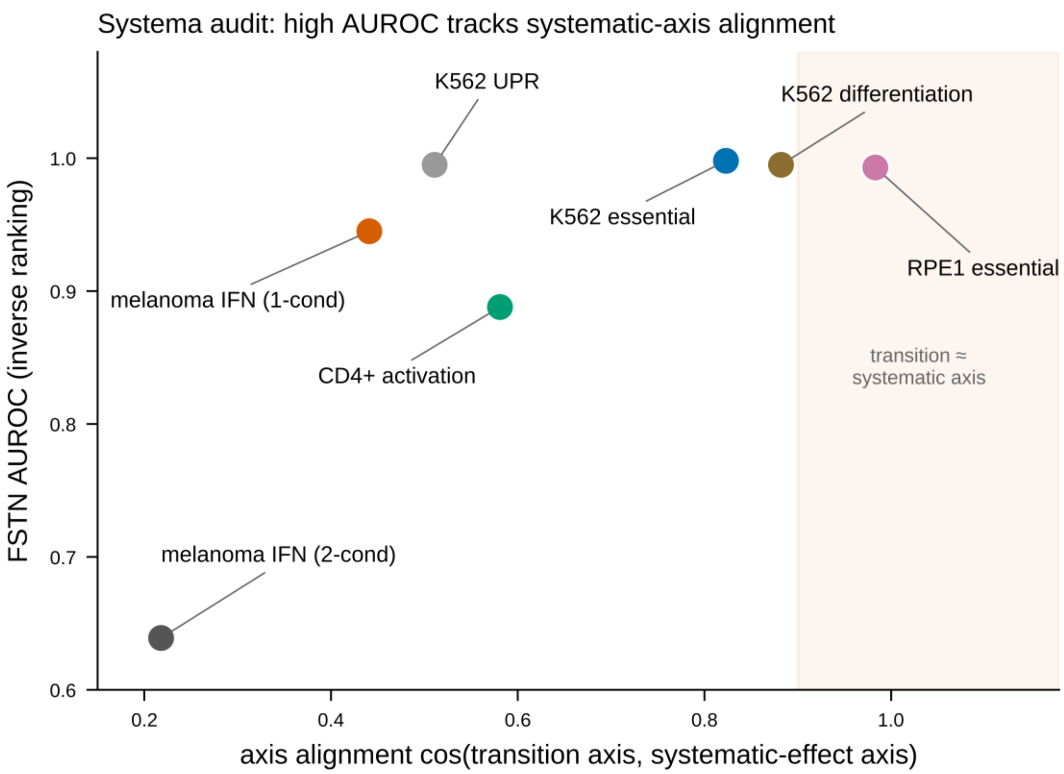


*Extended Data Fig. 8 | Systematic-variation (axis-alignment) diagnostic across the seven system columns (after Systema).*

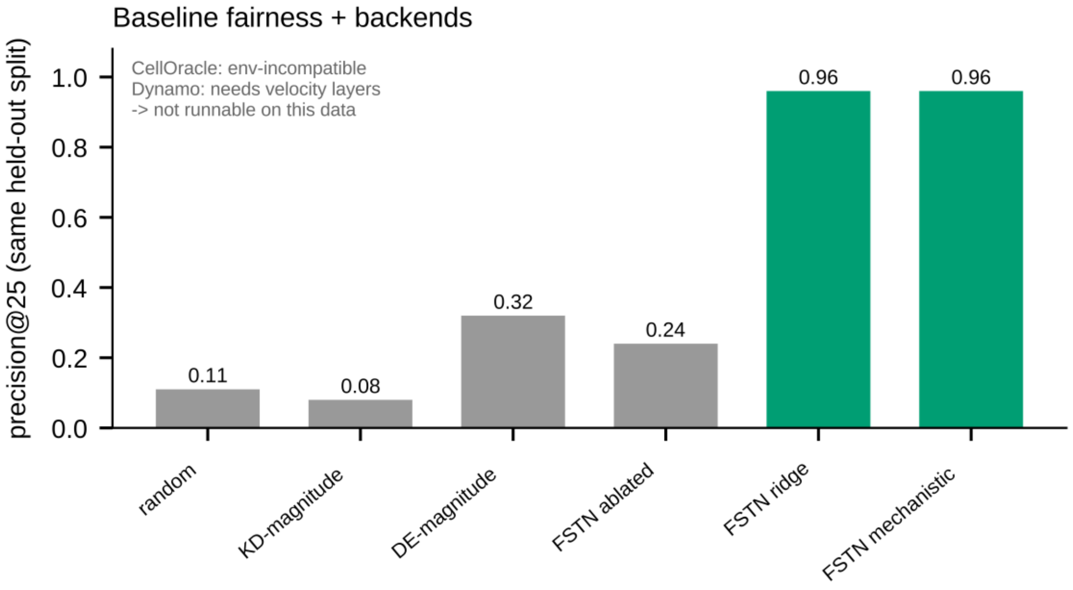


*Extended Data Fig. 9 | Baseline fairness: FSTN ridge and mechanistic backends versus computational baselines.*

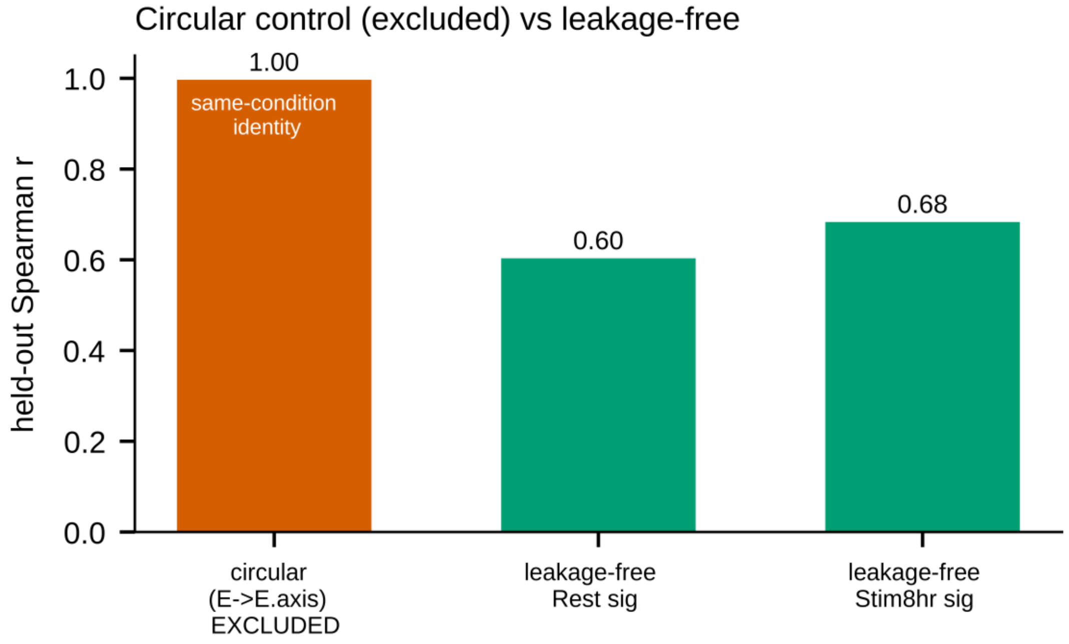


*Extended Data Fig. 10 | Leakage control: the excluded circular same-condition fit (Spearman 0.997).*

#### Supplementary Tables — datasets, targets, software

#### InverseBench - Supplementary Tables

Information contracts determine what inverse perturbation benchmarks measure

##### Supplementary Table 1 | Dataset manifest and accessions

| role | source | platform | access | n | used_in | notes |
|---|---|---|---|---|---|---|
| Melanoma TILs (primary demonstration) | GSE123139 | MARS-seq (UMI counts) | open | 65,035 cells post-QC | Supplementary Figures 1, 2, 4; Extended Data Fig. 5 and Extended Data Fig. 6 | reactivity via dysfunction program |
| CD4+ T-cell activation (primary benchmark system) | GSE314342 | genome-scale CRISPRi Perturb-seq | open | 11,526 perturbed genes; 33,983 profiles | Fig. 2a; Fig. 4; Table 1 | zscore layer: 33,983 perturbation-by-condition profiles across 11,526 genes in three conditions (resting, 8 h, 48 h); 8 h features, 48 h labels |
| Melanoma IFN response (Frangieh et al.) | ref. [36] | CRISPR-KO Perturb-seq | open | 246 perturbations | Reference Table N10a; Supplementary Notes 1 and 10 | analysed as both a two-condition and a single-condition system |
| K562 essential-gene screen (Replogle et al.) | ref. [27] | genome-wide CRISPRi Perturb-seq | open | 11,258 perturbations | Reference Table N10a; Supplementary Notes 1 and 10 | |
| RPE1 essential-gene screen (Replogle et al.) | ref. [27] | genome-wide CRISPRi Perturb-seq | open | 2,679 perturbations | Reference Table N10a; Supplementary Notes 1 and 10 | |

| role | source | platform | access | n | used_in | notes |
|---|---|---|---|---|---|---|
| K562 erythroid differentiation (Norman et al.) | GSE133344 | CRISPRa Perturb-seq | open | 236 perturbations | Reference Table N10a; Supplementary Notes 1 and 10 | second independent transition |
| K562 unfolded-protein response (Adamson et al.) | ref. [37] | CRISPRi Perturb-seq | open | 107 perturbations | Reference Table N10a; Supplementary Notes 1 and 10 | |
| Cross-tumor generalization (BCC/SCC) | GSE123813 | 10x scRNA-seq + TCR | open | 53,029 BCC + 26,016 SCC cells | Reference Table N10a; Supplementary Notes 1 and 10 | Yost et al. |
| Cross-tumor generalization (NSCLC) | GSE218262 | 10x scRNA-seq | open | 2,267 cells | Reference Table N10a; Supplementary Notes 1 and 10 | processed with the same QC |
| CD8+ CRISPR screen validation | GSE241933 | pooled CRISPR + scRNA-seq | open | transcription-factor screen | Reference Table N10a; Supplementary Notes 1 and 10 | effector/memory axis; per-gene phenotype = mean log2 fold change of a gene's guides |
| Dendritic-cell LPS activation (negative control) | GSE90063 | Perturb-seq | open | - | Reference Table N10a; Supplementary Notes 1 and 10 | Dixit et al.; honest-null control |
| neoTCR reactivity labels (reference only) | dbGaP phs002748 | 10x + neoTCR | CONTROLLED | - | Reference Table N10a; Supplementary Notes 1 and 10 | not used in any reported analysis; supplementary tables S7/S8 of the source are open |

| role | source | platform | access | n | used_in | notes |
|---|---|---|---|---|---|---|
| Surface PD-L1 destination | GSE153056 | ECCITE-seq RNA and ADT | open | 24 eligible gene interventions | Fig. 2d; Fig. 3c; Table 1 | Replicate-1 source RNA; replicate-2 primary protein; technical replicate-3 sensitivity |
| Fixed T-cell marker destination | GSE249595 | T-cell Perturb-seq | open | 1,883 eligible gene interventions | Fig. 2g; Table 1 | CD69/IL2RA target fixed before model fits; marker genes excluded from features; at least ten cells per arm |
| Independent functional destination | 10.25452/figshare.plus.27993248.v1 | DepMap 24Q4 pooled CRISPR knockout Chronos | CC BY 4.0 | 7,354 eligible K562 genes | Fig. 2c; Fig. 3a,b,d; Table 1 | K562 ACH-000551; paired with Replogle CRISPRi RNA; independent assay rather than new cell context |
| Melanoma/TIL killing endpoint | Frangieh et al. (2021); ST2g/k/o | Pooled CRISPR-Cas9 viability | Published workbook | 226 matched genes | Fig. 3d; Note 11 | Mean of three TIL ratios vs IFNγ control; shared RNA cohort, separate viability assay |

#### Supplementary Table 2 | Curated tumor-reactivity transcription-factor set

| curated_TF (frozen pre-scoring) |
|---|
| **TOX** |
| **TCF7** |
| **PRDM1** |
| **TBX21** |
| **EOMES** |
| **ID2** |
| **BATF** |
| **IRF4** |
| **NR4A1** |
| **NR4A2** |
| **NR4A3** |
| **GATA3** |
| **RUNX3** |
| **ZEB2** |
| **TOX2** |

#### Supplementary Table 3 | Full M2 regulator ranking (Spearman r with the transition coordinate)

| gene | spearman_r_with_transition | pole |
|---|---|---|
| GZMB | 0.5030 | reactive |
| LAG3 | 0.4882 | reactive |
| TIGIT | 0.4598 | reactive |
| CCL5 | 0.4451 | reactive |
| CXCL13 | 0.4108 | reactive |
| NKG7 | 0.3506 | reactive |
| CD8A | 0.3378 | reactive |
| IFNG | 0.3142 | reactive |
| PDCD1 | 0.3120 | reactive |
| PRF1 | 0.2961 | reactive |
| GZMA | 0.2724 | reactive |
| TOX | 0.2634 | reactive |
| HAVCR2 | 0.2225 | reactive |
| ENTPD1 | 0.1954 | reactive |
| EOMES | 0.1786 | reactive |
| ID2 | 0.1759 | reactive |
| TNFRSF9 | 0.1737 | reactive |
| GZMK | 0.1500 | reactive |
| TOX2 | 0.1486 | reactive |

| gene | spearman_r_with_transition | pole |
|---|---|---|
| GNLY | 0.1279 | reactive |
| BATF | 0.1217 | reactive |
| PRDM1 | 0.1145 | reactive |
| GATA3 | 0.0992 | reactive |
| ITGAE | 0.0939 | reactive |
| TBX21 | 0.0846 | reactive |
| RUNX3 | 0.0658 | reactive |
| NR4A2 | 0.0482 | reactive |
| KLRG1 | 0.0419 | reactive |
| IRF4 | -0.0255 | naive/memory |
| NR4A3 | -0.0287 | naive/memory |
| TCF7 | -0.0578 | naive/memory |
| CCR7 | -0.0839 | naive/memory |
| NR4A1 | -0.0884 | naive/memory |
| IL7R | -0.0932 | naive/memory |
| ZEB2 | -0.1570 | naive/memory |
| SELL | -0.1651 | naive/memory |

#### Supplementary Table 4 | Multi-metric benchmark (AUROC, AUPRC, precision@25, precision@100, Spearman)

Stored AUROC, AUPRC, precision@25, precision@100 and score Spearman summaries for the six configurations listed below. The separate Frangieh cross-condition and rank-target sensitivity analyses are not included in this table. These summaries do not establish equivalence or universal dominance.

| System | Method | AUROC | AUPRC | prec@25 | prec@100 | Spearman |
|---|---|---|---|---|---|---|
| CD4+ activation | FSTN (linear) | 0.888 | 0.57 | 0.94 | 0.81 | 0.67 |
| CD4+ activation | Additive (no learning) | 0.834 | 0.42 | 0.78 | 0.60 | 0.56 |
| CD4+ activation | CellOT | 0.882 | 0.56 | 0.93 | 0.79 | 0.65 |
| CD4+ activation | PAIRING | 0.845 | 0.39 | 0.57 | 0.55 | 0.59 |
| melanoma IFN (Frangieh) | FSTN (linear) | 0.945 | 0.76 | 0.33 | 0.09 | 0.88 |
| melanoma IFN (Frangieh) | Additive (no learning) | 0.953 | 0.78 | 0.34 | 0.09 | 0.90 |
| melanoma IFN (Frangieh) | CellOT | 0.902 | 0.61 | 0.31 | 0.09 | 0.70 |

| System | Method | AUROC | AUPRC | prec@25 | prec@100 | Spearman |
|---|---|---|---|---|---|---|
| melanoma IFN (Frangieh) | PAIRING | 0.817 | 0.47 | 0.26 | 0.09 | 0.62 |
| K562 essential (Replogle) | FSTN (linear) | 0.998 | 0.99 | 1.00 | 1.00 | 0.96 |
| K562 essential (Replogle) | Additive (no learning) | 0.998 | 0.99 | 1.00 | 1.00 | 0.96 |
| K562 essential (Replogle) | CellOT | 0.981 | 0.90 | 1.00 | 0.99 | 0.80 |
| K562 essential (Replogle) | PAIRING | 0.970 | 0.73 | 0.78 | 0.81 | 0.58 |
| RPE1 essential (Replogle) | FSTN (linear) | 0.993 | 0.94 | 1.00 | 0.89 | 1.00 |
| RPE1 essential (Replogle) | Additive (no learning) | 0.993 | 0.94 | 1.00 | 0.89 | 1.00 |
| RPE1 essential (Replogle) | CellOT | 0.959 | 0.77 | 0.93 | 0.71 | 0.88 |
| RPE1 essential (Replogle) | PAIRING | 0.918 | 0.50 | 0.61 | 0.51 | 0.88 |
| K562 differentiation (Norman) | FSTN (linear) | 0.995 | 0.97 | 0.39 | 0.10 | 0.96 |
| K562 differentiation (Norman) | Additive (no learning) | 0.996 | 0.97 | 0.39 | 0.10 | 0.96 |
| K562 differentiation (Norman) | CellOT | 0.924 | 0.69 | 0.34 | 0.10 | 0.82 |
| K562 differentiation (Norman) | PAIRING | 0.724 | 0.33 | 0.22 | 0.10 | 0.49 |
| K562 UPR (Adamson) | FSTN (linear) | 0.995 | 0.97 | 0.16 | 0.10 | 0.92 |
| K562 UPR (Adamson) | Additive (no learning) | 1.000 | 1.00 | 0.16 | 0.10 | 0.98 |
| K562 UPR (Adamson) | CellOT | 0.955 | 0.80 | 0.16 | 0.10 | 0.79 |
| K562 UPR (Adamson) | PAIRING | 0.782 | 0.40 | 0.14 | 0.10 | 0.53 |

#### Supplementary Table 5 | Hypothesis-generating melanoma target list with CD4+ cross-support

| rank | gene | mel_axis_effect | tractability | priority | CD4_effect | CD4_percentile |
|---|---|---|---|---|---|---|
| 1 | **LAG3** | +0.488 | 1.00 | 0.488 | +14.3 | 78 |
| 2 | **TIGIT** | +0.460 | 1.00 | 0.460 | +0.1 | 35 |
| 3 | **CXCL13** | +0.411 | 1.00 | 0.411 | n/a | n/a |
| 4 | **IFNG** | +0.314 | 1.00 | 0.314 | +14.4 | 79 |
| 5 | **PDCD1** | +0.312 | 1.00 | 0.312 | +2.4 | 43 |
| 6 | **CTLA4** | +0.281 | 1.00 | 0.281 | -2.8 | 26 |
| 7 | **CCL5** | +0.445 | 0.60 | 0.267 | +7.9 | 61 |
| 8 | **TOX** | +0.263 | 1.00 | 0.263 | -5.0 | 21 |
| 9 | **HAVCR2** | +0.223 | 1.00 | 0.223 | +6.5 | 57 |
| 10 | **GZMB** | +0.503 | 0.40 | 0.201 | +12.7 | 75 |
| 11 | **ENTPD1** | +0.195 | 1.00 | 0.195 | +27.8 | 95 |
| 12 | **EOMES** | +0.179 | 1.00 | 0.179 | +9.1 | 65 |

#### Supplementary Table 6 | Software versions (frozen environment)

| package | version |
|---|---|
| scvi-tools | 1.1.3 |
| torch | 2.3.1+cpu |
| torchvision | 0.18.1+cpu |
| scanpy | 1.10.1 |
| anndata | 0.11.4 |
| jax | 0.4.30 |
| jaxlib | 0.4.30 |
| numpy | 1.26.4 |
| scipy | 1.15.3 |
| scikit-learn | 1.6.1 |
| pandas | 2.3.3 |
| matplotlib | 3.10.9 |
| dynamo-release | 1.5.3 |

#### Supplementary Table 7 | Systematic-variation (axis-alignment) diagnostic

Axis alignment is the cosine between the transition axis and the average perturbation-effect axis, following Systema [9]. High alignment indicates overlap with a systematic response direction; it does not by itself establish that ranking is transition-specific or explain its accuracy. The single-condition Frangieh configuration has alignment 0.441, whereas control-to-IFN-gamma has alignment 0.218. Their different targets and information budgets prevent interpreting this contrast as an isolated test of alignment. Neither alignment nor measurement count has been validated as a sufficient predictor of held-out learning gain. The AUROC column reports continuous-target ridge, not the best method selected across specifications.

| System | axis alignment | FSTN AUROC |
|---|---|---|
| RPE1 essential (Replogle) | 0.983 | 0.993 |
| K562 differentiation (Norman) | 0.882 | 0.995 |
| K562 essential (Replogle) | 0.823 | 0.998 |
| CD4+ activation | 0.581 | 0.888 |
| K562 UPR (Adamson) | 0.511 | 0.995 |
| melanoma IFN, single-condition (Frangieh) | 0.441 | 0.945 |
| melanoma IFN, two-condition (Frangieh) | 0.218 | 0.639 |

#### Supplementary Table 8 | Cross-condition ranking configurations and map predictivity

Mean AUROC over ten held-out-perturbation splits, using source-condition features and a separate target-condition measurement. Header rho values summarize held-out source-to-target map predictivity. The three CD4+ pairs share one atlas, perturbations and donors and are within-atlas sensitivities. Frangieh co-culture has weak measured predictivity, and Schmidt shows no detected improvement over its covariance projection under the supplied axis; neither observation establishes zero information or population optimality. Cross-condition covariance projection and centroid contrast use training labels. The table retains continuous-target ridge; the rank-target sensitivity in Reference Table N10a changes the Frangieh comparison. The pert2state adaptation is evaluated under this benchmark's candidate, representation and target choices and does not establish a general ordering of the native methods. Differences between means are descriptive here; uncertainty for selected comparisons is reported separately.

| **method** | **CD4+ 8h->48h (rho=0.67)** | **CD4+ rest->8h (rho=0.66)** | **CD4+ rest->48h (rho=0.56)** | **Frangieh IFNγ (rho=0.20)** | **Frangieh co-culture (rho=0.09)** | **Schmidt T-cell (rho=0.63)** |
|---|---|---|---|---|---|---|
| FSTN (linear) | 0.888 | 0.908 | 0.838 | 0.639 | 0.587 | 0.820 |
| Covariance projection (training-label supervised) | 0.834 | 0.780 | 0.733 | 0.575 | 0.604 | 0.828 |
| CellOT | 0.882 | 0.904 | 0.829 | 0.761 | 0.591 | 0.831 |
| PAIRING | 0.845 | 0.880 | 0.804 | 0.722 | 0.594 | 0.815 |
| scGen-style | 0.764 | 0.802 | 0.750 | 0.725 | 0.563 | 0.720 |
| Label-informed centroid contrast | 0.887 | 0.909 | 0.837 | 0.759 | 0.594 | 0.840 |
| deep MLP | 0.869 | 0.886 | 0.807 | 0.565 | 0.534 | 0.816 |
| kernel ridge | 0.893 | 0.911 | 0.848 | 0.615 | 0.556 | 0.811 |
| gradient boosting | 0.883 | 0.898 | 0.839 | 0.546 | 0.529 | 0.776 |
| pert2state (prior art) | 0.589 | 0.607 | 0.533 | 0.663 | 0.494 | 0.526 |
